# The number of active metabolic pathways is bounded by the number of cellular constraints at maximal metabolic rates

**DOI:** 10.1101/167171

**Authors:** Daan H. de Groot, Coco van Boxtel, Robert Planqué, Frank J. Bruggeman, Bas Teusink

## Abstract

Growth rate is a near-universal selective pressure across microbial species. High growth rates require hundreds of metabolic enzymes, each with different nonlinear kinetics, to be precisely tuned within the bounds set by physicochemical constraints. Yet, the metabolic behaviour of many species is characterized by simple relations between growth rate, enzyme expression levels and metabolic rates. We asked if this simplicity could be the outcome of optimisation by evolution. Indeed, when the growth rate is maximized –in a static environment under mass-conservation and enzyme expression constraints– we prove mathematically that the resulting optimal metabolic flux distribution is described by a limited number of subnetworks, known as Elementary Flux Modes (EFMs). We show that, because EFMs are the minimal subnetworks leading to growth, a small active number automatically leads to the simple relations that are measured. We find that the maximal number of flux-carrying EFMs is determined only by the number of imposed constraints on enzyme expression, not by the size, kinetics or topology of the network. This minimal-EFM extremum principle is illustrated in a graphical framework, which explains qualitative changes in microbial behaviours, such as overflow metabolism and co-consumption, and provides a method for identification of the enzyme expression constraints that limit growth under the prevalent conditions. The extremum principle applies to all microorganisms that are selected for maximal growth rates under protein concentration constraints, for example the solvent capacities of cytosol, membrane or periplasmic space.

**Author summary:** The microbial genome encodes for a large network of enzyme-catalyzed reactions. The reaction rates depend on concentrations of enzymes and metabolites, which in turn depend on those rates. Cells face a number of biophysical constraints on enzyme expression, for example due to a limited membrane area or cytosolic volume. Considering this complexity and nonlinearity of metabolism, how is it possible, that experimental data can often be described by simple linear models? We show that it is evolution itself that selects for simplicity. When reproductive rate is maximised, the number of active independent metabolic pathways is bounded by the number of growth-limiting enzyme constraints, which is typically small. A small number of pathways automatically generates the measured simple relations. We identify the importance of growth-limiting constraints in shaping microbial behaviour, by focussing on their mechanistic nature. We demonstrate that overflow metabolism – an important phenomenon in bacteria, yeasts, and cancer cells – is caused by two constraints on enzyme expression. We derive experimental guidelines for constraint identification in microorganisms. Knowing these constraints leads to increased understanding of metabolism, and thereby to better predictions and more effective manipulations.

## Introduction

Fitter microorganisms drive competitors to extinction by synthesising more viable offspring [1,2]. The rate of offspring-cell synthesis per cell, i.e., the specific growth rate, is a common determinant of evolutionary success across microbial species [1]. A high growth rate requires high metabolic rates, which in turn require high enzyme concentrations [3]. Due to limited biosynthetic resources, such as ribosomes, polymerases, energy and nutrients, the expression of any enzyme is at the expense of others [4,5]. Consequently, the selective pressure towards maximal growth rate requires the benefits and costs of all enzymes to be properly balanced, resulting in optimally-tuned enzyme expressions [6–9].

Tuning all enzyme expression levels appears to be a highly complex task. First, the genome of a microorganism encodes for thousands of reactions with associated enzymes. Second, a change in expression level of one enzyme not only affects the rate of its associated reaction, but also changes intracellular metabolite concentrations. These metabolite concentrations influence the activities of many other enzymes in a nonlinear fashion. In mathematical terms, microorganisms thus have to solve a high-dimensional nonlinear optimization problem.

Surprisingly, experiments on many different microorganisms often show simple linear relations between growth rate, enzyme expression levels and metabolic rates [10–12], and the data can often be described by coarse-grained linear models. This suggests that microorganisms in fact only use few regulatory degrees of freedom for tuning metabolic flux and protein expression. It is currently unclear why this simple, low-dimensional behaviour results from the a priori enormously complicated tuning task. Given that the tendency towards simplicity is widespread amongst microorganisms, we expected this to be due to a general –evolutionary– principle.

We found an evolutionary extremum principle: growth-rate maximization drives microorganisms to minimal metabolic complexity. We provide the mathematical proof of this principle in the Model setup and theoretical derivations section. It is derived from basic principles, more specifically from (i) mass conservation, i.e., steady-state reaction-stoichiometry relations, and (ii) enzyme biochemistry, i.e., the linear dependence of enzyme activity on the amount of enzyme and its nonlinear dependence on substrate and product concentrations. Our results provide a novel perspective on metabolic regulation, one in which the complexity is not determined by the size of the network or the rate equations, but by the constraints acting on the enzyme concentrations.

## Model setup and theoretical derivations

In this section we will introduce the class of models that we studied, and mathematically prove our main result: the extremum principle. Readers that would like to skip the mathematical proof are strongly suggested to read the biological summary of the results at the end of the section.

### The model: Evolutionary rate maximization can only be studied in a kinetic model of metabolism with constraints on enzyme concentrations

The structure of any metabolic network can be given by a stoichiometric matrix *N*, indicating which metabolites (rows) are consumed or produced in each reaction (columns). Because we can split reversible reactions in two irreversible reactions [13], we will from now on assume that all reactions are irreversible. A steady-state flux distribution is then given by a vector of reaction rates ***v*** such that there is no accumulation or depletion of metabolites, and such that all irreversibility constraints are satisfied. The solutions together form a flux cone:

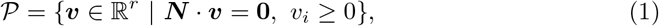

where *r* is the number of reactions. In steady state, we maximize the objective flux, which is a (linear combination of) component(s) of this flux vector. Often, the objective is chosen to be the overall cell-synthesis reaction, also called the biomass reaction *v*_BM_, which makes all cellular components in the right proportions according to the biomass composition [14].

To understand the resource allocation associated with a particular metabolic activity, we need to know the relation between the rates of enzyme-catalyzed reactions and enzyme concentrations. At constant metabolite concentrations, these are in general proportional [3] as captured by the rate equation:

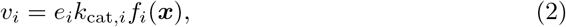

where *e*_*i*_ is the concentration of the enzyme catalyzing this reaction, *k*_cat,*i*_ is its maximal catalytic rate and *f_i_*(***x***) is the ‘saturation function’ of the enzyme, which is dependent on metabolite concentrations ***x***. This function, *f*_*i*_(***x***), is often nonlinear, includes the thermodynamic driving force, (allosteric) activation or inhibition, and other enzyme-specific effects.

To model the maximization of the cell-synthesis flux we have to account for bounds on enzyme concentrations, originating for example from limited solvent capacities of cellular compartments, or from a limited ribosomal protein synthesis capacity. We model these biophysical limits by imposing *K* constraints, each modelled by a weighted sum of enzyme concentrations:

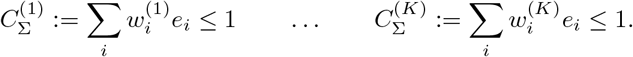

These constraints correspond to limited enzyme pools. Overexpression of one enzyme is therefore at the expense of others that are subject to the same biophysical constraint. The weights, 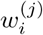, determine the fraction that one mole/liter of the *i*^th^ enzyme uses up from the *j*^th^ constrained enzyme pool. For example, for a constraint describing the limited solvent capacity of the membrane, the weight of an enzyme is the fraction of the available membrane area that is used up by this enzyme; this weight is thus nonzero only for membrane proteins. We call a constraint ‘active’ when it limits the cell in increasing its growth rate, indicating that the corresponding enzyme pool is fully used. One enzyme can belong to one, several or none of these limited pools.

Note that these constraints on enzyme concentrations are different from the constraints on reaction rates that are often used in stoichiometric methods (e.g., through Flux Balance Analysis). For these linear models, it is known -similar to what we will derive in the general, nonlinear case in this work- that few minimal pathways constitute the optimal solutions in such models [15]. However, constraints on reaction rates do not reflect the ability of microorganisms to adjust their enzyme content: any reaction rate constraint could in principle be overcome by an increase of the corresponding enzyme’s concentration. The enzyme constraints that we model are due to biophysical laws and can thus not be alleviated by metabolic regulation. These must thus be investigated to study the evolution of metabolism, although this forces us to include the complicated (and often unknown) enzyme saturation functions, *f*_*i*_(***x***), in our theory.

The number of constraints and the exact value of the weights may vary per organism. In general we expect this number to be low, and indeed not many different enzyme expression constraints have been proposed in the literature. Many aspects of microbial growth have been successfully described using constraints that are (or can be reformulated as) enzyme expression constraints, like limited reaction rates and limited solvent capacities within cellular compartments [4,5,10,16–20].

The introduction of enzyme kinetics in Equation (2) allows us to rewrite the enzyme constraints as:

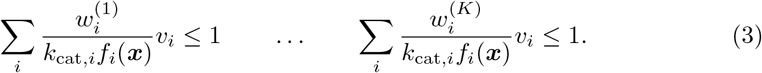

We note that, although written in terms of the fluxes, these constraints are not equivalent to the normal flux constraints used in FBA, since the weighted sums now depend on metabolite concentrations. To maximize the cell-synthesis flux, not only the enzyme concentrations should be optimized, but also the intracellular metabolite concentrations. Due to the necessary inclusion of enzyme kinetics, flux maximization is turned into a complicated nonlinear problem. This is the problem we have investigated. Remarkably, we will prove below that the solution still uses only a few minimal metabolic pathways.

### The minimal building blocks: Elementary Flux Modes

A minimal metabolic pathway is called an ‘Elementary Flux Mode’ (EFM). In words, EFMs are support-minimal subnetworks that can sustain a steady state [21]. The ‘support’ of a flux vector is the set of participating reactions: *R*(***v***) = {*j*: *v*_*j*_ ≠ 0}. That an EFM, **EFM**, is support-minimal means that if there is another flux vector, 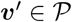, such that *R*(***v***′) ⊆ *R*(**EFM**) then we must have ***v***′ = α**EFM** for some *α* ≥ 0. Another way of phrasing this is that none of the used reactions can be set to zero in the EFM without violating the steady state condition. These metabolic subnetworks turn out to be determined completely by reaction stoichiometry, and thus for their identification no kinetic information is needed. However, because of the many combinations of parallel, alternative metabolic routes in metabolic networks, it is currently computationally infeasible to find the complete set of EFMs in a genome-scale network [22,23].

We exploit EFMs because any steady state flux distribution can be decomposed into positive linear combinations of EFMs. Indeed, Gagneur and Klamt showed that in any metabolic network in which reversible reactions are split in two irreversible reactions, the EFMs coincide with the extreme rays of the pointed polyhedral cone 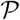 [13]. We can thus write:

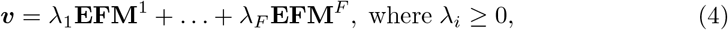

where the multiplication factors *λ*_*i*_ denote how much the *i*^th^ EFM is used and *F* denotes the total number of EFMs in the network. Equation (4) shows that EFMs are the basic building blocks of steady state metabolism. Note that, although the Elementary Flux Modes are constant vectors defined by stoichiometry, the *λ*_*i*_-factors are variable and dependent on metabolite concentrations. We will make this dependence more precise in S1 Appendix Section 5.

EFMs are defined up to a constant: if ***v*** is an EFM, then so is *α**v*** for any *α* ∈ ℝ_≥0_. This has two important consequences. First, the ratio between flux entries in an EFM are fixed, and second, we may scale one entry of an EFM to 1. We will consider optimisation of some objective flux *v*_*r*_ at steady state. Therefore, we only need to consider those EFMs which have a nonzero *r*^th^ flux value. Without loss of generality, we can make this the last entry in the vector, and we will always scale this entry to 1. The *i*^th^ EFM can thus be denoted by 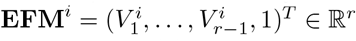, with all 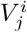 uniquely determined by stoichiometry. The *λ*_*i*_ factor in (4) can now be reinterpreted as the flux that **EFM**^*i*^ contributes to the objective flux.

Using EFMs, we can unambiguously quantify metabolic complexity as the number of flux-carrying Elementary Flux Modes. We call an EFM a minimal unit of metabolic complexity because the flux values through its participating reactions can only scale with one overall factor. A flux distribution that is a sum of *K* EFMs thus has *K* flux degrees of freedom. A small number of degrees of freedom gives rise to metabolic behaviour with simple relations between the growth rate and flux values.

### The cost vectors: a low-dimensional view at metabolism

Given *K* constraints, we can, for each EFM, calculate the cost per constraint for making one unit objective flux. These *K* costs turn out to comprise all relevant information for growth rate optimisation. Therefore, we will here define the *cost vectors* that have these costs as their entries. We will use the cost vectors to study metabolism in low-dimensional *constraint space* throughout this paper.

As discussed above, we can rescale each EFM such that it is a vector of the form 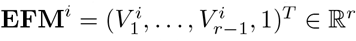. To produce one unit objective flux, we thus need a flux of 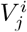 through reaction *j*. Since we have *v_j_* = *k*_cat,*j*_*e_j_ f_j_*(***x***), we get

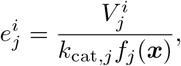

where 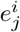 denotes the necessary concentration of enzyme *j* for one unit objective flux through EFM *i*. We can then define the *cost vector* ***d***^*i*^(***x***) for the *i*^th^ EFM, with components given by the total costs that this EFM brings per constraint:

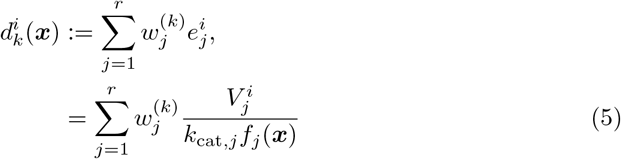

Because enzyme kinetics determine the enzyme concentrations and thereby the enzymatic costs, it is unlikely that several EFMs have exactly the same costs. Different EFMs use at least one different enzyme, and it is highly improbable that the necessary concentrations of these different enzymes are exactly the same real number. If one of these non-overlapping enzymes is part of a constrained pool, the EFMs will thus have different costs.^1^ If, however, none of the non-overlapping enzymes are part of the constrained pools, several EFMs can indeed have the same costs. To deal with this case we introduce the notion of *equivalent EFMs*.

#### Definition 1.

*Given a set of constraints*, 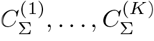, *two EFMs*, **EFM**_1_, **EFM**_2_, *are called* equivalent with respect to the constraints *if their associated cost vectors are equal: **d***^1^(***x***) = ***d***^2^(***x***).

Because the cost vectors play a central role in the whole paper, we illustrated their definition and use in Figure 1. Many of our results followed from studying these cost vectors.

**Fig 1.**
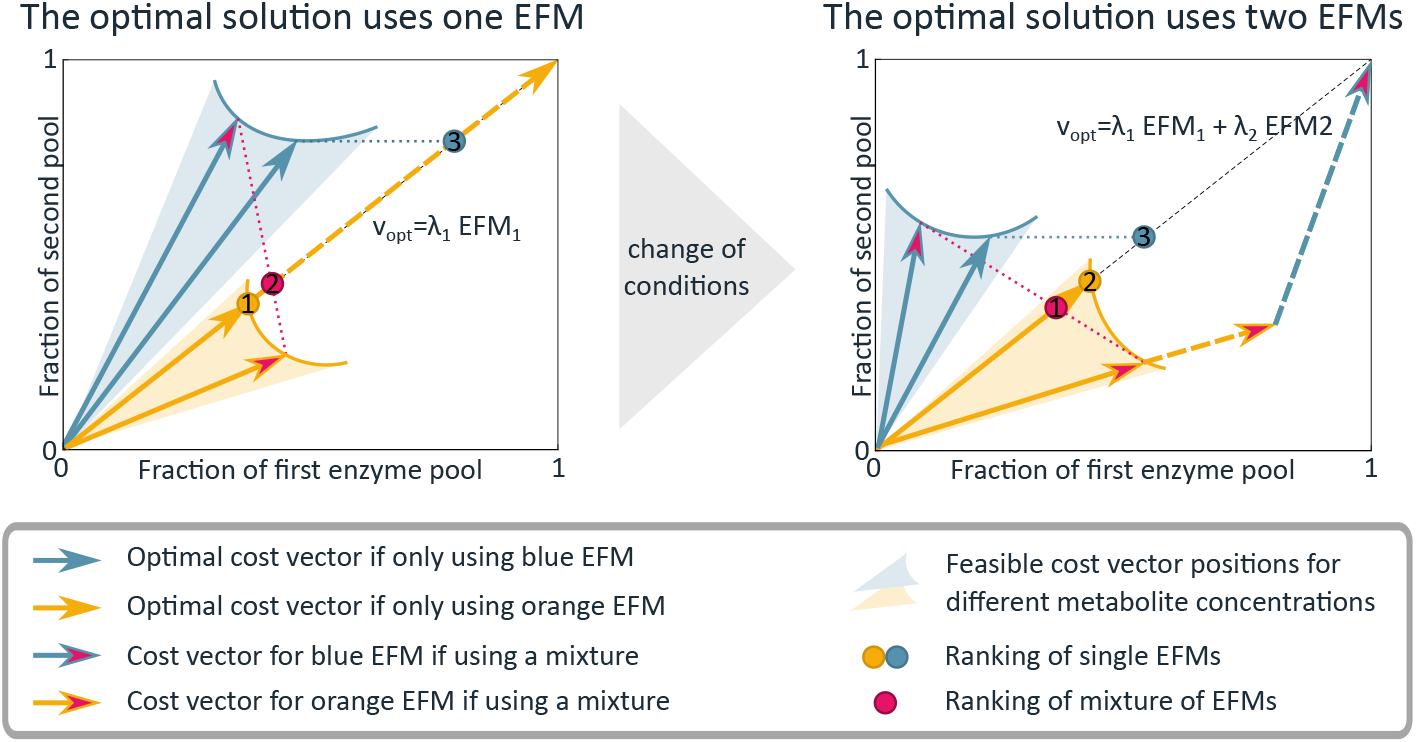
The cost vector formalism shows what determines the number of EFMs in the optimal solution. We here consider a simplified model with 2 EFMs (blue and orange), and 2 constraints. In reality, the costs of many more EFMs have to be compared, and potentially also of more constraints. The *cost vector* 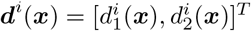 of the *i*^th^ EFM denotes the fractions of the first and second constrained enzyme pool that this EFM uses when producing one unit of objective flux. The cell-synthesis flux produced by EFM *i* is denoted by *λ*_*i*_, and the corresponding enzyme costs are *λ*_*i*_*d*_*i*_(***x***). The cost of mixing EFMs 1 and 2 corresponds to the weighted sum of the cost vectors: *λ*_1_***d***^1^(***x***) + *λ*_2_***d***^2^(***x***). The mixture is feasible as long as none of the constraints is exceeded: *λ*_1_***d***^1^(***x***) + *λ*_2_***d***^2^(***x***) ≤ **1**. The objective value, *λ*_1_ + *λ*_2_, is maximized by fitting a vector sum of as many vectors as possible in the constraint box. This solution is shown by the dashed vectors. The pure usage of one EFM with off-diagonal cost vector leads to underuse of one constraint, while diagonal cost vectors can exhaust both constrained pools. A mixture of EFMs will always be a combination of an above-diagonal and a below-diagonal vector. All EFMs and mixtures thereof, can be ranked by a dot on the diagonal that denotes the average cost per unit cell-synthesis flux (see Lemma 4 in S1 Appendix for a proof). Pure usage of above-diagonal cost vectors is ranked by projecting the cost vector horizontally to the diagonal, while pure usage of below-diagonal vectors is ranked by vertical projection. Mixtures are ranked by placing a dot at the intersection of the diagonal with the line between the two cost vectors. The (mixture of) EFM(s) with the lowest average cost (i.e., with the dot closest to the origin) leads to the highest growth rate (the mathematical proof is included in S1 Appendix). The enzymatic costs of an EFM depend on the intracellular metabolite concentrations, i.e., the saturation of enzymes. The shaded regions indicate alternative positions for the cost vectors at different intracellular metabolite concentrations, two of them are shown. The blue and orange cost vectors lead to the highest growth rate when using only that EFM. We see that in the left figure the orange EFM gives rise to a higher growth rate. Upon a change of environmental conditions, the cost vectors can change, and the mixture of EFMs can become better than either single EFM (right figure). A change like this would lead to a change in metabolic behaviour.

### The extremum principle: the number of active EFMs is determined by the number of constraints on enzyme expression

We here prove the main result of this study, the extremum principle. For a general metabolic model, as introduced above, it states a necessary condition for a flux vector 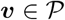 to be a maximizer of the objective flux.

#### Theorem 1.

*Consider a metabolic network characterized by the stoichiometric matrix **N**. Let v_r_ be an objective flux, which is to be maximized at steady state, under K linear-enzymatic constraints of the form*:

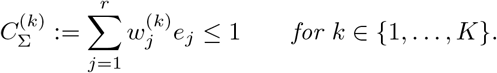

*Then, at most K non-equivalent Elementary Flux Modes are used in the optimal solution*.

#### Proof.

We assumed that *v*_*j*_ ≥ 0 for all reactions in the network because, without loss of generality, we split all reversible reactions into a forward and a backward reaction [13]. Let us for now also assume that none of the EFMs are equivalent (where equivalence is defined according to Definition 1) we will handle the case with equivalent EFMs at the end of the proof.

According to Equation (4), the optimal solution can always be expressed as a conical combination of EFMs. As before, we rescale every EFM such that it is a vector of the form 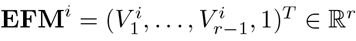. The objective flux for a flux vector ***v*** can now be written as

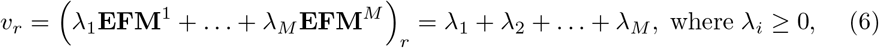

where *M* is the number of EFMs containing a nonzero *v*_*r*_. Since the EFMs are fixed vectors, the *λ*_*i*_ become our optimisation variables. Since 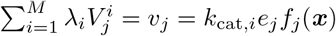, we have

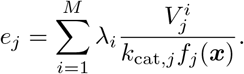

This allows us to rewrite enzyme constraint 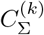 as

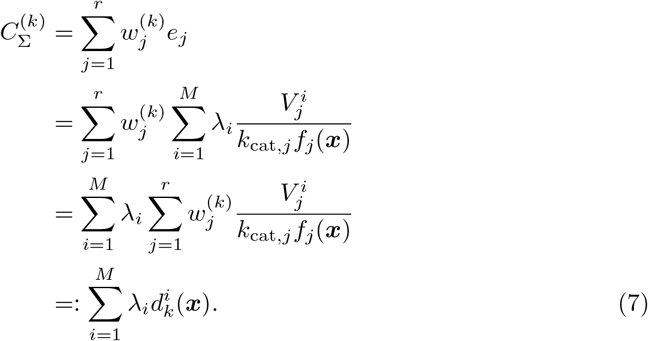

In the last step, we recognized the cost vector components defined in Equation (5).

The *k*^th^ entry of cost vector i denotes the cost for the enzymes in constraint *k* (the *k-enzymes*) to obtain one unit of objective flux through **EFM**^*i*^ (and therefore also the enzymatic cost to increase this flux by some factor). We can rewrite our optimization problem in terms of these cost vectors. We will hereby designate each metabolite concentration as either external, ***x***^*E*^, or internal, ***x***^*I*^, such that: ***x*** = (***x**^E^, **x***^*I*^). This distinction is important, because the external concentrations are given by the environment and therefore part of the parmeters of the optimisation problem, while the internal concentrations can be tuned by the cell and are therefore part of the solution. We need to solve

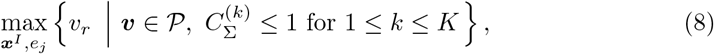

and using Equations (6) and (7), this is equivalent to

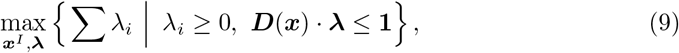

where ***D*** = [***d***^1^(***x***) · · · ***d**^r^*(***x***)] is the cost vector matrix. The relation ***D***(***x***) · ***λ*** ≤ 1 shows that the optimal ***λ*** vector indeed depends on the metabolite concentrations ***x***, as was indicated below equation (4).

Following^2^ Wortel et al. [24], we now use a subtle mathematical argument. We fix ***x*** = ***x***_0_, so that the enzyme saturations *f*_*j*_(***x***_0_) are constant. This will give us a fixed cost vector for each EFM. The remaining optimization problem is then visualized in Figure 1, where cost vectors of some EFMs are plotted in a box of constraints. Finding the optimal solution is equivalent to finding a sum of scalar multiples of the cost vectors without leaving the box of constraints while maximizing the sum of these multiplicities. The example in Figure 1 shows only 2 constraints, but in general we would have *M* vectors in a *K*-dimensional cube.

In the general case, it might seem intuitive that *K* constraints lead to the usage of at most *K* EFMs since all *K* linearly-independent vectors form a basis of a *K*-dimensional space. We can thus always take a combination of *K* vectors to reach the point where all constraints are met with equality. However, we should be careful because we could end up with negative *λ*’s for some of the EFMs. We continue with the proof by rewriting the problem in a Linear Programming (LP) form,

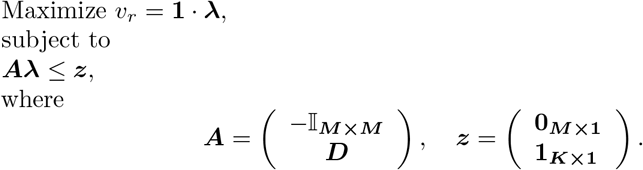

The solutions of this linear programming problem form a polytope in ℝ^*M*^, bounded by the hypersurfaces given by the constraints. The most important theorem of LP teaches us that an optimal solution is found among the vertices of this polytope. The dimension of such vertices is zero, which means that optimal solutions satisfy at least *M* of the *K* + *M* constraints with equality. Therefore at most (*K* + *M*) – *M* = *K* constraints can be satisfied with strict inequality. These *K* inequalities could be concentrated in the *λ*_*i*_ ≥ 0 part, which means that the corresponding *K* Elementary Flux Modes are used. Thus, an optimal solution can use no more EFMs than there are active constraints in the system, thereby proving the theorem for any arbitrary vector of metabolite concentrations ***x***.

There is one possible exception to the above reasoning. Let’s say that *K* EFMs are used in the optimum: 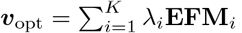. If one EFM, say **EFM**_*K*_, has an equivalent EFM, say **EFM**_*K*+1_, then we can replace the usage of EFM *K* by any convex combination of EFMs *K* and *K* +1 and the solution will still be optimal. So, in the case that the costs of several EFMs are the same, the optimal flux vector could consist of more EFMs than the number of constraints. That’s why the theorem only tells us that no more than *K non-equivalent* EFMs are used in the optimal solution.

Finally, it follows that, since the theorem is true for any set of metabolite concentrations ***x***, it is of course also true for the *optimal* set, 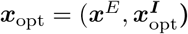.

We note that the optimal internal concentrations, the choice of EFMs, and thereby the optimal enzyme concentrations, all depend on the external concentrations ***x***^*E*^. Which specific EFMs are the optimal ones, thus does not follow directly from the theorem.

We think that the case where several EFMs are equivalent is not very common in biology. First, the constraints on enzyme expression are due to biophysical limits and we expect these to act on many enzymes together. This reduces the chance of having several EFMs that use exactly the same enzymes within the constrained pool of enzymes. Second, even if several EFMs would use the same enzymes, then the enzyme costs depend on the enzyme saturations, and these depend on the optimal metabolite concentrations. These optimal concentrations depend on the rest of metabolism, such that the non-overlapping part of the EFMs can still influence the enzyme costs. For these two reasons, we will assume in the rest of this work that EFMs are generally not equivalent.

The previously published theorem that maximal specific flux, 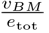, is attained in an EFM [24,25] is a special case of Theorem 1. In the cost vector formalism that we described in Figure 1, it is visualized by cost vectors on a line rather than in a box, because there is only one enzymatic constraint (total enzyme concentration is bounded). In this case, there is indeed a shortest cost vector for all but a negligible subset of situations (as discussed in the proof).

The following corollary can be used to find out how many constraints are active when we observe a certain number of active EFMs. It is the contrapositive of Theorem 1 and therefore mathematically equivalent. The reason that it is stated separately is the difference in biological applicability: the theorem is a predictive statement while the corollary is descriptive. As we will see in the Results section, the theorem tells us that metabolic complexity is low because the number of enzymatic constraints is typically low. The corollary however, enables us to infer from experimental data how many constraints must be active, and thus gives us physiological insight from population-level data.

#### Corollary 2.

*If a flux v_r_ is optimized and K non-equivalent Elementary Flux Modes are used, then at least K linear enzymatic constraints must be active.*

EFMs are not the only set of building blocks that we could have used. In the context of Flux Balance Analysis, constraint-based rate maximization can be studied by calculating Elementary Flux Vectors (EFVs) [26,27], which are the minimal pathways that generate all flux distributions that satisfy not only the steady-state assumption, but also the additional constraints. Therefore, for fixed enzyme saturations and constraints, EFVs provide a set of feasible building blocks of which convex combinations automatically satisfy all constraints. However, since every EFV is a conical combination of EFMs, and since we wanted to study evolutionary growth-rate maximization, we preferred to do our analysis on the set of EFMs. This is because the EFMs provide a set of invariant (at least on timescales on which stoichiometry is not evolved) objects for which regulatory circuits can be evolved. In principle, the extremum principle can also be written in terms of EFVs. We can show, in a similar manner as in the proof above, that rate-maximal solutions will use only one EFV, which is a convex combination of at most *K* EFMs.

### Biological summary of the extremum principle and its proof

The extremum principle, stated in Theorem 1, is a statement about all metabolic networks, independent of the network size, topology, or the specific enzyme kinetics. All microorganisms are subjected to a small number of enzymatic constraints, and all metabolic networks have Elementary Flux Modes as their building blocks: minimal pathways that make all cellular components from external sources. The fluxes through the participating reactions in an EFM can only be rescaled with one overall factor. We concluded that the use of an additional EFM thus only adds one flux degree of freedom, so that experimental data will show low complexity if few EFMs are used. We then proved the extremum principle, stating that the number of flux-carrying EFMs in the maximal growth rate solution is always bounded by the number of constraints on enzyme expression. As a whole, this leads to the prediction that microbial behaviour will show low complexity.

In the proof, we compared the costs and benefits of the different EFMs. To be precise, we rescaled the EFMs such that the benefit of each EFM was equal: they all give one unit of objective flux. If we have *K* constraints, we also have *K* different costs for which we need to compare the different EFMs. We showed that the optimal solution is a combination of up to *K* of these EFMs. This is in accordance with the intuition that one EFM can be selected for each constraint because it has a low cost with respect to this constraint.

To find the proof, we developed a framework using cost vectors. In Figure 1 we summarize how this framework allows us to study high-dimensional metabolism in the few dimensions that actually matter: we can compare the enzyme costs of all EFMs in the low-dimensional ‘constraint space’ defined by the limited enzyme pools. This perspective enables us to design experiments that characterize the active biophysical constraints, as we will discuss in the Results section.

## Results

### The metabolic complexity is typically very low

We called an EFM a minimal unit of metabolic complexity because the ratios between the fluxes through all participating reactions are fixed, and none of its reactions can be removed. Consequently, a microorganism that uses one EFM can only change all reaction rates with the same factor. In other words, there is only one regulatory degree of freedom, instead of many if all reaction rates could have been tuned separately. In this case, flux values can be described by only one straight line. This becomes more complex when the number of flux-carrying (active) EFMs increases. Using this knowledge, the number of active EFMs can be estimated from flux measurements.

We re-analysed data from carbon-limited chemostats and indeed observed that uptake rates of glucose and oxygen could be described by a straight line for a large range of growth rates, testimony of single EFM usage (S1 Appendix Section 8). A possibility that we cannot exclude, however, is that many EFMs are used, but that these EFMs all have the same relation between growth rate, glucose uptake and oxygen uptake. On the other hand, the experimentally measured linear growth laws between cellular building blocks and growth [11,12,18], and the success of coarse-grained models [4,5], do provide some additional indications of the usage of a small number of EFMs. A more definite proof could be found in two ways. First, if many different reaction rates are measured in balanced growth across slightly different environments, or second, if all internal fluxes in the cell are measured, and complete knowledge of the stoichiometric network is available. However, to our knowledge, currently available fluxome datasets were collected across mutants, or across very different growth environments, making them unsuitable for our purposes. For now, based on the available data, we cautiously argue that the number of simultaneously active EFMs is typically very low, in the order of 1 to 3. That microorganisms would choose only a handful of EFMs out of billions of alternatives is in accordance to our extremum principle, Theorem 1. These alternatives are apparently not evolutionarily equivalent, and only a small number has been selected because of their superior kinetics.

### The extremum principle: the low number of biophysical constraints causes low metabolic complexity

The extremum principle states: when the rate of a particular reaction in a metabolic network is maximized, the number of flux-carrying EFMs is at most equal to the number of constraints on enzyme concentrations that limit the objective flux. In particular, the principle holds for the cell-synthesis reaction. Therefore, if the number of active constraints is low, so is the number of active EFMs at maximal growth rate. This is the basis of our finding that maximal growth rate requires minimal metabolic complexity, and this extends the result that rates are maximized by one EFM under a total protein constraint [24,25]. This earlier result could not explain –from a resource allocation perspective– datasets in which several metabolic pathways are used, such as overflow metabolism, metabolic switches, and the expression of unutilized proteins.

The extremum principle holds regardless of the complexity of the metabolic network, i.e., of its kinetics and its structure. The metabolic complexity is only determined by the number of active constraints; the kinetics and structure subsequently determine which EFMs are optimal and selected by evolution - as illustrated by *in silico* evolution of metabolic regulation towards only one active EFM [34]. For this reason, also genome-scale metabolic models, which contain all the annotated metabolic reactions that a microorganism’s genome encodes [35], and even the ones that have been studied with different additional resource constraints [36,37], behave qualitatively similar to simplified core models. Coarse-grained models can thus be used without loss of generality, which greatly facilitates our understanding of metabolic behaviour.

Using the cost vector formalism that we used in the proof of Theorem 1, we can study metabolism in the low-dimensional constraint space, instead of in the high-dimensional flux space (see Figure 3). In the case of two constraints (also illustrated in Figure 1), the extremum principle states that both constrained enzyme pools can always be fully used with two cost vectors (EFMs), not more. However, an EFM with a diagonal cost vector can make full use of both pools on its own: hence, the number of EFMs that maximize flux can also be less than the number of active constraints. Another instance in which only one EFM is optimal, is when all cost vectors lie above or below the diagonal. In this case, there is only one active constraint; the other pool does not limit the total possible flux of the system under these conditions. We have derived the necessary and sufficient conditions under which it is optimal to use EFMs in mixtures (S1 Appendix Section 5). Plotting the cost vectors for different internal metabolite concentrations also shows that the length and direction of the cost vectors are affected by metabolite concentrations via enzyme kinetics (depicted by the shaded areas in Figure 1). We show in S1 Appendix Section 5 that this metabolite-dependency makes it much more probable that less than *K* EFMs are used in a system with *K* constraints, because internal concentrations can be changed to make cost vectors diagonal.

**Fig 2.**
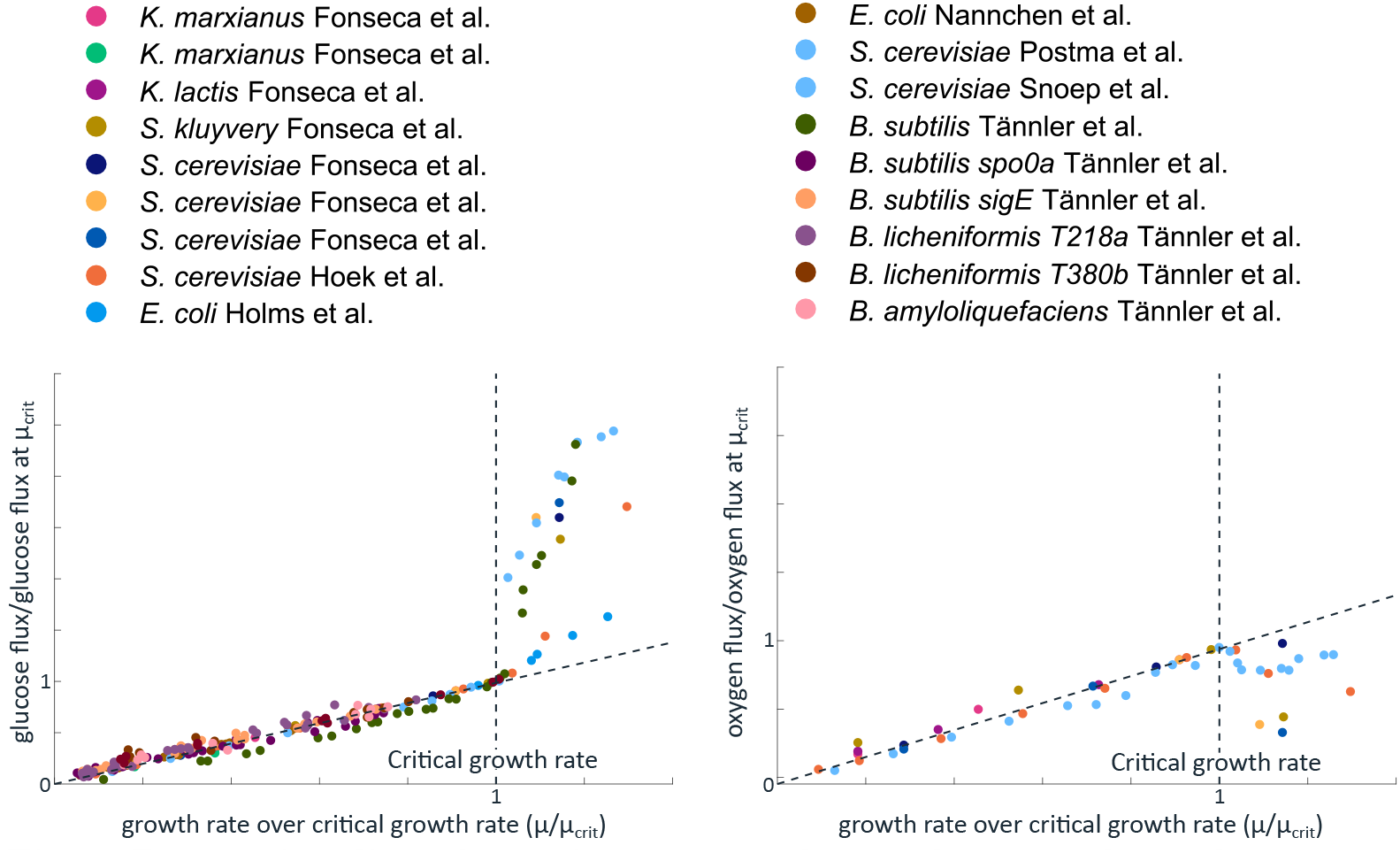
Proportionality of reaction rates and growth rates, shown by many microorganisms, is an indication of low metabolic complexity. Measured uptake rates [28–33] were gathered from experiments in which growth rate was varied in carbon-limited chemostats. For each species we normalized the measured growth rate to the so-called critical growth rate: the growth rate at which the production of overflow products starts. Uptake rates were normalized relative to the uptake rate of the species at the critical growth rate. Up to the critical growth rate, all microorganisms show a simple proportional relation between the growth rate and uptake rates of glucose and oxygen. In Section 8 we explain why this proportionality is an indication of the usage of only one EFM. After the critical growth rate, the reaction rates are no longer proportional, a phenomenon called overflow metabolism.

**Fig 3.**
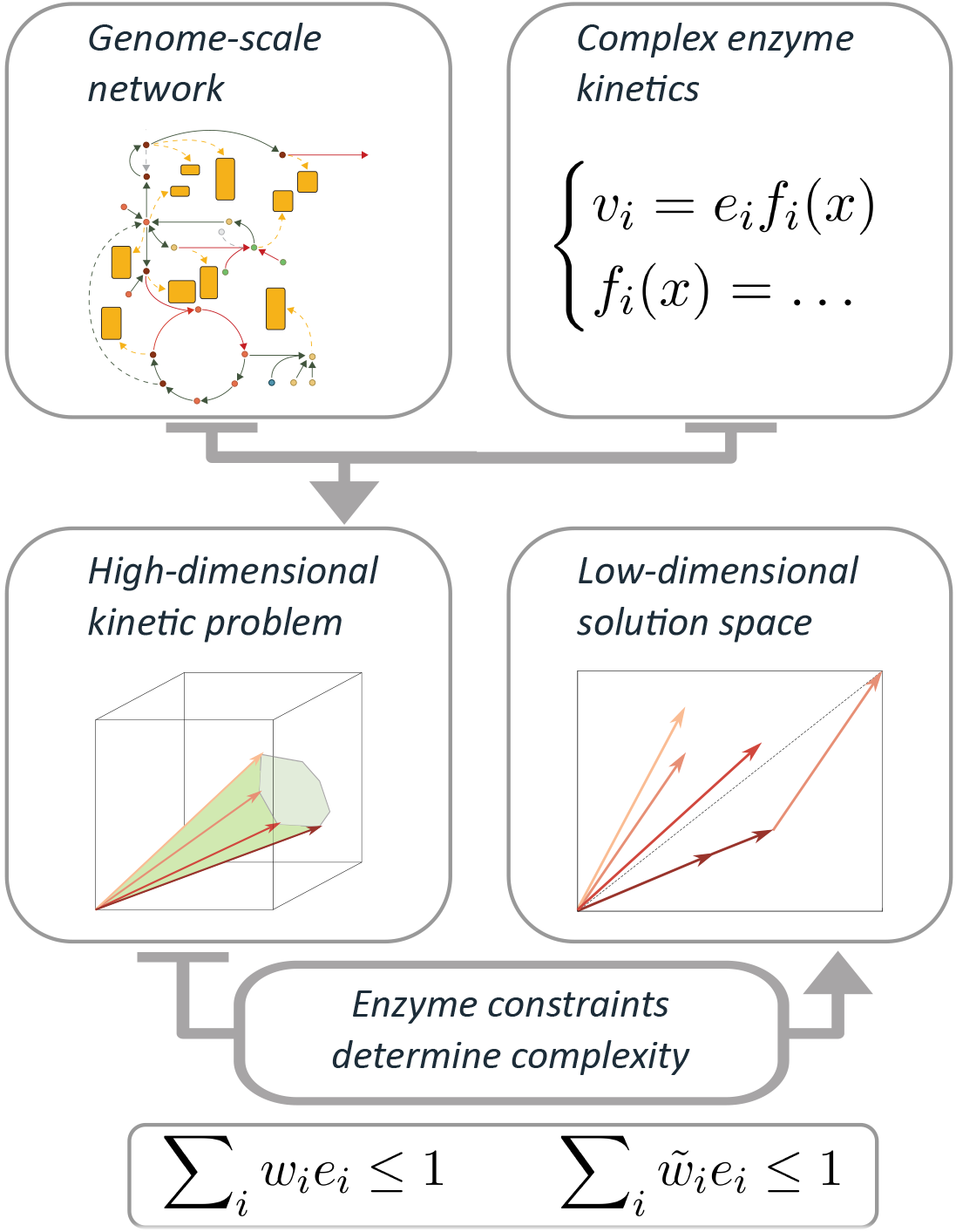
Illustration of the extremum principle. The extremum principle states that the dimensionality of the solution space is determined by the number of enzyme-expression constraints, rather than by the dimensionality of the metabolic network. The constraints result from biophysical limits, e.g., limited solvent capacities within cellular compartments. Our cost vector formalism, explained in Figure 1, enables us to analyze metabolism in the low-dimensional constraint space, instead of in the high-dimensional flux space that is normally used.

### The number of enzymatic constraints can be inferred from experimental data: the extremum principle applied to overflow metabolism

A well-known phenomenon observed across microbes is overflow metabolism: the apparently wasteful excretion of products. Examples are the aerobic production of ethanol by yeasts (Crabtree effect), lactate by cancer cells (Warburg) or acetate by *Escherichia coli* [4,38,39]. The onset of overflow metabolism is generally studied as a function of growth rate (e.g., in chemostats where the growth rate is set by the dilution rate of the culture). Before some critical growth rate, cells fully respire, but when the growth rate is increased above some critical value, respiratory flux decreases and the flux of overflow metabolism emerges.

According to our theory, an additional enzymatic constraint must have become active at the critical growth rate (see Figure 2). Below the critical growth rate, the respiratory flux is proportional to the growth rate, which is a characteristic of single EFM usage (see S1 Appendix). Above the critical growth rate however, the decreasing respiratory flux and increasing overflow flux indicate that at least two EFMs and therefore two constraints must be active. Indeed, current models of overflow metabolism all use such an additional constraint, but the biophysical nature of the first constraint (mostly an uptake constraint) is often kept implicit. Many explanations of overflow metabolism therefore appeared to have only one constraint, for example linked to total protein [4], or membrane protein [40], but within our theory an optimal flux distribution with two EFMs is only possible with at least two constraints.

We can gain more insight on overflow metabolism by applying the cost vector formalism on a coarse-grained model (Figure 4). Note however, that this model has an illustrative purpose only, to show that overflow metabolism can be easily explained with two enzyme expression constraints. We do not claim that the imposed constraints are the real constraints; for this, experiments are needed, as we will explain later. The model includes a respiration pathway and an acetate overflow branch. All steps include enzyme kinetics, and constraints are imposed on two enzyme pools: total cytosolic protein, and total membrane protein. We model overflow metabolism as a function of the glucose concentration.^3^ At low extracelullar glucose concentrations, all cost vectors have high membrane costs and lie above or at best at the diagonal (as the membrane constraint is on the y-axis): the membrane pool limits substrate uptake and therefore favours efficient use of glucose via respiration. Our core model predicts that, as extracellular glucose concentrations increase, so does the saturation level of the glycolytic enzymes such that flux can increase without a change in protein level. Consequently, across a large range of external substrate concentrations pure respiration leads to maximal growth rate by fully exploiting the two available enzyme pools. The membrane constraint is however more growth-limiting, i.e., loosening this constraint will give a larger growth rate benefit. At high glucose concentrations, transporters are more saturated (cost vectors become shorter in the membrane direction) and the respiration cost vector becomes below-diagonal: pure respiration will leave the membrane protein pool underused, while the cytosolic pool limits respiration. A better strategy is to respire less and make some of the cytosolic pool available for another EFM that can exploit the underused membrane pool. The net outcome is that a mixture of EFMs attains a higher growth rate than either of the two EFMs alone.

**Fig 4.**
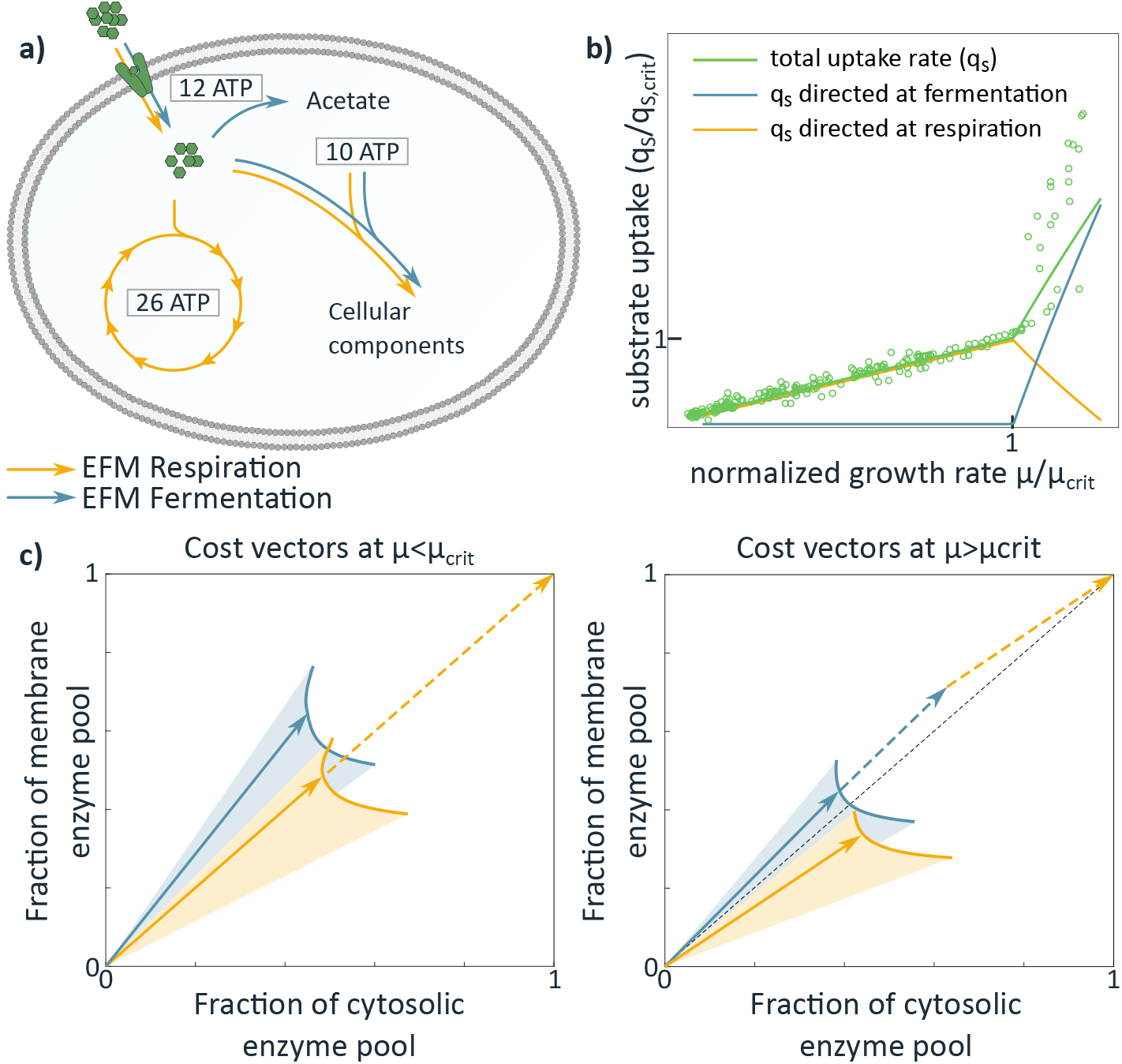
The cost vector formalism provides insight in how growth rate maximization leads to overflow metabolism. **a)** A core model with two EFMs that individually lead to cell synthesis (orange: respiration and blue: acetate overflow). All considered reactions have an associated enzyme, whose activity depends on kinetic parameters and the metabolite concentrations. We varied growth rate by changing the external substrate concentration. Given this external condition, the growth rate was optimized under two enzymatic constraints (limited cytosolic enzyme Σ *e*_*i*,cyto_ ≤ 1 and limited membrane area *e*_transport_ ≤ 0.3). **b)** The predicted substrate uptake fluxes directed towards respiration and overflow are in qualitative agreement with the experimental data (shown before in Figure 2) of several microorganisms scaled with respect to the growth rate (*μ*_crit_) and uptake rate (*q*_crit_) at the onset of overflow [4,38,39]. **c)** The cost vectors (solid arrows) of the two EFMs before (left) and after (right) the respirofermentative switch. The *x*-coordinate of the cost vectors denote the fraction of the cytosolic volume that is needed to produce one unit objective flux with the corresponding EFM. The *y*-coordinate shows the necessary fraction of the available mebrane area. The position of the cost vectors are shown for the optimized metabolite concentrations; the shaded regions show alternative positions of the cost vectors at different enzyme and metabolite concentrations. The dashed vectors show the usage of the EFMs in the optimal solution.

We think that many published explanations of overflow metabolism are unified by the extremum principle. The added value is not that it gives yet another model that qualitatively captures overflow metabolism, but rather that it explains why published models are successful by offering an overarching theory. Indeed, we show in S1 Appendix Section 4 that explanations for overflow metabolism offered by other modeling methods, imposing different constraints, such as coarse-grained whole cell models [4,5] and constraint-based genome-scale M-models [19,41–43] are mathematically all instances (or simplifications) of the exact same constrained optimization problem that we study here. Their maximizers thus all follow the extremum principle, and overflow metabolism must be the result of a second constraint that becomes active. So-called ME-models [36] fall under a slightly different class of mathematical problems, but the onset of overflow metabolism is still caused by an additional active constraint. However, since the above explanations all capture the phenomenon with different constraints and solve the same mathematical problem, we cannot conclude on the mechanistic nature of the constraints, yet.

### The identity of the enzymatic constraints can be revealed by experimental perturbations

We can predict the effect of experimental perturbations on metabolism with the cost vector formalism. Examples of such perturbations are the expression of non-functional proteins or the inhibition of enzymes, which can respectively be interpreted as reducing a limited enzyme pool, or lengthening the cost vectors. The effect of such perturbations on growth, when two EFMs are expressed, was analysed in the cost vector formalism (see S1 Appendix Section 6 and 7 for the analysis). In Figure 5a-d we predict the (qualitative) effect of reducing the accessible area in constraint space for two cases (i) reduction of both enzyme pools by the same amount; or (ii) reduction of only the first constrained pool. We subsequently compare these predictions with the perturbation experiments carried out by Basan et al. [4] (see SI for a mathematical analysis).

**Fig 5.**
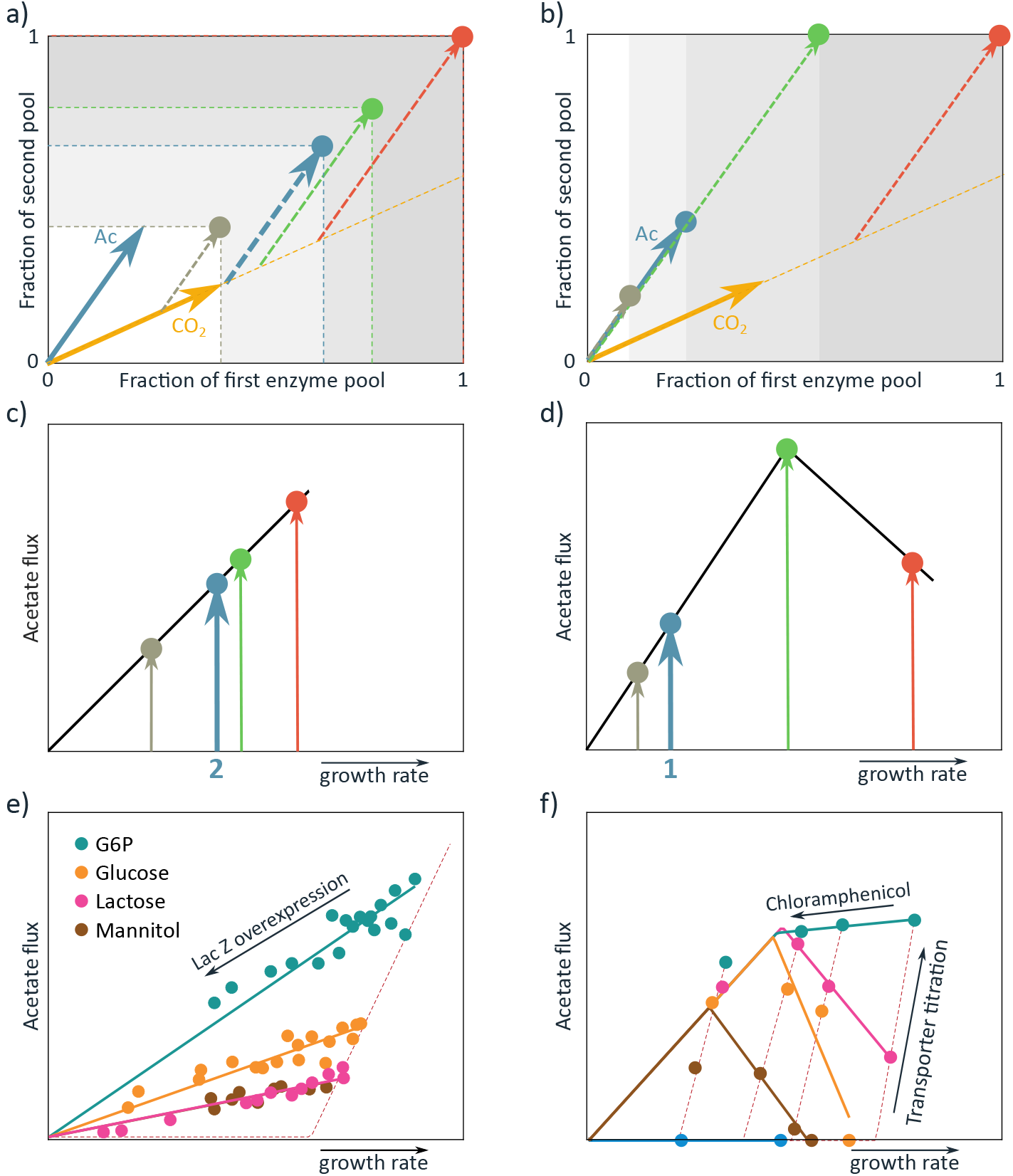
Predictions and experimental results of the perturbation of the size of limited enzyme pools during growth using a mixture of EFMs. In the cost vector plots, panels **a)** and **b)**, the red vector denotes the optimal solution in the unperturbed organism. Upon experimental perturbation, the available area in constraint space can change, indicated by the shaded grey areas. The green, blue, and grey vectors show the new optimal solutions under increasingly strong perturbations. The predicted effect on the flux through the acetate branch is shown in panels **c** and **d)**. **a,c)** Analysis of perturbations that tighten both protein pools with the same amount shows that flux and growth rate will decrease proportionally, as observed experimentally **(e)**) for the overexpression of LacZ on different carbon sources (data from Basan et al. [4]). **b,d**) Perturbations that tighten an enzyme pool that is mostly used by one EFM (here denoted by CO_2_) initially cause an increase in flux through the other EFM in the mixture(*Ac*). Eventually, at stronger limitations, this flux also decreases. **f)** This behaviour is observed, a.o., for translation inhibitor experiments using chloramphenicol (S1 Appendix Section 7).

With this analysis, we suggest a broadly-applicable experimental approach for validating likely growth-limiting constraints. Given a candidate constraint, the theory suggests a perturbation of the size of the corresponding limited enzyme pool, e.g., by the expression of a nonfunctional protein in this pool. Then, the effect of this perturbation on the flux through the active EFMs can be compared with the predictions, as in Figure 5. Now, we can validate or falsify whether certain limited enzyme pools are truly growth-limiting.^4^

The perturbation predictions can also be used to re-interpret published experiments. For example, the overexpression of the unused protein LacZ coincides with our predicted effect of an equal reduction of two enzyme pools (Figure 5e). The cost of making the cytosolic protein LacZ thus takes up an equal fraction of both constraints. We think this can be explained because LacZ can be considered an average protein in terms of resource requirements. Since metabolism was already tuned to optimally use both limited enzyme pools, all EFMs will now require more of both limited enzyme pools to maintain the growth rate (the cost vectors are lengthened). Therefore, the additional synthesis costs reduce both constrained pools to a similar extent. As a consequence, this analysis cannot decide on the biological interpretation of the constraints.

The addition of chloramphenicol is an example where our analysis does indicate that one enzymatic pool is affected more than the other (Figure 5f)). Chloramphenicol inhibits translation and the cell therefore needs a larger number of ribosomes per unit flux. This again adds a cost for protein synthesis, thereby reducing both pools. The dataset however shows that chloramphenicol has a more dominant effect on the first pool (x-axis) than on the second pool (y-axis). This means that the increased number of ribosomes has an additional effect on the first pool, which could well be related to the large cytosolic volume that the ribosomes take up. This suggests that one of the constrained pools is the sum of cytosolic proteins.^5^

### Our kinetic, constraint-based approach provides novel biological insight

Under-utilization of enzymes appears to be in conflict with optimal resource allocation. For example, Goel et al. [44] studied the switch of *L. lactis* from mixed-acid fermentation to homolactic fermentation. Since they found constant protein expression as a function of growth rate, they concluded that this metabolic switch cannot be explained from protein cost considerations. However, in Figure 6a) we show that a kinetic model that incorporates different strengths of product inhibition of ATP onto the fermentation pathways can lead to the experimentally observed behaviour when protein allocation is optimized. In our model, the saturation of homolactic fermentation enzymes rapidly increases with growth rate, while the saturation of mixed acid fermentation enzymes decreases slightly due to the increased product inhibition of ATP. As such, metabolic flux can be reallocated without a change in protein allocation (we provide the details in S1 Appendix Section 10). Another example is the expression of large fractions of under-utilized proteins by *E. coli* at low growth rates [45]. This is also in agreement with optimal resource allocation when one considers the kinetics of enzymes, such that their saturation with reactants is variable. In these two examples, the underutilization of proteins is thus used as an indication that microorganisms do not optimally allocate their resources. We here showed that these supposed counterexamples can in fact be in agreement with optimal resource allocation when one considers a kinetic model, thus including variable metabolite concentrations and enzyme saturations.

**Fig 6.**
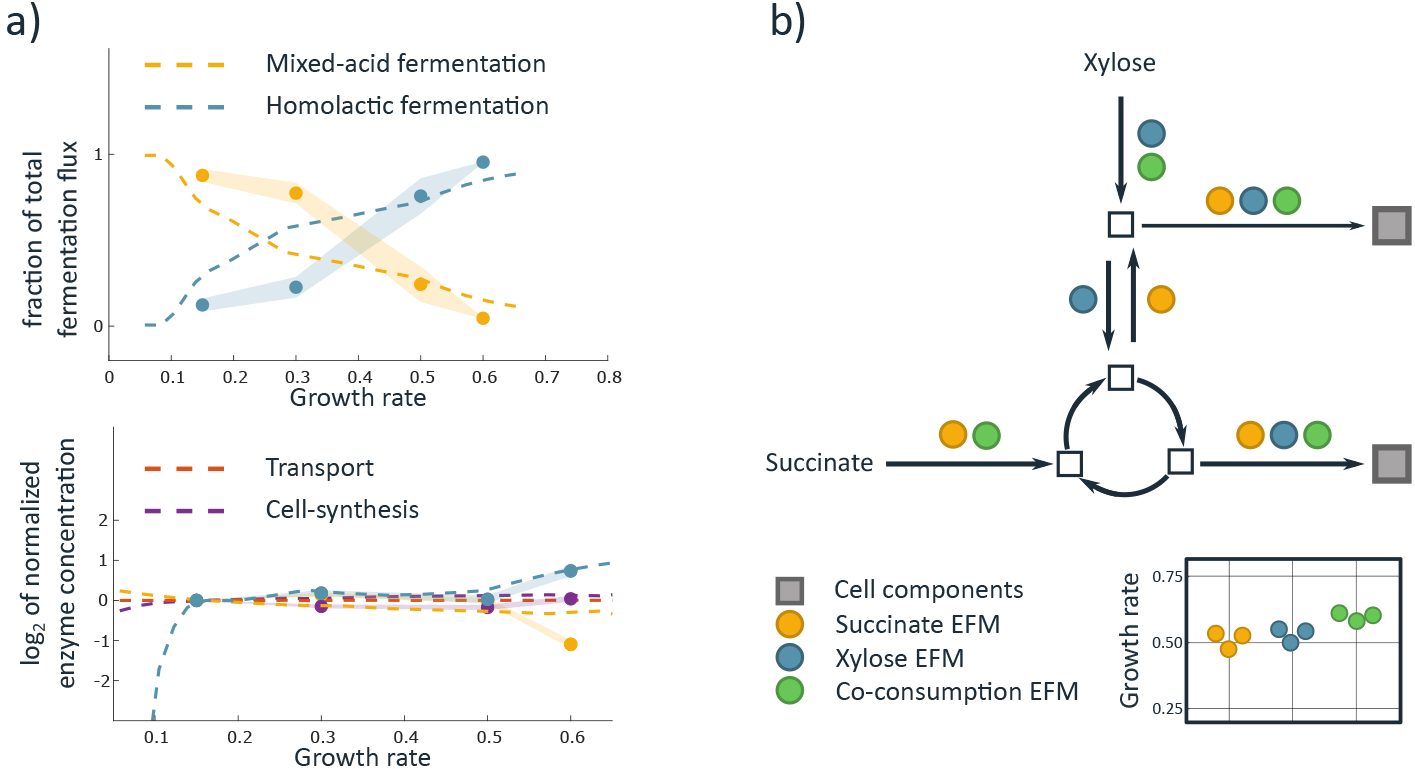
Under-utilization of enzymes and co-consumption can be understood with our kinetic, constrained-based approach. **a)** Model simulations of the metabolic switch of *L. lactis* are shown (dashed lines), along with experimental data from [44]. The flux predictions for both pathways are expressed as a fraction of the total flux through both pathways. Enzyme concentrations are normalized to the concentrations at a growth rate of 0.15 and then log-scaled. The model reproduces the switch from mixed-acid to homolactic fermentation at constant enzyme concentrations, because of its consideration of enzyme kinetics. Details of this model are described in S1 Appendix Section 10. To obtain a perfect fit with the data, a larger model should be invoked, but this is beyond the scope of this paper. We emphasize that protein concentrations can remain constant while pathway usage changes. **b)** An example is shown of a metabolic network with EFMs that use either succinate or xylose (orange and blue circles respectively), and an EFM (green circles) that uses two carbon sources. Grey squares denote products that are essential for cell growth. The co-consumption EFM can synthesize one cell component with succinate, and the other with xylose. The reaction that connects the upper and lower parts of the network therefore becomes inessential. This leads to a possible reduction in protein costs and therefore to a growth rate advantage. We indeed measured a growth rate increase by the co-consumption of succinate and xylose, as shown in the inset in which different biological replicates are indicated with different points. Results of the other combinations that were tested can be found in S1 Appendix Section 12.

In the presence of multiple carbon sources, microorganisms might consume them simultaneously [46–48]. We confirmed experimentally that *E. coli* only co-consumes carbon sources when this increases its growth rate (S1 Appendix Section 12). However, it is yet unclear why co-consumption can be favourable. Optimization models have been made that show simultaneous substrate uptake [47, 48], but the approach of Hermsen et al. [47] is mechanistic and does not provide a fundamental cause, and Beg et al. [48] state that “cells preferentially using the more efficient carbon source would outgrow those that allow the simultaneous utilization of other carbon sources”. Aidelberg et al. [46] state that single objective optimization approaches cannot explain co-consumption. However, we show that co-consuming EFMs (S1 Appendix Section 11) exist that reduce resource costs per unit growth rate, hence leading to higher growth rates. These new EFMs exist when each substrate makes a different set of precursors (see Figure 6b) for an illustration). Consequently, co-consumption can become favourable when reactions connecting a carbon source to a distant precursor are no longer essential. Following this reasoning, one would expect the largest growth benefit if substrates are co-consumed that enter the metabolic network far from each other. Indeed we, as well as others [47], observed the largest growth benefit when lower-glycolytic substrates are combined with upper-glycolytic substrates.

Some microbial strategies are seemingly growth rate reducing, such as the anticipatory expression of stress proteins [39] and alternative nutrient transporters [49], and the overcapacity of ribosomes [50]. That these strategies were still selected by evolution is often ascribed to fitness benefits in dynamic conditions. However, in our constraint-based approach these types of behaviours do not have to be growth rate reducing. Some of the protein pools might not be completely exploited, and the expression of proteins might then bring little or no costs. For example, our analysis of overflow metabolism shows that one of the constrained enzyme pools is underused at low growth rates. This underused pool can accommodate proteins that might be favourable for future conditions. For example, say that a microorganism faces a cytosolic and a membrane constraint, but suppose that only the membrane constraint is active at low growth rates. The unused cytosolic capacity can then be exploited for other purposes. The sole activity of a membrane constraint at low growth rates indeed explains why O’Brien et al. observed *E. coli* to have a ‘nutrient-limited’ [36] growth region at slow growth.

## Discussion

The extremum principle that we derived and illustrated in this work predicts the evolutionary direction on a short timescale, dictating optimal enzyme expression levels. At a given time, the extremum principle predicts that resources are reallocated to the most efficient enzymes at the expense of others that are less active per unit enzyme: evolution reduces the number of active EFMs. On a longer timescale, kinetic parameters and network stoichiometry can evolve, thereby changing the phenotypic potential: evolution modifies the cost vectors. In this new setting, the extremum principle will again predict minimal complexity, although the EFMs that are selected and the flux through these EFMs may have changed. Our theory predicts that a microorganism selected for maximal growth rate will, in static conditions, only express a small number of EFMs and therefore its metabolism is low-dimensional. This could very well be the explanation of the simple linear relations that many experimentally measured relations show [10–12]. This simplicity may also provide an explanation how only a few number of metabolites or proteins (‘‘master regulators” such as CcpA or Crp) seem to regulate (central) metabolism [51].

The insight that the dimensionality of metabolism is bounded by the number of active constraints is applicable to earlier modelling approaches that have used resource allocation principles. Furthermore, we show that the same principles also hold for nonlinear models that include enzyme kinetics and thereby metabolite dependencies. The kinetic self-replicator model presented by Molenaar et al. [5] for example, does not show mixed strategies, but an abrupt switch between respiration and fermentation, testimony of a single active constraint. Indeed, although a membrane protein constraint was included, the size of cells could be freely adjusted to alleviate this constraint. In many studies with genome-scale stoichiometric models, mixed strategies do occur. In all these studies the glucose uptake flux was constrained (first constraint), in combination with some linear combination of fluxes that reflects the (second) constraint that was the focus of the study (solvent capacity, osmotic pressure [19,52], proteome limits [4,42], membrane [20,40]). Also in so-called ME (Metabolism and Expression) models [36] and variants thereof [53], growth rate is fixed and nutrient uptake is minimized. Again, overflow is observed in these models when an additional constraint (total proteome) is hit.

Even though growth-rate maximization at constant conditions might at times be a rather crude approximation of the selective pressure, we expect the extremum principle to provide an ‘evolutionary arrow of time’. When conditions change frequently, other aspects might come into play and fitness will be captured by the mean growth rate over environments, i.e., the geometric growth rate [1]. Whether extremum principles hold for the maximization of geometric growth rate is an open problem for future theoretical work.

Even in static conditions, our theory is based on the assumption that a metabolic rate is maximized. In principle, this rate does not have to be the cell-synthesis rate, but could be another metabolic reaction. This might for example occur in case of specialization in multicellular organisms. However, we do not know if in these cases the selective pressure is strong enough to maximize this rate. Moreover, even microorganisms are not always optimally tuned, as it was shown that titration of ArcA could increase the growth rate of *E. coli* on glycolytic substrates significantly [54]. Indeed, the extremum principle does not describe metabolism if no rate is maximized, and our theory thus does not describe all suboptimal points in the fitness landscape. However, a principle that characterizes the peaks and shows the direction of increase at every point in a landscape, can still be of great guidance.

The success of constraint-based modeling methods suggests that indeed biophysical constraints shape microbial metabolism. However, most constraints used in the literature are postulated and remain unvalidated. Also, the imposed constraints can often not be directly deduced from the physiology of the microorganisms. Our theory suggests a mechanistic way forward for future constraint-based modeling methods. Our theory suggests that a constraint should be imposed for each cellular compartment with a limited solvent capacity for proteins. Since the number of compartments in prokaryotes is generally less than in eukaryotes, because they lack organelles, metabolic behaviour of prokaryotes is generally simpler.

Large-scale kinetic models are not yet used to study optimal metabolism. Growth rate maximization in such models quickly becomes computationally infeasible, because all metabolite and enzyme concentrations have to be tuned. Our results can offer some guidance in these large, nonlinear optimization problems. Say there are *K* constraints in the model, the extremum principle ensures that the optimum has to be found among conical combinations of *K* EFMs. This fact was already exploited in the case of one constraint in a medium-scale network [55]: EFMs could be optimized separately (which is a strictly convex problem [56]) and the one with the highest growth rate was picked. However, it is doubtful if this computational feasibility can be extended to models with more constraints. With two constraints all pairs of EFMs should already be considered and rate maximization in two EFMs under two constraints is not convex anymore.

The extremum principle is a null hypothesis about the course of a particular evolutionary process [57]. It has direct operational implications for evolutionary engineering strategies, when increasing or decreasing the complexity of microbial metabolism might be desired, for example in industrial biotechnology when co-consumption of different sugars from biomass-hydrolysates is pursued, or if prevention of overflow metabolism during heterologous protein production is attempted. Perhaps, when the growth-limiting constraints for the microorganism of interest have been identified, these could be perturbed to direct evolution in the preferred direction.

## Conclusion

Our theory suggests that metabolism has only a few operational degrees of freedom. By shifting perspective on rate maximization from the entire metabolic network to its representation in the cost vector formalism, we have reduced the problem to its essential dimensions, equal to the number of growth-limiting biophysical constraints. Together with the extremum principle, this work provides a species-overarching, molecular, constraint-based perspective on microbial metabolism.

## Supporting information

**S1 Appendix Theoretical derivations, mathematical proofs, core models, and a co-consumption experiment.**

**S1 Source Code Data Analysis Coconsumption Experiment.** All raw data and the Matlab-code used for data analysis can be found in the compressed folder attached to the supplements.

**S2 Source Code Kinetic model of overflow metabolism.** The Matlab-code used for modeling overflow metabolism is attached in a compressed folder as a supplement. In the compressed folder, we have also added a text-file with instructions.

**S3 Source Code Kinetic model of** ***L. lactis.*** The Matlab-code used for the kinetic model of *L. lactis* is attached in a compressed folder as a supplement. In the compressed folder, we have also added a text-file with instructions.

**S4 Source Code Finding coconsumption EFMs** The Python and Matlab-code used for finding co-consuming EFMs are attached in a compressed folder as a supplement. In the compressed folder, we have also added a text-file with instructions.

**S1 Dataset Growth rates co-consumption experiments.** `SI_growth_rates.txt` Estimated growth rates from separate biological replicates.

**S2 Dataset Substrate concentrations co-consumption experiments.** `SI_OD_conc_per_cond.xlsx` For all different growth media, we include an excell-sheet. Shown are the measured concentrations of carbon sources (normalized for initial concentration), with the corresponding Optical Density (OD). The letters that indicate the conditions denote the available carbon sources in the medium: S=Succinate, L=maLtose, M=Mannose, X=Xylose, G=Glucose.

**S3 Dataset Estimated uptake rates co-consumption experiments.** `SI_q_S_comp_cond.xlsx` Shown are the estimated uptake rates (mean and standard deviation) of different carbon sources (normalized for initial concentration) on the different growth media. The letters that indicate the conditions denote the available carbon sources in the medium: S=Succinate, L=maLtose, M=Mannose, X=Xylose, G=Glucose.

## Acknowledgments

We thank Sieze Douwenga, Willi Gottstein, Dennis Botman, Johan van Heerden, and Rinke van Tatenhove-Pel for comments, questions and proofreading. This work was supported by NWO VICI grant 865.14.005, by NWO STAR grant 022.005.031, and by Era-Industrial Biotechnology project nr. 053.80.772.

## Footnotes

1 In modelling methods that do not include kinetic information, such as FBA, it is much more probable for two EFMs to have the same costs. The optimal solutions in these modelling methods are therefore often multi-dimensional subspaces.

2 Note that in the proof of Wortel et al. the vector of metabolite concentrations ***x*** was fixed to its optimal value ***x***_opt_ before proceeding. This is not directly possible, since the optimal value is dependent on the choice of enzyme concentrations and these have yet to be determined. We use a small adaptation: we give an argument that works for all fixed ***x***, and therewith for the optimal ***x***.

3 Although experimentally the growth rate is set by the dilution rate of the glucose-limited chemostat, growth rate always correlates with the available glucose concentration.

4 Alternatively, a specific enzyme could be inhibited; this however introduces the risk of inhibiting some EFMs more than others, leaving the results potentially uninterpretable.

5 Technically, the inhibition of translation could possibly lengthen the cost vectors of all EFMs in the x-direction to different extents. We study this case separately in S1 Appendix Section 7 and show that the effects are equivalent to the effects of resizing the first enzyme pool.

## Notes

#### Summary of Updates

Revised such that the main theoretical result is more central to the paper.

