## Appendix 1 for "The number of active metabolic pathways is bounded by the number of cellular constraints at maximal metabolic rates"

#### Theoretical derivations, mathematical proofs, core models, and a co-consumption experiment

In this document all mathematical results are grouped and extensively described. In order to provide a document that can be read and understood on its own, we will not only describe our own results, but we will also describe and explain some of the prerequisite theory. This is also the reason that some of the theory below appears both in the main text and in this appendix.

### 1 Elementary Flux Modes, enzymatic constraints, and cost vectors

The structure of a metabolic network can be denoted by its stoichiometric matrix $\boldsymbol{N}$. Each column of this matrix denotes which metabolites are used (negative entries) and produced (positive entries) in one reaction. In a metabolic network with $m$ metabolites and $r$ reactions this results in an $m \times r$-matrix. A steady-state flux vector $\boldsymbol{v} \in \mathbb{R}^r$ must then be in the nullspace of this matrix, and thus satisfy $\boldsymbol{N} \cdot \boldsymbol{v} = 0$. Some reactions are reversible, but others are modelled as irreversible, $v_j \geq 0$. However, by splitting all reversible reactions into a forward and a backward reaction, we can consider networks with only irreversible reactions. From now on, we will thus only consider irreversible reactions. This causes the feasible flux vectors to form a polyhedral cone $\mathcal{P}$ (rather than a linear subspace) in so-called flux space [1].

For a flux vector, $\boldsymbol{v} \in \mathbb{R}^r$, its 'support' is the set of participating reactions $R(\boldsymbol{v}) = \{j : v_j \neq 0\}$. An elementary flux mode $\mathbf{EFM} = (V_1, \dots V_r)^T \in \mathcal{P}$ minimizes the number of elements in $R(\mathbf{EFM})$. In other words, if there is a $\boldsymbol{v}' \in \mathcal{P}$ such that $R(\boldsymbol{v}') \subseteq R(\mathbf{EFM})$ then we must have $\boldsymbol{v}' = \alpha\mathbf{EFM}$ for some $\alpha \neq 0$. Another way of phrasing this is that none of the used reactions can be set to zero in the EFM without violating the steady state condition.

Gagneur and Klamt showed that in any metabolic network, these EFMs coincide with the extreme rays of the pointed polyhedral cone $\mathcal{P}$ [1]. This implies that in a metabolic network every steady state flux vector $\boldsymbol{v}$ can be written as a conical combination of the EFMs [1]:

$$\boldsymbol{v} = \lambda_1 \mathbf{EFM}^1 + \ldots + \lambda_F \mathbf{EFM}^F, \text{ where } \lambda_i \geq 0, \tag{1}$$

where the multiplication factors $\lambda_i$ denote how much the $i^{\text{th}}$ EFM is used and $F$ denotes the total number of EFMs in the network. Note that, although the Elementary Flux Modes are constant vectors defined by stoichiometry, the $\lambda_i$-factors are variable and dependent on metabolite concentrations. We will make this dependence more precise in 5. Equation (1) shows that EFMs are the basic building blocks of steady state metabolism.

EFMs are defined up to a constant: if $\boldsymbol{v}$ is an EFM, than so is $\alpha\boldsymbol{v}$ for any $\alpha \in \mathbb{R}$. This has two important consequences. First, the ratio between flux entries in an EFM are fixed, and second, we may scale one entry of an EFM to 1. We will consider optimisation of some objective flux $v_r$ at steady state. Therefore, we only need to consider those EFMs which have a nonzero $r^{\text{th}}$ flux value. Without loss of generality, we can make this the last entry in the vector, and will scale the objective flux to 1 throughout this work, and will denote the $i^{\text{th}}$ EFM by $\mathbf{EFM}^i = (V_1^i, \dots, V_{r-1}^i, 1)^T \in \mathbb{R}^r$, with all $V_j^i$ uniquely determined by stoichiometry. Since the $V_r^i$ factors are all equal to one, the $\lambda_i$ factors in (1) may now be reinterpreted as the objective flux that $\mathbf{EFM}^i$ is contributing.

Individual metabolic reactions in the network are assumed to be catalyzed by enzymes, and reaction rates generally are well described by $v_j = k_{\mathrm{cat},j} e_j f_j(\boldsymbol{x})$, where $e_j$ denotes the concentration of enzyme $j$, and $f_j(\boldsymbol{x})$ denotes the reaction kinetics [2]. We will assume this type of dependence throughout the SI, and assume that each enzyme catalyzes one reaction. In some situations, we will distinguish between external concentrations $\boldsymbol{x}^E$ (which are assumed to be parameters), and internal concentrations $\boldsymbol{x}^I$ (which may be varied, for instance to obtain a maximal objective flux).

As we argued in the main text, the expression of enzymes will be subject to biophysical limits. We will model this by including $K$ linear enzymatic constraints of the form:

$$C_\Sigma^{(k)} := \sum_{j=1}^{r} w_j^{(k)} e_j \leq 1 \qquad \text{for } k \in \{1, \ldots, K\}.$$

#### The cost vectors: a low-dimensional view at metabolism

Given $K$ constraints, we can, for each EFM, calculate the cost per constraint for making one unit objective flux. These $K$ costs turn out to comprise all relevant information for growth rate optimisation. Therefore, we will here define the *cost vectors* that have these costs as their entries. We will use the cost vectors to study metabolism in low-dimensional *constraint space* throughout this paper.

As discussed above, we can rescale each EFM such that it is a vector of the form $\mathbf{EFM}^i = (V_1^i, \ldots, V_{r-1}^i, 1)^T \in \mathbb{R}^r$. To produce one unit objective flux, we thus need a flux of $V_j^i$ through reaction $j$. Since we have $v_j = k_{\mathrm{cat},j} e_j f_j(\boldsymbol{x})$, we get

$$e_j^i = \frac{V_j^i}{k_{\mathrm{cat},j} f_j(\boldsymbol{x})},$$

where $e_j^i$ denotes the necessary concentration of enzyme $j$ for one unit objective flux through EFM $i$. We can then define the *cost vector* $\boldsymbol{d}^i(\boldsymbol{x})$ for the $i^{\text{th}}$ EFM, with components given by the total costs that this EFM brings per constraint:

$$\begin{aligned} d_k^i(\boldsymbol{x}) &:= \sum_{j=1}^{r} w_j^{(k)} e_j^i, \\ &= \sum_{j=1}^{r} w_j^{(k)} \frac{V_j^i}{k_{\mathrm{cat},j} f_j(\boldsymbol{x})} \end{aligned} \tag{2}$$

Because enzyme kinetics determine the enzyme concentrations and thereby the enzymatic costs, it is unlikely that several EFMs have exactly the same costs. Different EFMs use at least one different enzyme, and it is highly improbable that the necessary concentrations of these different enzymes are exactly the same real number. If one of these non-overlapping enzymes is part of a constrained pool, the EFMs will thus have different costs.[1] If, however, none of the non-overlapping enzymes are part of the constrained pools, several EFMs can indeed have the same costs. To deal with this case we introduce the notion of *equivalent EFMs*.

[1] In modelling methods that do not include kinetic information, such as FBA, it is quite likely for two EFMs to have the same costs. The optimal solutions in these modelling methods are therefore often multi-dimensional subspaces.

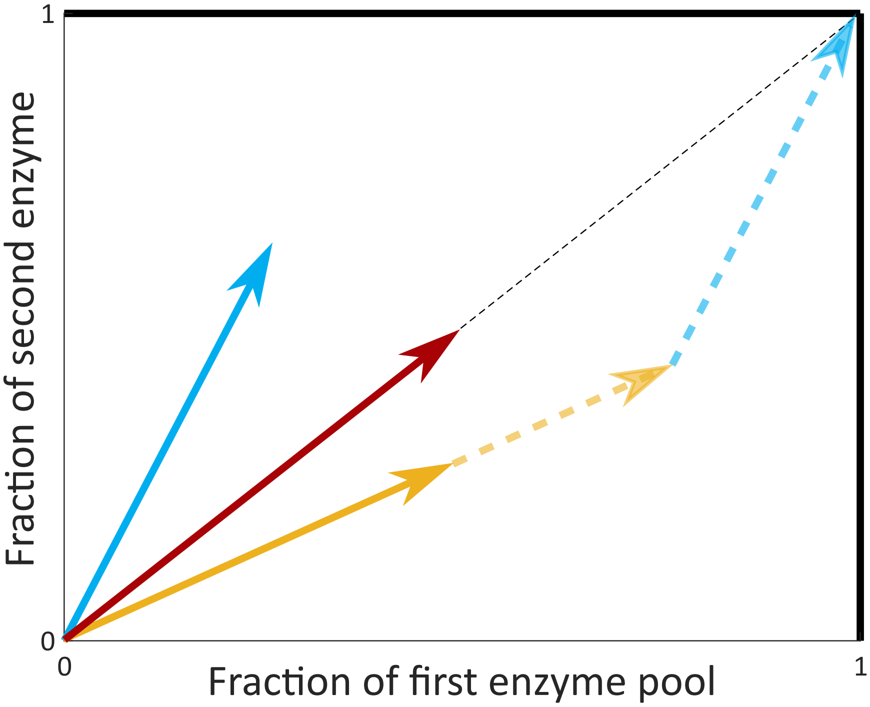


Figure 1: Cost vectors in the space of constraints. A cost vector denotes the fractions of the two limited enzyme pools that are needed to produce one unit of objective flux through this EFM. The objective flux is optimized by taking a sum of multiples $\lambda_1 \boldsymbol{d}^1 + \cdots + \lambda_m \boldsymbol{d}^M$ without leaving the box of constraints and while maximizing the sum of these multiples $\lambda_1 + \cdots + \lambda_m$. The optimal use of the EFMs is here indicated by the dashed lines.

**Definition 1.** *Given a set of constraints, $C_\Sigma^{(1)}, \ldots, C_\Sigma^{(K)}$, two EFMs, $\mathbf{EFM}^1, \mathbf{EFM}^2$, are called* equivalent with respect to the constraints *if their associated cost vectors are equal: $\boldsymbol{d}^1(\boldsymbol{x}) = \boldsymbol{d}^2(\boldsymbol{x})$.*

The definition of the cost vectors is illustrated in Figure 1.

### 2 Elementary Flux Modes are the optimal building blocks of metabolism

We here state the main result of this study, the extremum principle. Let us consider a metabolic network characterized by stoichiometry matrix $\boldsymbol{N}$. We ask which vector $\boldsymbol{v}$ in the steady state flux cone $\mathcal{P}$ maximizes a given objective flux $v_r$ under a given set of $K$ enzymatic constraints.

**Theorem 1.** *Consider a metabolic network characterized by $\boldsymbol{N}$. Let $v_r$ be an objective flux, which is to be maximized at steady state, under $K$ linear enzymatic constraints of the form:*

$$C_\Sigma^{(k)} := \sum_{j=1}^{r} w_j^{(k)} e_j \leq 1 \qquad \text{for } k \in \{1, \ldots, K\}.$$

*Then, at most $K$ non-equivalent Elementary Flux Modes are used in the optimal solution.*

*Proof.* The proof is shown in the main text. □

We think that the case where several EFMs are equivalent is not very common in biology. First, the constraints on enzyme expression are due to biophysical limits and we expect these to act on many enzymes together. This reduces the chance of having several EFMs that use exactly the same enzymes within the constrained pool of enzymes. Second, even if several EFMs would use the same enzymes, then the enzyme costs depend on the enzyme saturations, and these depend on the optimal metabolite concentrations. These optimal concentrations depend on the rest of metabolism, such that the non-overlapping part of the EFMs can still influence the enzyme costs. For these two reasons, we will assume in the rest of this work that EFMs are generally not equivalent.

The following corollary can be used to find out how many constraints are active when we observe a certain number of active EFMs. It is the contrapositive of Theorem 1 and therefore mathematically equivalent. The reason that it is stated separately is the difference in biological applicability: the theorem is a predictive statement while the corollary is descriptive.

**Corollary 2.** *If a flux $v_r$ is optimized and $M$ elementary flux modes are used, then at least $M$ linear enzymatic constraints must be active.*

### 3 The generality of enzyme constraints

Recall that we model growth optimisation as the maximisation of the cell synthesis flux ($v_r$). We assume that all reactions are catalyzed by an enzyme and that the reaction rate is proportional to the enzyme concentration (as long as the metabolite concentrations in the cell are kept constant), i.e. $v_i = k_{\text{cat},i} e_i f_i(\boldsymbol{x})$. Let us now pick a certain steady state flux distribution that leads to cell growth: $(v_1, \ldots, v_r)$. If the growth rate is limited, this means that there is a maximal value for $v_r$. However, without enzyme-concentration constraints, we could always increase the flux by adding enzymes in the right proportions: $(\frac{v_1}{f_1(\boldsymbol{x})}, \ldots, \frac{v_r}{f_r(\boldsymbol{x})})$. The fact that the growth rate is not infinite therefore either shows that enzyme constraints must be active, or that we are in some limit where $v_i = k_{\mathrm{cat},i} e_i f_i(\boldsymbol{x})$ is no longer true. This last possibility cannot be excluded, but we expect the enzyme constraints to have a more dominant effect.

Other growth-limiting constraints that have been studied, such as a limited solvent capacity of the (mitochondrial) membrane or the cytosol can generally be viewed as protein-concentration constraints. A limited membrane area results in a constrained pool of membrane proteins:

$$\sum_{i | e_i = \text{membrane}} w_i e_i \leq C,$$

where $w_i$ is the needed membrane area for one mol of the $i^{\text{th}}$ membrane protein and $C$ is the membrane area per cell volume.

Cytosolic volume can be written as

$$\sum_{i | e_i = \text{cytosol}} w_i e_i \leq C,$$

where $w_i$ is now the needed cytosolic volume for one mol $e_i$ and $C$ is the cytosolic volume per cell volume.

### 4 Most models of mixed behaviour are instances of the extremum principle

In this section we show that most existing modeling methods can be seen as an instance of the optimization problem that we studied here. This means that the optimal solutions are described by the extremum principle: the number of flux-carrying EFMs is bounded by the number of active constraints.

We here divide the existing models into the following classes:

a. **Genome-scale, classical FBA-models with only fluxes as variables and flux constraints.** These models omit enzyme kinetics and metabolite concentrations. When all reversible reactions are split into two irreversible reactions (as we have done before, 2), the steady-state assumption leads to a flux cone. This cone has extreme rays which are the Elementary Flux Modes of the metabolic network [1]. Under one flux constraint, this cone can become a bounded polyhedron. It follows from linear programming that an objective, which in this context is a weighted sum of fluxes, will be maximised in a vertex of the polyhedron, i.e. the flux is maximised in an EFM [3]. The addition of a new constraint will intersect the polyhedron to create a new polyhedron, with a new set of vertices. This set will contain some of the old vertices, as well as new vertices which are convex combinations of the old vertices. These new vertices are thus convex combinations of two EFMs. If the objective flux is maximal in such a new vertex, then two constraints are active. If however, the objective is optimised in an old vertex, then only one constraint is active and one EFM will be used. In any case, the number of active EFMs is bounded by the number of active constraints.

b. **Genome-scale and coarse-grained FBA-models with fluxes as variables and enzyme-concentration constraints [4–7].** The enzyme-concentration constraints can, at fixed metabolite concentrations and under the assumption that $v_i = k_{\mathrm{cat},i} e_i f_i(\boldsymbol{x})$, be transformed into linear flux constraints. Then we can treat this class of models similar to the models in class a..

c. **Coarse-grained kinetic models** [8]. These models obey the extremum principle, provided that, when we set the metabolite concentrations to their optimal values, the resulting optimisation problem (either in flux or enzyme space) is a linear program.

d. **ME-models with enzyme concentrations as variables and enzyme-concentration constraints.** Strictly speaking, ME-models do not fall under the class of models that obey the extremum principle. However, a similar principle exists for flux modes associated with ME-models, an unpublished finding by ourselves.

### 5 'Unfixing' the metabolite concentrations

Fixing the metabolite concentrations to an arbitrary $\boldsymbol{x_0}$ is a subtle step in the proof, since the outcome of the optimisation procedure depends on the choice of $\boldsymbol{x_0}$. Because of the fixed concentrations, the only quantities that influence the cell's metabolism are the enzyme concentrations. These can then be easily optimized. Although considering fixed concentrations helps in the proof, in biological applications we would often like to know how changing a specific external concentration influences a cell's optimal choice. For example, in a yeast cell growing on glucose we know that the cell has different strategies corresponding to different external glucose concentrations. It will use the efficient respiration strategy at low glucose levels, while it uses a combination of respiration and fermentation at higher levels.

As was argued in the proof of Theorem 1, we may rewrite the optimisation problem so that the $\boldsymbol{x}$-dependence is restricted to the cost vectors

$$d_k^i(\boldsymbol{x}) = \sum_{j=1}^{r} w_j^{(k)} \frac{V_j^i}{f_j(\boldsymbol{x})}.$$

This means that the general picture will always be as in 1, but that the cost vectors can be adjusted by the metabolite concentrations $\boldsymbol{x}$, see the shaded areas in 2. It is important to note that $\boldsymbol{x}$ consists of the concentrations of all metabolites that affect the metabolism in the network. As mentioned before, these can be divided into two groups: an external metabolite group $\boldsymbol{x^E}$ which can not be controlled by the cell and an internal metabolite group $\boldsymbol{x^I}$ which can be adjusted. The external concentrations are part of the optimization problem, while $\boldsymbol{x^I}$ is part of the solution.

Using the cost vector formalism, we can provide some insight into how the metabolite concentrations affect the arrows. Recall that maximizing the objective flux is equivalent to fitting the largest sum of multiples of cost vectors in the box of constraints. An important point to note is that a solution using $K$ elementary flux modes in the case of $K$ constraints is not as probable as one might suspect. There are two reasons for this.

First, internal metabolite concentrations $\boldsymbol{x^I}$ may be adjusted to use one EFM which satisfies more than one constraint simultaneously. In 1 this means that the cost vector of one EFM is pointed exactly diagonally, such as the red arrow. In the overflow metabolism examples, the diagonal arrow is the respiration cost vector. To what extent cost vectors can be adjusted by internal concentrations to become diagonal is largely dependent on the enzyme kinetics of the network.

Second, it could be suboptimal or not feasible to satisfy some constraint with equality. In this case there are $K-1$ active constraints and a maximum of $K-1$ Elementary Flux Modes will thus be chosen for any internal metabolite concentration. A cell will only start using an extra EFM when this new EFM can provide more flux by hitting the currently inactive constraint. This means that the enzyme usage of the currently active constraints per unit flux must be lower for the new EFM than for the current combination of EFMs. Otherwise, it is better not to hit the last constraint at all and keep using the $K-1$ EFMs.

What should be clear from the above discussion is that a mixed strategy is a very special situation in metabolism. The observation of a mixed strategy therefore provides much information about the number of constraints that act on the system. In the case of 2 EFMs we can summarize this in the following conditions, which we prove to be necessary and sufficient in Theorem 6. The conditions are illustrated in 2.

A cell that optimizes an objective flux uses a mixture of EFMs if and only if
**1)** there are at least two enzymatic constraints,
**2)** EFMs with off-diagonal cost vectors exist in the direction of both constraints,
**3)** each constraint has a different EFM that uses the smallest fraction of the corresponding enzyme pool,
**4)** it is not feasible to hit both constraints with a diagonal EFM for which the cost vector is in the triangle spanned by the two cost vectors.

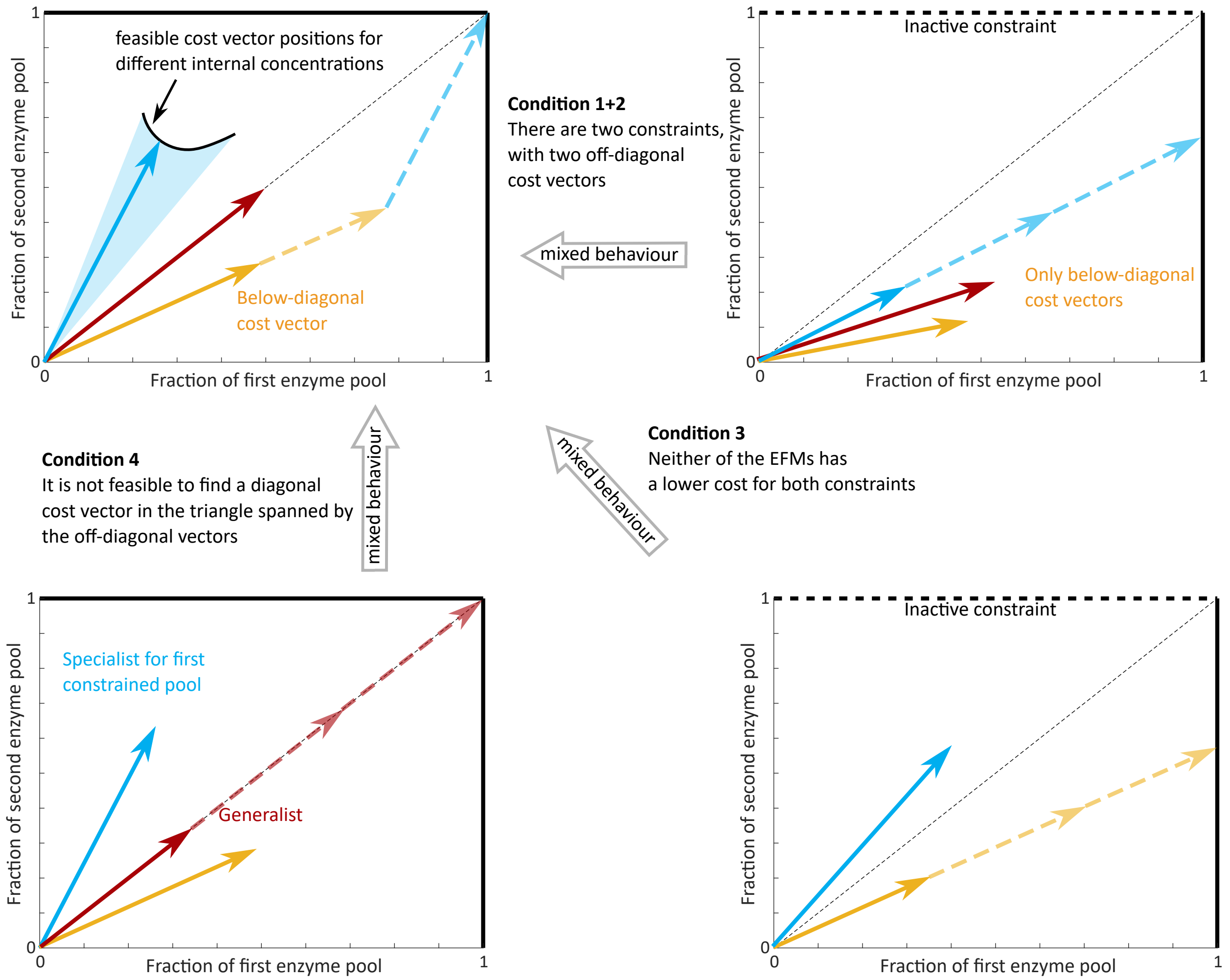


Figure 2: We distinguish four necessary and sufficient conditions for a mixture of EFMs to be optimal. The black line and blue shaded region in the left upper plot indicates the feasible cost vector positions at different internal concentrations. These positions were left out for the other cost vectors for simplicity, only the optimal cost vectors are shown there.

Before we prove this in a theorem we need some preparation in the form of a definition and two lemmas.

**Definition 2.** *Let two cost vectors* $\boldsymbol{d}^1$ *and* $\mathbf{d}^2$ *satisfy* $d_1^1 \leq d_2^1$, *and* $d_1^2 \geq d_2^2$. *They are said to be in a* mixable position *if* $d_1^1 > d_1^2$ *and* $d_2^1 < d_2^1$.

In many of the results that follow, the optimisation problem as phrased in the previous theorem, but now involving only two cost vectors, will be studied. For future reference, we state it here in full,

$$\max_{\boldsymbol{x}^I, \boldsymbol{\lambda}} \left\{ \sum \lambda_i \mid \lambda_i \geq 0, \; \boldsymbol{D}(\boldsymbol{x}) \cdot \boldsymbol{\lambda} \leq \mathbf{1} \right\}, \tag{3}$$

where $\boldsymbol{D} = \left[\boldsymbol{d}^1(\boldsymbol{x}), \boldsymbol{d}^2(\boldsymbol{x})\right]$ is the cost vector matrix.

**Lemma 3.** *Consider two cost vectors $\boldsymbol{d}^1(\boldsymbol{x})$, $\boldsymbol{d}^2(\boldsymbol{x})$ in mixable position, and consider Problem (3). Then the optimal solution will satisfy $\boldsymbol{D} \cdot \boldsymbol{\lambda} = [1, 1]^\intercal$.*

*Proof.* Let us say, for a contradiction that this is not true. Assume without loss of generality that $\boldsymbol{D} \cdot \boldsymbol{\lambda}^{\text{opt}} = [1, 1-c]^\intercal$. But then there is only one active constraint so only one EFM will be used in the optimal solution, let us say $\lambda_1^{\text{opt}} \boldsymbol{d}^1 = [1, 1-c]^\intercal$. This means that $d_1^1 > d_2^1$ and since the vectors are in mixable position we know that $d_1^2 \leq d_2^2, d_1^2 < d_1^1, d_2^2 > d_2^1$.

We can now find a better solution by combining the cost vectors. Choose $\epsilon = \frac{c}{2(d_2^2 - d_2^1)}$. Then consider the combination $(\lambda_1^{\text{opt}} - \epsilon)\boldsymbol{d}^1 + \epsilon \boldsymbol{d}^2$. We have

$$(\lambda_1^{\text{opt}} - \epsilon) d_1^1 + \epsilon d_1^2 = 1 - \frac{c}{2(d_2^2 - d_2^1)} d_1^1 + \frac{c}{2(d_2^2 - d_2^1)} d_1^2 = 1 - \frac{c(d_1^1 - d_1^2)}{2(d_2^2 - d_2^1)} < 1,$$

$$(\lambda_1^{\text{opt}} - \epsilon) d_2^1 + \epsilon d_2^2 = 1 - c - \frac{c}{2(d_2^2 - d_2^1)} d_2^1 + \frac{c}{2(d_2^2 - d_2^1)} d_2^2 = 1 - c + \frac{c(d_2^2 - d_2^1)}{2(d_2^2 - d_2^1)} = 1 - \frac{c}{2} < 1.$$

We thus hit none of the constraints while we obtain the same objective value $(\lambda_1^{\text{opt}} - \epsilon) + \epsilon = \lambda_1^{\text{opt}}$. Therefore we can increase either $\lambda_1$ or $\lambda_2$ which will give a higher objective value, in contradiction with the assumption that $\lambda_1^{\text{opt}}$ was the maximum. □

**Lemma 4.** *Consider Problem (3), and choose metabolite concentrations $\boldsymbol{x}$. Assume that $\boldsymbol{d}^1(\boldsymbol{x}), \boldsymbol{d}^2(\boldsymbol{x})$ are in mixable position. Then we have $\lambda_1^{opt}(\boldsymbol{x}) + \lambda_2^{opt}(\boldsymbol{x}) = \frac{1}{\beta}$, where $\beta$ is the intersection of the convex combination of the cost vectors and the diagonal of the constraint box, i.e.*

$$\beta \begin{bmatrix} 1 \\ 1 \end{bmatrix} = t \begin{bmatrix} d_1^1(\boldsymbol{x}) \\ d_2^1(\boldsymbol{x}) \end{bmatrix} + (1-t) \begin{bmatrix} d_1^2(\boldsymbol{x}) \\ d_2^2(\boldsymbol{x}) \end{bmatrix} \quad \textit{for suitable } t, \beta. \tag{4}$$

*Proof.* We suppress the dependence on $\boldsymbol{x}$ in the notation, as $\boldsymbol{x}$ is fixed throughout. Because the vectors are in mixable position, $\boldsymbol{D}$ is invertible, and Lemma 3 tells us that the optimal $\boldsymbol{\lambda}$ for these metabolite concentrations satisfies $\boldsymbol{D} \cdot \boldsymbol{\lambda}^{\text{opt}} = [\boldsymbol{d}^1, \boldsymbol{d}^2] \cdot \boldsymbol{\lambda}^{\text{opt}} = [1, 1]^\intercal$. Let $t$ and $\beta$ be chosen such that the convex combination of $\boldsymbol{d}^1$ and $\boldsymbol{d}^2$ lies on the diagonal, i.e., $t\boldsymbol{d}^1 + (1-t)\boldsymbol{d}^2 = \beta \boldsymbol{1}$. Then

$$\boldsymbol{D} \cdot \begin{bmatrix} t \\ 1-t \end{bmatrix} = \beta \boldsymbol{1}, \qquad \Longleftrightarrow \qquad \boldsymbol{D} \cdot \begin{bmatrix} \frac{t}{\beta} \\ \frac{1-t}{\beta} \end{bmatrix} = \boldsymbol{1}.$$

But also $\boldsymbol{D} \cdot \boldsymbol{\lambda}^{\text{opt}} = \boldsymbol{1}$, and we conclude

$$\boldsymbol{\lambda}^{\text{opt}} = \begin{bmatrix} \frac{t}{\beta} \\ \frac{1-t}{\beta} \end{bmatrix},$$

and

$$\lambda_1^{\text{opt}} + \lambda_2^{\text{opt}} = \frac{1}{\beta}.$$

□

This Lemma gives a very useful way to compare different pairs of EFMs in the constraint box. For each pair in mixable position, we determine the point where the convex combination of the pair intersects the diagonal. The pair of vectors whose intersection point lies closest to the origin has highest sum of $\lambda$'s, and gives rise to the highest flux.

**Corollary 5.** *A diagonal cost vector will give rise to a higher objective flux than a combination of two off-diagonal cost vectors if and only if the diagonal cost vector is contained in the triangle spanned by the off-diagonal vectors.*

**Theorem 6.** *Consider Problem (3) for any number of cost vectors. The global maximum will have at least two nonzero $\lambda$'s, say $\lambda_1, \lambda_2 > 0$, if and only if*

a. *there are at least two linear enzymatic constraints, i.e.* $\sum_{j=1}^{r} C_j^{(1)} e_j \leq 1$ *and* $\sum_{j=1}^{r} C_j^{(2)} e_j \leq 1$, *where the vectors* $\boldsymbol{C}^{(1)}$ *and* $\boldsymbol{C}^{(2)}$ *are linearly independent,*

b. *corresponding to each constraint there is an off-diagonal cost vector, i.e.* $d_1^i < d_2^i$ *and* $d_1^j > d_2^j$,

c. *each constraint has a different EFM with the smallest component in the direction of that constraint, i.e.* $d_1^i < d_1^j$ *and* $d_2^i > d_2^j$,

d. *we can not find an EFM with a diagonal cost vector* $\boldsymbol{d}^p$ *such that* $\boldsymbol{d}^p = \lambda_i d^i + \lambda_j d^j$, *with* $\lambda_i + \lambda_j \leq 1$.

*Proof.* We can prove the 'only if' part of this theorem by its contrapositive. Let us for each condition assume that it is not satisfied and show that this will result in the usage of fewer than 2 EFMs.

a. If there are no two constraints, Theorem 1 tells us that at most one EFM will be used, so only one $\lambda$ will be nonzero.

b. A combination of two EFMs will give a total objective flux of $\lambda_i + \lambda_j$, while it should satisfy $\lambda_i d_1^i + \lambda_j d_1^j \leq 1$ and $\lambda_i d_2^i + \lambda_j d_2^j \leq 1$. Let us say, without loss of generality, that there is no off-diagonal cost vector for the first constraint, which means that for all EFMs we have $d_1^i \geq d_2^i$. If both vectors are diagonal, the global maximum will only use the shortest vector (except for the negligible situation in which both diagonal vectors have the same length). So, let us assume that one of the vectors is off-diagonal $d_1^i < d_2^i$, because two diagonal vectors will never partake in a mixture of EFMs. Then the second constrained pool will never be fully used, since
$$\lambda_i d_2^i + \lambda_j d_2^j < \lambda_i d_1^i + \lambda_j d_1^j \leq 1.$$
But then there is in fact only one active constraint and Theorem 1 thus tells us that only one EFM will be used.

c. Let us say, without loss of generality, that $\mathbf{EFM}^i$ has the off-diagonal cost vector for constraint 1 and $\mathbf{EFM}^j$ for constraint 2. By definition $d_1^i < d_2^i$ and $d_2^j < d_1^j$, but to contradict the condition we also have $d_1^j \leq d_1^i$, which implies $d_2^j < d_2^i$ too, then
$$(\lambda_i + \lambda_j) d_k^i < \lambda_i d_k^i + \lambda_j d_k^j \leq 1$$
for both constraints. The feasible flux by using only $\mathbf{EFM}^i$ can therefore be larger than $\lambda_i + \lambda_j$ and only $\lambda_i$ will be greater than zero in the optimum.

d. If there was a diagonal cost vector in the triangle spanned by the off-diagonal cost vectors then the corresponding EFM would give rise to a higher flux than the combination of the off-diagonals by Lemma 4.

The 'if'-part of the theory remains. Assume for a contradiction that a pure EFM, $\mathbf{EFM}^p$ is used in the optimal solution. We know that there are two constraints, so this EFM could either hit both of these constraints or only hit one. In the first case, this EFM has a diagonal cost vector, but by conditions b. and d. there are then also two off-diagonal cost vectors that span a triangle which does not contain the diagonal cost vector. By Lemma 4, we know that this combination could then produce a higher flux than the pure EFM. In the case that the EFM hits only one constraint, say the first, the EFM has an off-diagonal cost vector, i.e. $d_1^p > d_2^p$. Pure usage of this EFM will not hit the second constraint, $\lambda_p d_2^p < 1$. Conditions b. and c. imply that there is another EFM, $\mathbf{EFM}^i$ in mixable position with $\mathbf{EFM}^p$. Pure usage of this $i^{\text{th}}$ EFM will not hit the first constraint, but Lemma 3 states that the maximum with these cost vectors will hit both constraints. Therefore, the maximum must have $\lambda_i, \lambda_p > 0$. This completes the proof. □

#### 6 The effect of tightening the constraints

An experimental perturbation can often be explained as affecting the constrained enzyme pools. Either one of the constrained enzyme pools is tightened, or the cost with respect to one of the enzyme pools for an EFM to make one unit flux is increased. These two types of perturbations are mathematically equivalent, so that from now on we will focus on the tightening of constraints. The situation in which the costs for one EFM is increased more than the cost for the other EFM will be discussed in 7. Here we show how different perturbations affect the gradual transition from usage of one EFM to a combination of two EFMs (as in overflow metabolism).

First we define the exact point of transition between one and two EFMs in the optimum.

**Definition 3.** *Consider Problem (3) and call the optimal flux solutions at certain environmental conditions $\lambda^{opt}(\boldsymbol{x}^E)$. The* critical conditions *are external environmental conditions $\boldsymbol{x}^E_{cr} \in \mathbb{R}^e$, for which there is a continuous map $p(\tau) : [-1, 1] \to \mathbb{R}^e$ with $p(0) = x^E_{cr}$ and such that*

$$\lambda_2^{opt}(p(\tau)) = 0 \quad \textit{for} \quad \tau \leq 0,$$
$$\lambda_1^{opt}(p(\tau)), \lambda_2^{opt}(p(\tau)) > 0 \quad \textit{for} \quad \tau > 0.$$

*Moreover, we define* gradual critical conditions *if $\boldsymbol{\lambda}^{opt}(p(\tau))$ depend continuously on $\tau$.*

Tightening the constraint that limits the active EFM (let us call it the first constraint) most, will force the growth-maximising micro-organism to switch already at these conditions. This is proven in full generality for any flux-maximising metabolic network in Subsection 6.1 below. Tightening the other constraint will, however, delay the switch. In many experiments these 'critical conditions' correspond to a concentration of a growth-limiting substrate. This critical concentration at the switch will decrease (resp. increase) if the first (resp. second) constraint is tightened. Therefore this is the first quantitative marker that can be used to determine which enzyme pool is affected most by the perturbation.

At non-critical environmental conditions we can still draw some conclusions. If both constrained enzyme pools are tightened equally by the experimental perturbation, we know exactly what happens to points on the flux-versus-growth rate plot. A growth-maximising micro-organism will keep the same ratio of its fluxes versus its growth rate, although both decrease due to the perturbation, see Theorem 11.

It is more difficult to understand fully what happens when we are at non-critical conditions and the perturbation affects one pool more than the other. However, note that biological data is often well-described by models in which enzyme saturations are kept fixed. This is equivalent to the assumption that internal metabolite concentrations vary only mildly over different growth conditions. Fixing these saturations would for example give rise to straight acetate excretion lines in overflow metabolism, exactly as found in the literature [5]. We therefore use this assumption to make a qualitative prediction of how experimental perturbations affect points on the fermentation flux curve, while further environmental conditions are kept fixed. Tightening the constraint that bounds respiration most (the first constraint) will decrease the growth rate more than it will decrease the fermentation flux. Tightening the other constraint will decrease fermentation more than the growth rate, and tightening both constraints equally will decrease fermentation and growth rate proportionally. We give the details in Subsection 6.2.

##### 6.1 The critical substrate concentrations are affected differently for tightening the different constraints

In the next Theorem we will focus on the metabolic switch from a pure strategy to a mixed strategy. We pick the exact environmental conditions at which the microorganism switches from one to two EFMs, see Definition 3, and then tighten one of the constraints. Tightening the constraint that bounds the active EFM more than the inactive EFM (in the overflow metabolism examples this is the first constraint) will force the organism to choose the mixed strategy already at these conditions. Tightening the other constraint will postpone the mixing of EFMs.

**Theorem 7.** *Consider again the following maximisation problem*

$$\max_{\boldsymbol{x}^I,\boldsymbol{\lambda}} \left\{ \lambda_1 + \lambda_2 \mid \lambda_i \geq 0,\ \boldsymbol{D}(\boldsymbol{x}^E, \boldsymbol{x}^I) \cdot \boldsymbol{\lambda} \leq \begin{bmatrix} 1 \\ 1 \end{bmatrix} \right\}, \tag{5}$$

*where* $\boldsymbol{D} = \left[\boldsymbol{d}^1(\boldsymbol{x}^E, \boldsymbol{x}^I),\ \ \boldsymbol{d}^2(\boldsymbol{x}^E, \boldsymbol{x}^I)\right]$ *is the cost vector matrix. Assume that the cost vectors are in mixable position, that* $\boldsymbol{x}^E$ *is such that the global maximum has* $\lambda_2 = 0$ *and additionally that at this optimum* $\frac{\mathrm{d}(\lambda_1+\lambda_2)}{\mathrm{d}x^i} = 0$ *for each* $i \in I$*. Then the maximisation problem*

$$\max_{\boldsymbol{x}^I,\boldsymbol{\lambda}} \left\{ \lambda_1 + \lambda_2 \mid \lambda_i \geq 0,\ \boldsymbol{D}(\boldsymbol{x}^E, \boldsymbol{x}^I) \cdot \boldsymbol{\lambda} \leq \begin{bmatrix} 1-\nu \\ 1 \end{bmatrix} \right\}, \tag{6}$$

*with* $\nu > 0$ *has a global maximum at* $\lambda_2 > 0$*.*

*Proof.* For given external conditions $\boldsymbol{x}^E$ we have a set of optimal internal concentrations $\boldsymbol{x}^I$ and an optimal set of fluxes $\boldsymbol{\lambda}$. Let us call the initial external concentrations $\boldsymbol{x}_0^E$ and the corresponding optima $\boldsymbol{x}_0^I$ and $\boldsymbol{\lambda}(\boldsymbol{x}_0)$. We will keep $\boldsymbol{x}_0^E$ fixed throughout, and are interested in the change of the optimal $\boldsymbol{\lambda}$ under tightening constraints. By assumption, $\lambda_2(\boldsymbol{x}_0) = 0$ and $\left.\frac{\mathrm{d}(\lambda_1(\boldsymbol{x}_0)+\lambda_2(\boldsymbol{x}_0))}{\mathrm{d}x^i}\right|_{\boldsymbol{x}_0} = 0$ for $i \in I$.

To exploit our knowledge about this zero derivative, we must use that the two-component cost vectors $\boldsymbol{d}^1, \boldsymbol{d}^2$ depend smoothly on all internal metabolite concentrations $(x_1, \ldots x_n)$ in a neighbourhood of $\boldsymbol{x}_0^I$. The dependence on $\boldsymbol{x}$ in the cost vectors appears in the rate kinetics functions $f_j(\boldsymbol{x})$. Since these depend smoothly on $\boldsymbol{x}$, so do the cost vectors.

Let $\boldsymbol{\lambda} = \boldsymbol{\lambda}(\boldsymbol{x}_0)$ denote the solution to $\boldsymbol{D}(\boldsymbol{x}_0) \cdot \boldsymbol{\lambda} = \mathbf{1}$. Solving this explicitly for $\lambda_1$ and $\lambda_2$ yields

$$\lambda_1(\boldsymbol{x}_0) = \frac{d_2^2(\boldsymbol{x}_0) - d_1^2(\boldsymbol{x}_0)}{\det \boldsymbol{D}}, \quad \lambda_2(\boldsymbol{x}_0) = \frac{-d_2^1(\boldsymbol{x}_0) + d_1^1(\boldsymbol{x}_0)}{\det \boldsymbol{D}}.$$

Using that $\boldsymbol{d}^1$ is a diagonal vector, we find for $i \in I$ that

$$0 = \frac{\mathrm{d}}{\mathrm{d}x_i}\left(\lambda_1(\boldsymbol{x}_0) + \lambda_2(\boldsymbol{x}_0)\right) = \frac{\frac{\partial d_1^1(\boldsymbol{x}_0)}{\partial x_i} + \frac{\partial d_2^1(\boldsymbol{x}_0)}{\partial x_i}}{\det \boldsymbol{D}}.$$

Recognizing this as a dot product, we find that $\frac{\partial}{\partial x_i}\boldsymbol{d}^1(\boldsymbol{x}_0) \perp \begin{bmatrix} d_2^2 - d_1^1 \\ d_1^1 - d_1^2 \end{bmatrix} \perp \boldsymbol{d}^1 - \boldsymbol{d}^2$, where we again used that $\boldsymbol{d}^1$ is diagonal. Because we do this analysis in two dimensions, we get

$$\frac{\partial}{\partial x_i}\boldsymbol{d}^1(\boldsymbol{x}_0) = c\left(\boldsymbol{d}^2(\boldsymbol{x}_0) - \boldsymbol{d}^1(\boldsymbol{x}_0)\right) \tag{7}$$

for a suitable nonzero $c \in \mathbb{R}$. We conclude therefore that the derivative of the first cost vector with respect to *all possible* changes in the internal metabolite concentrations points in the direction of the second cost vector, see 3.

Expanding $d_1^1$ around $\boldsymbol{x}_0$,

$$d_1^1(\boldsymbol{x}) = d_1^1(\boldsymbol{x}_0) + \left[(\boldsymbol{x} - \boldsymbol{x}_0) \cdot \nabla_{\boldsymbol{x}} d_1^1(\boldsymbol{x}_0)\right] + \left[(\boldsymbol{x} - \boldsymbol{x}_0) \cdot (H(d_1^1)(\boldsymbol{x}_0) \cdot (\boldsymbol{x} - \boldsymbol{x}_0))\right] + \cdots, \tag{8}$$

where

$$H(d_1^1) = \begin{bmatrix} \frac{\partial^2 d_1^1}{\partial x_1^2} & \cdots & \frac{\partial^2 d_1^1}{\partial x_1 \partial x_m} \\ \vdots & \ddots & \vdots \\ \frac{\partial^2 d_1^1}{\partial x_m \partial x_1} & \cdots & \frac{\partial^2 d_1^1}{\partial x_m^2} \end{bmatrix}. \tag{9}$$

Using (7), and setting $\boldsymbol{x} = \boldsymbol{x}_0 + \boldsymbol{\delta x}$, with $\boldsymbol{\delta x}^{\boldsymbol{E}} = \boldsymbol{0}$ (i.e, we only vary the internal concentrations), we get

$$\boldsymbol{d}^1(\boldsymbol{x}) = \boldsymbol{d}^1(\boldsymbol{x}_0) + \left[c\boldsymbol{\delta x} \cdot \left(\boldsymbol{d}^2(\boldsymbol{x}_0) - \boldsymbol{d}^1(\boldsymbol{x}_0)\right)\right] + \left[\boldsymbol{\delta x} \cdot (H(\boldsymbol{d}^1)(\boldsymbol{x}_0) \cdot \boldsymbol{\delta x})\right] + \cdots, \tag{10}$$

where we have used slightly abusive notation for the vector-valued Hessian:

$$H(\boldsymbol{d}^1) = \begin{bmatrix} \frac{\partial^2 \boldsymbol{d}^1}{\partial x_1^2} & \cdots & \frac{\partial^2 \boldsymbol{d}^1}{\partial x_1 \partial x_m} \\ \vdots & \ddots & \vdots \\ \frac{\partial^2 \boldsymbol{d}^1}{\partial x_m \partial x_1} & \cdots & \frac{\partial^2 \boldsymbol{d}^1}{\partial x_m^2} \end{bmatrix}.$$

Equation (31) describes the feasible positions of the first cost vector under different internal metabolite concentrations.

Let us now consider the new maximisation problem, given by Equation (6). This gives a new diagonal for the constraint box $s[1-\nu, 1]^T$ with $s \in [0, 1]$. Note that the generalisations of Definition 2 and Lemmas 3 and 4 remain valid, because we can simply rescale the new constraints and cost vectors to get the unit box again. There are now two options for the new maximum of $\lambda_1 + \lambda_2$, either $\lambda_2 = 0$ or $\lambda_2 > 0$. According to Theorem 6 the first option ($\lambda_2 = 0$) will only be optimal if there is a new set of metabolite concentrations for which $\boldsymbol{d}^1$ is on the new diagonal. Therefore we distinguish two strategies to find a global optimum: 1) adjust the metabolite concentrations such that the first cost vector will be on the new diagonal and 2) take a combination of the cost vectors to end up on the new diagonal. For option 2) we at least know one feasible combination of cost vector positions, which are the old positions $\boldsymbol{d}^1(\boldsymbol{x}_0)$ and $\boldsymbol{d}^2(\boldsymbol{x}_0)$. The intersection of this combination of two cost vectors with the new diagonal will be along the line

$$\boldsymbol{d}^1(\boldsymbol{x}_0) + m(\boldsymbol{d}^2(\boldsymbol{x}_0) - \boldsymbol{d}^1(\boldsymbol{x}_0)) \qquad m \in [0,1]. \tag{11}$$

Option 1) is a little bit more complicated because it involves the Hessian $H(\boldsymbol{d}^1)$. It was recently shown that for practically any rate law, sums of the form $\sum_k \frac{c_k}{f_k(\boldsymbol{x})}$ are strictly convex in the optimum [9]. The cost vectors $\boldsymbol{d}^1(\boldsymbol{x})$ and $\boldsymbol{d}^2(\boldsymbol{x})$ have exactly this form. Therefore, for each component of the cost vector, the Hessian is strictly positive definite: $\boldsymbol{\delta x} \cdot (H(d^1_k)(\boldsymbol{x}_0) \cdot \boldsymbol{\delta x}) > 0$.

We will now use this result to prove that the intersection obtained by option 1) is always further from the origin than the intersection obtained by option 2), thereby proving that the new global maximum will have $\lambda_2 > 0$. In order to do this we define a linear map $\omega$ that assigns a real number to the cost vector $\boldsymbol{d}^1(\boldsymbol{x})$. This number is equal to the length of the projection of the cost vector to a vector perpendicular to $\boldsymbol{d}^2(\boldsymbol{x}_0) - \boldsymbol{d}^1(\boldsymbol{x}_0)$. We can write this down as

$$\omega\left(\begin{bmatrix} a \\ b \end{bmatrix}\right) = \begin{bmatrix} d^2_2(\boldsymbol{x}_0) - d^1_2(\boldsymbol{x}_0) \\ d^1_1(\boldsymbol{x}_0) - d^2_1(\boldsymbol{x}_0) \end{bmatrix} \cdot \begin{bmatrix} a \\ b \end{bmatrix} =: \begin{bmatrix} v \\ w \end{bmatrix} \cdot \begin{bmatrix} a \\ b \end{bmatrix}, \tag{12}$$

for which it is only important that $v$ and $w$ are positive. This is true because the vectors are in mixable position. This linear map is equal-valued for all the points on the line between the two original cost vectors:

$$\omega\left(\boldsymbol{d}^1(\boldsymbol{x}_0) + m(\boldsymbol{d}^2(\boldsymbol{x}_0) - \boldsymbol{d}^1(\boldsymbol{x}_0))\right) = \begin{bmatrix} v \\ w \end{bmatrix} \cdot \begin{bmatrix} d^1_1(\boldsymbol{x}_0) \\ d^1_2(\boldsymbol{x}_0) \end{bmatrix} = v d^1_1(\boldsymbol{x}_0) + w d^1_2(\boldsymbol{x}_0). \tag{13}$$

The linear map applied to the feasible positions of $\boldsymbol{d}^1$ (Equation (31)) yields

$$\begin{aligned} \omega\left(\boldsymbol{d}^1(\boldsymbol{x})\right) &= v d^1_1(\boldsymbol{x}_0) + w d^1_2(\boldsymbol{x}_0) + \begin{bmatrix} v \\ w \end{bmatrix} \cdot \left[\boldsymbol{\delta x} \cdot (H(\boldsymbol{d}^1)(\boldsymbol{x}_0) \cdot \boldsymbol{\delta x})\right] + \cdots, \\ &= v d^1_1(\boldsymbol{x}_0) + w d^1_2(\boldsymbol{x}_0) + \left[\boldsymbol{\delta x} \cdot (H(v d^1_1 + w d^1_2)(\boldsymbol{x}_0) \cdot \boldsymbol{\delta x})\right] + \cdots. \end{aligned}$$

Since $v, w > 0$, the function $v d^1_1(\boldsymbol{x}) + w d^1_2(\boldsymbol{x})$ is still strictly convex near its optimum and it has an optimum exactly at $\boldsymbol{x}_0$, since

$$\begin{aligned} \frac{\partial}{\partial x_i}\left(v d^1_1(\boldsymbol{x}) + w d^1_2(\boldsymbol{x})\right)\Big|_{\boldsymbol{x}=\boldsymbol{x}_0} &= \frac{\partial}{\partial x_i}\left(\left(d^2_2(\boldsymbol{x}_0) - d^1_2(\boldsymbol{x}_0)\right) d^1_1(\boldsymbol{x}) + \left(d^1_1(\boldsymbol{x}_0) - d^2_1(\boldsymbol{x}_0)\right) d^1_2(\boldsymbol{x})\right), \\ &= \begin{bmatrix} d^2_2(\boldsymbol{x}_0) - d^1_2(\boldsymbol{x}_0) \\ d^1_1(\boldsymbol{x}_0) - d^2_1(\boldsymbol{x}_0) \end{bmatrix} \cdot \frac{\partial}{\partial x_i}\boldsymbol{d}^1(\boldsymbol{x}) \\ &= \begin{bmatrix} d^2_2(\boldsymbol{x}_0) - d^1_2(\boldsymbol{x}_0) \\ d^1_1(\boldsymbol{x}_0) - d^2_1(\boldsymbol{x}_0) \end{bmatrix} \cdot \left(\boldsymbol{d}^2(\boldsymbol{x}_0) - \boldsymbol{d}^1(\boldsymbol{x}_0)\right) \\ &= 0. \end{aligned}$$

This strict convexity means that the extra term $(\boldsymbol{\delta x} \cdot (H(v d^1_1 + w d^1_2)(\boldsymbol{x}_0) \cdot \boldsymbol{\delta x}))$ is positive, so we get the following inequality for the intersection points with the diagonal for the pure and the mixed strategy

$$\omega\left(\boldsymbol{d}^1(\boldsymbol{x})\right) > \omega\left(\boldsymbol{d}^1(\boldsymbol{x}_0) + m(\boldsymbol{d}^2(\boldsymbol{x}_0) - \boldsymbol{d}^1(\boldsymbol{x}_0))\right). \tag{14}$$

To be able to determine which of the two strategies, with $\lambda_2 = 0$ or $\lambda_2 > 0$ yields higher fitness, we invoke Corollary 5 and study the points of intersection with the new diagonal $[1-\nu, 1]^\intercal$. The strategy with intersection closest to the origin then yields highest flux. The intersection point of a strategy with value $\lambda_2 = \lambda_2^*$ is $s[1-\nu, 1]^\intercal$ for a suitable $s > 0$, and the $s$-value indicated by $s_{\lambda_2=\lambda_2^*}$.

We can now easily show that $s_{\lambda_2>0} > s_{\lambda_2=0}$, so that we will have $\lambda_2 > 0$ in the global maximum for the new maximisation problem. This follows from

$$s_{\lambda_2=0}\left((1-\nu)v + w\right) = \left(\omega\left(s_{\lambda_2=0}(1-\nu, 1)\right) = \omega\left(\boldsymbol{d}^1(\boldsymbol{x})\right) > \omega\left(\boldsymbol{d}^1(\boldsymbol{x}_0) + m(\boldsymbol{d}^2(\boldsymbol{x}_0) - \boldsymbol{d}^1(\boldsymbol{x}_0))\right)\right.$$
$$= \omega\left(s_{\lambda_2>0}(1-\nu, 1)\right) = s_{\lambda_2>0}\left((1-\nu)v + w\right),$$

so that $s_{\lambda_2=0} > s_{\lambda_2>0}$. Combined with Lemma 4 this implies that tightening the first constraint will lead to a combination at environmental conditions where a pure strategy was first used. □

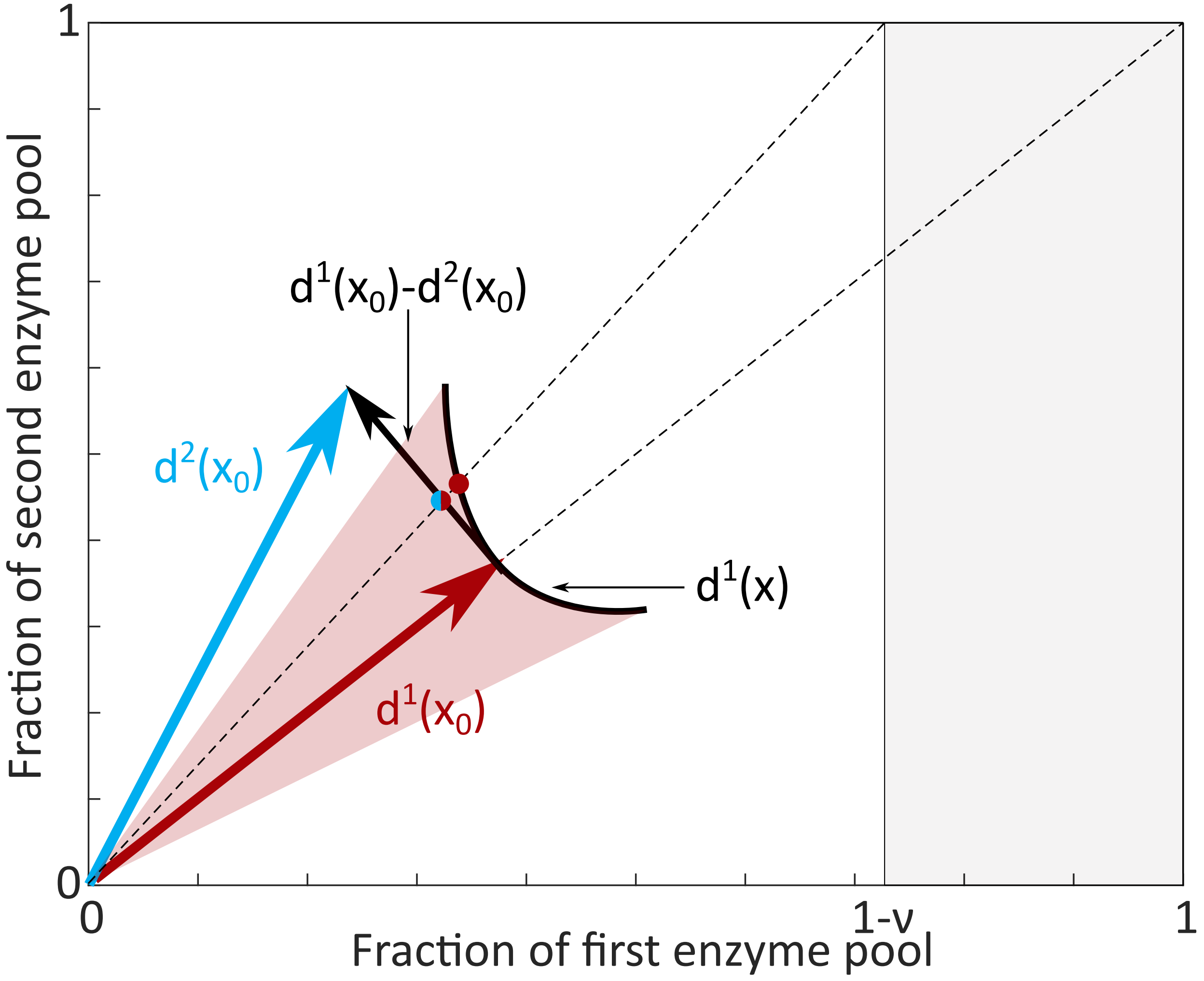

Figure 3: A visualisation of the proof of Theorem 7. At the critical environmental conditions, the derivative of cost vector $\boldsymbol{d}^1(x)$ with respect to the metabolite concentrations is in the direction of $\boldsymbol{d}^2(x)$. Moreover, the curve of feasible positions for $\boldsymbol{d}^1(x)$ lies completely outside the triangle spanned by the two cost vectors. This means that upon tightening of the first constraint, and thereby changing the diagonal of the constraint box, a mixture of EFMs will give rise to a higher flux than pure use of the red EFM.

**Corollary 8.** *Assume that we choose the gradual critical environmental conditions (following Definition 3). Experimentally tightening the constraint that limits the active EFM will force the microorganism to mix EFMs already at these conditions.*

*Proof.* According to the proof of Theorem 1, a growth maximising wildtype microorganism will solve the following maximisation problem

$$\max_{\boldsymbol{x}^I, \boldsymbol{\lambda}} \left\{ \lambda_1 + \lambda_2 \mid \lambda_i \geq 0,\ \boldsymbol{D}(\boldsymbol{x}^E, \boldsymbol{x}^I) \cdot \boldsymbol{\lambda} \leq \begin{bmatrix} 1 \\ 1 \end{bmatrix} \right\}, \tag{15}$$

where $\boldsymbol{D} = \begin{bmatrix} \boldsymbol{d}^1(\boldsymbol{x}^E, \boldsymbol{x}^I) & \boldsymbol{d}^2(\boldsymbol{x}^E, \boldsymbol{x}^I) \end{bmatrix}$ is the cost vector matrix. The definition of gradual critical conditions (Definition 3) states that there is a path $p(\tau)$ with $p(0) = x^E_{cr}$ that leads from pure EFM usage $\lambda_2 = 0$ to mixed EFM usage $\lambda_2 > 0$. We will here prove that we are then in the situation to which Theorem 7 applies.

We start by showing that the optimal internal concentration $\boldsymbol{x}^I$ for vectors in mixable positions depend continuous differentiable on the external concentrations $\boldsymbol{x}^E$. Recall that these optima are calculated by setting the derivatives to zero of the objective function $\lambda_1 + \lambda_2$ with respect to all internal concentrations. By Lemma 4 we know that this is equivalent to setting the derivatives of $\beta(\boldsymbol{x}^E, \boldsymbol{x}^I)$ to zero, where $\beta$ is determined by $t(\boldsymbol{x}^E, \boldsymbol{x}^I)\boldsymbol{d}^1(\boldsymbol{x}^E, \boldsymbol{x}^I) + (1 - t(\boldsymbol{x}^E, \boldsymbol{x}^I))\boldsymbol{d}^2(\boldsymbol{x}^E, \boldsymbol{x}^I) = [\beta(\boldsymbol{x}^E, \boldsymbol{x}^I), \beta(\boldsymbol{x}^E, \boldsymbol{x}^I)]^T$. In the optimum we have

$$\nabla_{\boldsymbol{x}^I} \beta(\boldsymbol{x}^E, \boldsymbol{x}^I) = \boldsymbol{0}. \tag{16}$$

We can now apply the Implicit Function Theorem (IFT) if the Hessian $H(t\boldsymbol{d}^1 + (1-t))\boldsymbol{d}^2)$ is invertible. Since the Hessian is positive definite [9], it is also invertible. The IFT then states that there are continuous differentiable functions around the optimum that describe the optimal internal concentrations as a function of the external concentrations, $\boldsymbol{x}^I(\boldsymbol{x}^E)$.

Since the cost vectors are smooth functions of the internal and external concentrations, we may define a continuously differentiable map $\bar{\boldsymbol{d}}$ of $\tau$ by setting

$$\bar{\boldsymbol{d}}(\tau) := \boldsymbol{d}(p(\tau), \boldsymbol{x}^I(p(\tau))).$$

We also know that for all $\tau > 0$ we have a mixed strategy, i.e. $\lambda_2 > 0, \lambda_1 > 0$. This will only be optimal when the vectors are in mixable position (Theorem 6). Since the vectors in mixable position form a set in $\mathbb{R}^2$. Since the continuous map $\bar{\boldsymbol{d}}(\tau)$ maps the open interval $(0, 1)$ in this closed set, we know that it will map the limit point $\tau = 0$ into this set or its boundary. The boundary of the set consists of those combinations of vectors for which $d_1^1 = d_1^2$ or $d_2^1 = d_2^2$. In that case, there will be no global optimum, so that the assumption of gradual critical environmental conditions would be violated. Hence, the vectors at $\tau = 0$ must still be in mixable position.

Since the objective function $\lambda_1 + \lambda_2$ is also a continuous differentiable function of the concentrations, these will also vary smoothly with $\boldsymbol{x}^E$. This gives us a continuous differentiable map $(\lambda_1 + \lambda_2)(p(\tau))$ from the path $p$.

For all $\tau > 0$ we know that $(\lambda_1 + \lambda_2)(p(\tau))$ is the optimal flux value, and $\nabla_{\boldsymbol{x}^I}(\lambda_1 + \lambda_2)|_{\tau>0} = 0$. So, the continuous map $\nabla_{\boldsymbol{x}^I}(\lambda_1 + \lambda_2)(\tau) = 0$ is zero throughout the open interval $(0, 1)$, but therefore also has to be zero on the limit point.

We conclude that at $\tau = 0$, the cost vectors are in mixable position, $\lambda_2 = 0$ is a global optimum (by definition of critical), and $\nabla_{\boldsymbol{x}^I}(\lambda_1 + \lambda_2)|_{\tau=0} = 0$. We therefore are in the exact situation to which Theorem 7 applies. Experimentally tightening the constraint that inhibits the active EFM $\lambda_1$ most is now equivalent to changing the constraint in the maximisation problem to $\boldsymbol{D}(\boldsymbol{x}^E, \boldsymbol{x}^I) \cdot \boldsymbol{\lambda} \leq \begin{bmatrix} 1-\nu \\ 1 \end{bmatrix}$. The theorem now tells us that the new global optimum will have $\lambda_2 > 0$ and therefore the microorganism will mix EFMs already at these, formerly critical, conditions. □

##### 6.2 The effect of perturbations on points in the flux versus growth rate plane

**Theorem 9.** *Consider the maximisation problem as in Equation (5) at fixed external $\boldsymbol{x}_0^E$ and internal $\boldsymbol{x}_0^I$ metabolite concentrations and assume that we have two cost vectors in mixable position. We consider the situation in which we tighten the first constraint, i.e. we solve the following maximisation problem*

$$\max_{\boldsymbol{x}^I, \boldsymbol{\lambda}} \left\{ \lambda_1 + \lambda_2 \mid \lambda_i \geq 0,\ \boldsymbol{D}(\boldsymbol{x}_0^E, \boldsymbol{x}_0^I) \cdot \boldsymbol{\lambda} \leq \begin{bmatrix} 1-\nu \\ 1 \end{bmatrix} \right\}, \tag{17}$$

*for $\nu \in [0, 1]$. Then the flux through the second EFM versus growth rate ($\mu = \lambda_1 + \lambda_2$) at these concentrations $\boldsymbol{x}_0$ will follow a piecewise linear path to the origin:*

$$(\mu(\nu), \lambda_2(\nu))_{\boldsymbol{x}_0} = \begin{cases} \frac{1}{\det(\boldsymbol{D})} \left( d_1^1 - d_2^1 + d_2^2 - d_1^2 + \nu \left( d_2^1 - d_2^2 \right), d_1^1 - d_2^1 + \nu d_2^1 \right) & \text{if } \nu \leq 1 - \frac{d_1^2}{d_2^2}, \\ \left( \frac{1-\nu}{d_1^2}, \frac{1-\nu}{d_1^2} \right) & \text{if } \nu > 1 - \frac{d_1^2}{d_2^2}. \end{cases} \tag{18}$$

*Proof.* We have two vectors in mixable position, so Lemma 3 states that the flux $\boldsymbol{\lambda}$ is described by the equality $\boldsymbol{D} \cdot \boldsymbol{\lambda} = [1-\nu, 1]^T$. We therefore invert $\boldsymbol{D}$ to get an expression for the $\lambda$'s.

$$\begin{bmatrix} \lambda_1 \\ \lambda_2 \end{bmatrix} = \frac{1}{\det(\boldsymbol{D})} \begin{bmatrix} d_2^2 & -d_1^2 \\ -d_2^1 & d_1^1 \end{bmatrix} \cdot \begin{bmatrix} 1-\nu \\ 1 \end{bmatrix} = \frac{1}{\det(\boldsymbol{D})} \begin{bmatrix} d_2^2 - d_1^2 - \nu d_2^2 \\ -d_2^1 + d_1^1 + \nu d_2^1 \end{bmatrix} \text{ﬄ}$$

Note that we must have nonnegative fluxes $\lambda_i > 0$. This constrains the values of $\nu$ in which a combination of the EFMs will be chosen:

$$0 \leq \lambda_1 = \frac{1}{\det(\boldsymbol{D})} \left( d_2^2 - d_1^2 - \nu d_2^2 \right) \quad \Rightarrow \quad \nu \leq 1 - \frac{d_1^2}{d_2^2}.$$

With $\nu > 1 - \frac{d_1^2}{d_2^2}$ the vectors are not in mixable position anymore, so that the optimal solution will involve only the second EFM and solve the inequality

$$\lambda_2 \begin{bmatrix} d_1^2 \\ d_2^2 \end{bmatrix} \leq \begin{bmatrix} 1-\nu \\ 1 \end{bmatrix} \quad \Rightarrow \quad \lambda_2 = \frac{1-\nu}{d_1^2}.$$

For smaller $\nu$, we find

$$\lambda_2 = \frac{1}{\det(\boldsymbol{D})} \left( d_1^1 - d_2^1 + \nu d_2^1 \right),$$

$$\mu = \lambda_1 + \lambda_2 = \frac{1}{\det(\boldsymbol{D})} \left( d_1^1 - d_2^1 + d_2^2 - d_1^2 + \nu \left( d_2^1 - d_2^2 \right) \right)$$

$\square$

The following theorem is proved in an identical fashion.

**Theorem 10.** *Consider the maximisation problem as in Equation (5) at fixed external $\boldsymbol{x}_0^E$ and internal $\boldsymbol{x}_0^I$ metabolite concentrations and assume that we have two cost vectors in mixable position. We consider the situation in which we tighten the second constraint, i.e. we solve the following maximisation problem*

$$\max_{\boldsymbol{x}^I, \boldsymbol{\lambda}} \left\{ \lambda_1 + \lambda_2 \mid \lambda_i \geq 0,\ \boldsymbol{D}(\boldsymbol{x}_0^E, \boldsymbol{x}_0^I) \cdot \boldsymbol{\lambda} \leq \begin{bmatrix} 1 \\ 1-\nu \end{bmatrix} \right\}, \tag{19}$$

*for $\nu \in [0,1]$. Then the flux through the second EFM versus growth rate ($\mu = \lambda_1 + \lambda_2$) at these concentrations $\boldsymbol{x}_0$ will follow a two-part linear path to the origin:*

$$(\mu(\nu), \lambda_2(\nu))_{\boldsymbol{x}_0} = \begin{cases} \frac{1}{\det(\boldsymbol{D})} \left( d_1^1 - d_2^1 + d_2^2 - d_1^2 + \nu \left( d_1^2 - d_1^1 \right), d_1^1 - d_2^1 - \nu d_1^1 \right) & \text{if } \nu \leq 1 - \frac{d_2^1}{d_1^1} \\ \left( \frac{1-\nu}{d_2^1}, 0 \right) & \text{if } \nu > 1 - \frac{d_2^1}{d_1^1} \end{cases} \tag{20}$$

**Theorem 11.** *Consider the maximisation problem as in Equation (5) at fixed external $\boldsymbol{x}_0^E$ but arbitrary internal $\boldsymbol{x}^I$ metabolite concentrations and assume that we have two cost vectors in mixable position. We consider the situation in which we tighten both constraints, i.e. we solve the following maximisation problem*

$$\max_{\boldsymbol{x}^I, \boldsymbol{\lambda}} \left\{ \lambda_1 + \lambda_2 \mid \lambda_i \geq 0,\ \boldsymbol{D}(\boldsymbol{x}_0^E, \boldsymbol{x}^I) \cdot \boldsymbol{\lambda} \leq \begin{bmatrix} 1-\nu \\ 1-\nu \end{bmatrix} \right\}, \tag{21}$$

*for different values of $\nu \in [0,1]$. Then the flux through the second EFM versus growth rate ($\mu = \lambda_1 + \lambda_2$), at these concentrations $\boldsymbol{x}_0^E$, will follow a straight line to the origin*

$$(\mu(\nu), \lambda_2(\nu))_{\boldsymbol{x}} = \frac{1}{\det(\boldsymbol{D})}((d_1^1 - d_2^1 + d_2^2 - d_1^2)(1-\nu), (d_1^1 - d_2^1)(1-\nu)) \qquad \text{for } 0 \leq \nu \leq 1. \tag{22}$$

*Proof.* Note that we can not just copy the proof of Theorem 9 because this theorem makes a stronger statement. The internal concentrations are not fixed, such that the cost vectors can, in principle, adapt to the experimental perturbations. We will show however, that the optimal solutions at all $\nu$ will have the same internal concentrations.

We have two vectors in mixable position, so Lemma 3 states that the flux $\boldsymbol{\lambda}$ is described by the equality $\boldsymbol{D}(\boldsymbol{x}) \cdot \boldsymbol{\lambda} = [1-\nu, 1]^T$. We therefore invert $\boldsymbol{D}$ again to get an expression for the $\lambda$'s.

$$\begin{bmatrix} \lambda_1 \\ \lambda_2 \end{bmatrix} = \frac{1}{\det(\boldsymbol{D})} \begin{bmatrix} d_2^2 & -d_1^2 \\ -d_2^1 & d_1^1 \end{bmatrix} \cdot \begin{bmatrix} 1-\nu \\ 1-\nu \end{bmatrix} = \frac{1}{\det(\boldsymbol{D})} \begin{bmatrix} (d_2^2 - d_1^2)(1-\nu) \\ (d_1^1 - d_2^1)(1-\nu) \end{bmatrix}, \tag{23}$$

but note that the cost vector components $d_i^j$ are still dependent on the internal metabolite concentrations $\boldsymbol{x}^I$. We therefore want to maximize the sum of fluxes

$$O(\boldsymbol{x}) = \lambda_1 + \lambda_2 = \frac{1}{\det(\boldsymbol{D})}(d_1^1(\boldsymbol{x}) - d_2^1(\boldsymbol{x}) + d_2^2(\boldsymbol{x}) - d_1^2(\boldsymbol{x}))(1-\nu)$$

over all $x$. In the optimum all derivatives of this objective function with respect to $\boldsymbol{x}^I$ must be zero, but it is easy to see that this is independent of the value of $\nu$:

$$\nabla_{\boldsymbol{x}^I} O = (1-\nu)\nabla_{\boldsymbol{x}^I}\left(\frac{1}{\det(\boldsymbol{D})}\left(d_1^1(\boldsymbol{x}) - d_2^1(\boldsymbol{x}) + d_2^2(\boldsymbol{x}) - d_1^2(\boldsymbol{x})\right)\right) = 0.$$

Therefore, the vector of internal concentrations $\boldsymbol{x}_0^I$ that was optimal at $\nu = 0$ will also be the maximum for all other values of $\nu$. The cost vector components will therefore not change and Equation (23) shows us that all fluxes, and therefore also $\mu$, scale with $1-\nu$. □

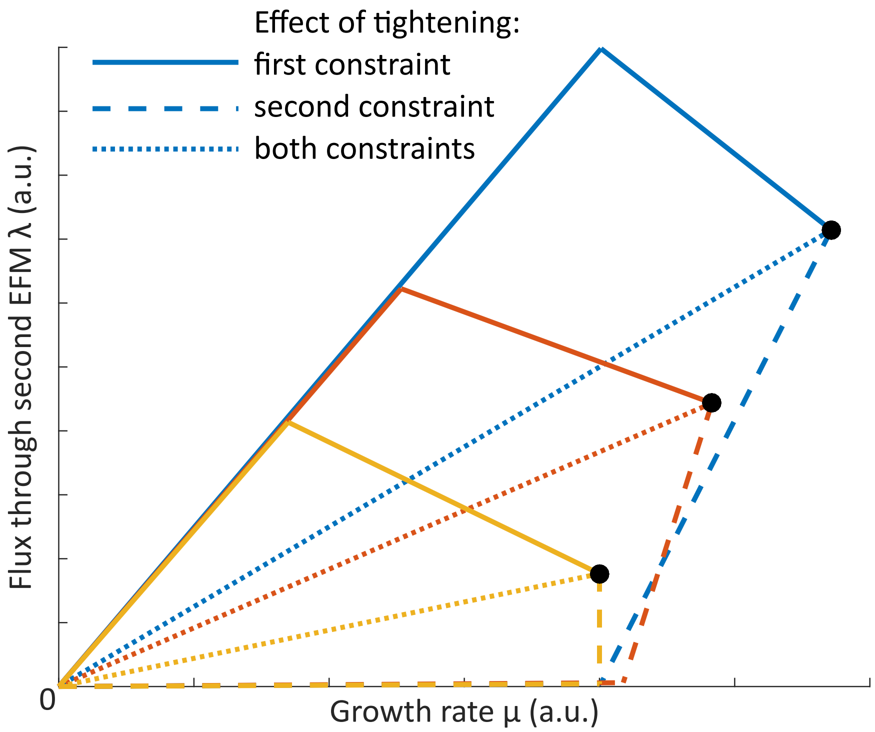


Figure 4: Illustration of the conclusions from Theorems 9, 10 and 11. Tightening different constraints will have different effects on the growth rate $\mu$ and the flux through one of the EFMs, such that we could in principle infer from data which constraint was affected most.

The results from Theorem 9, 10 and 11 are summarized in 4. The following corollary is a biological interpretation of these Theorems, and is summarized in 5.

**Corollary 12.** *Consider a growth maximising wildtype microorganism that shows a metabolic shift from one EFM to two EFMs when the growth rate is increased. The biomass flux through the added EFM can be plotted against the growth rate. A point on this line* $(\mu_{WT}, \lambda_{2,WT})$ *is associated to a vector of metabolite concentrations* $\boldsymbol{x}$. *If these metabolite concentrations are kept fixed while the constrained enzyme pools are experimentally tightened, this will affect the coordinates of the point in the following way.*

a. *tightening the first constraint will increase the ratio* $(\frac{\lambda_2}{\mu})$ *until* 1, *which corresponds to a pure EFM* *b. tightening the second constraint will decrease the ratio $(\frac{\lambda_2}{\mu})$ to zero*

*c. tightening both constraints proportionally will keep the ratio $(\frac{\lambda_2}{\mu})$ constant while the growth rate is decreased to zero*

*Proof.* According to the proof of Theorem 1 a growth maximising microorganism will solve the maximisation problem of Equation (**??**). After the metabolic shift to two EFMs Theorem 6 states that the cost vectors must be in mixable positions. We can then plot the flux through the second EFM $\lambda_2$ versus the growth rate which is proportional to the total biomass flux $\lambda_1 + \lambda_2$. This brings us in the situation in which the Theorems 9, 10 and 11 apply.

Theorem 9 states that tightening the first constraint will move points on the $(\mu, \lambda_2)$-line over the following path

$$(\mu(\nu), \lambda_2(\nu))_{\boldsymbol{x}_0} = \begin{cases} \frac{1}{\det(\boldsymbol{D})}\left(d_1^1 - d_2^1 + d_2^2 - d_1^2 + \nu\left(d_2^1 - d_2^2\right), d_1^1 - d_2^1 + \nu d_2^1\right) & \text{if } \nu \leq 1 - \frac{d_1^2}{d_2^2}, \\ \left(\frac{1-\nu}{d_1^2}, \frac{1-\nu}{d_1^2}\right) & \text{if } \nu > 1 - \frac{d_1^2}{d_2^2}. \end{cases} \tag{24}$$

The ratio $\frac{\lambda_2}{\mu}$ is obviously 1 in the second part of this path. In the first part of the path we see that $\mu$ decreases since $(d_2^1 - d_2^2) < 0$ because the vectors are in mixable position, and $\lambda_2$ increases. Therefore the ratio $\frac{\lambda_2}{\mu}$ increases.

Theorem 10 states that tightening the second constraint will move points on the $(\mu, \lambda_2)$-line over the following path

$$(\mu(\nu), \lambda_2(\nu))_{\boldsymbol{x}_0} = \begin{cases} \frac{1}{\det(\boldsymbol{D})}\left(d_1^1 - d_2^1 + d_2^2 - d_1^2 + \nu\left(d_1^2 - d_1^1\right), d_1^1 - d_2^1 - \nu d_1^1\right) & \text{if } \nu \leq 1 - \frac{d_2^1}{d_1^1}, \\ \left(\frac{1-\nu}{d_2^1}, 0\right) & \text{if } \nu > 1 - \frac{d_2^1}{d_1^1}. \end{cases} \tag{25}$$

The ratio $\frac{\lambda_2}{\mu}$ is 0 in the second part of this path. In the first part of the path the ratio is

$$\frac{\lambda_2}{\mu} = \frac{d_1^1 - d_2^1 - \nu d_1^1}{d_1^1 - d_2^1 + d_2^2 - d_1^2 + \nu\left(d_1^2 - d_1^1\right)},$$

such that

$$\begin{aligned}
\frac{\partial}{\partial \nu}\frac{\lambda_2}{\mu} &= \frac{-\left(d_1^1 - d_2^1 + d_2^2 - d_1^2 + \nu\left(d_1^2 - d_1^1\right)\right) d_1^1 - \left(d_1^1 - d_2^1 - \nu d_1^1\right)\left(d_1^2 - d_1^1\right)}{\left(d_1^1 - d_2^1 + d_2^2 - d_1^2 + \nu\left(d_1^2 - d_1^1\right)\right)^2}, \\
&= \frac{-\left(d_1^1 - d_2^1 + d_2^2 - d_1^2\right) d_1^1 - \left(d_1^1 - d_2^1\right)\left(d_1^2 - d_1^1\right) + \nu d_1^1\left(d_1^2 - d_1^1\right) - \nu\left(d_1^2 - d_1^1\right) d_1^1}{\left(d_1^1 - d_2^1 + d_2^2 - d_1^2 + \nu\left(d_1^2 - d_1^1\right)\right)^2}, \\
&= \frac{-\left(d_1^1 - d_2^1 + d_2^2 - d_1^2\right) d_1^1 - \left(d_1^1 - d_2^1\right)\left(d_1^2 - d_1^1\right)}{\left(d_1^1 - d_2^1 + d_2^2 - d_1^2 + \nu\left(d_1^2 - d_1^1\right)\right)^2}, \\
&= \frac{-d_2^2 d_1^1 + d_2^1 d_1^2}{\left(d_1^1 - d_2^1 + d_2^2 - d_1^2 + \nu\left(d_1^2 - d_1^1\right)\right)^2}, \\
&\leq \frac{-d_2^2 d_1^1 + d_2^2 d_1^1}{\left(d_1^1 - d_2^1 + d_2^2 - d_1^2 + \nu\left(d_1^2 - d_1^1\right)\right)^2}, \\
&= 0.
\end{aligned}$$

Theorem 11 states that tightening both constraints will move points on the $(\mu, \lambda_2)$-line over the following path

$$(\mu(\nu), \lambda_2(\nu))_{\boldsymbol{x}} = \frac{1}{\det(\boldsymbol{D})}((d_1^1 - d_2^1 + d_2^2 - d_1^2)(1-\nu), (d_1^1 - d_2^1)(1-\nu)) \qquad \text{for } 0 \leq \nu \leq 1. \tag{26}$$

The ratio $\frac{\lambda_2}{\mu} = \frac{(d_1^1 - d_2^1)(1-\nu)}{(d_1^1 - d_2^1 + d_2^2 - d_1^2)(1-\nu)} = \frac{d_1^1 - d_2^1}{d_1^1 - d_2^1 + d_2^2 - d_1^2}$ is of course constant. □

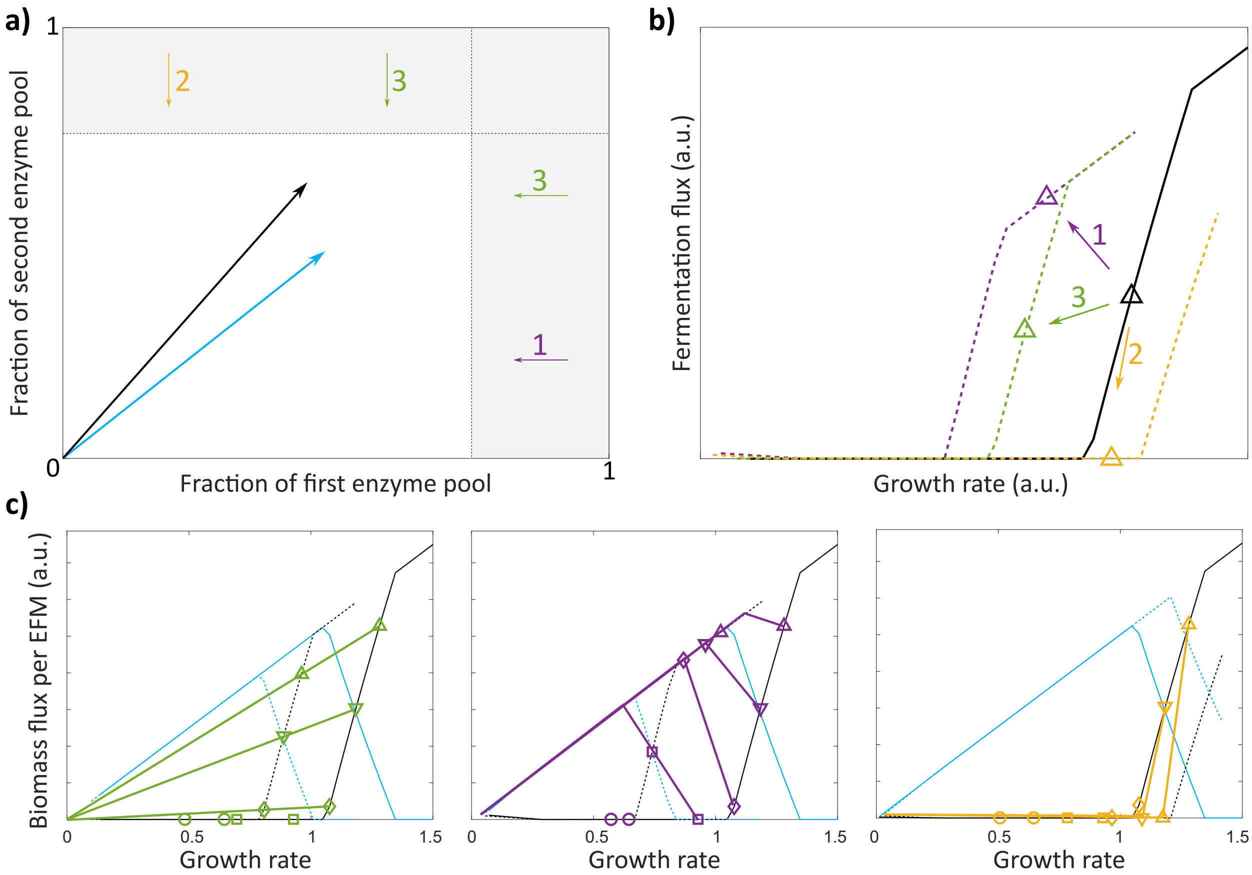


Figure 5: **a)** We used our core model of overflow metabolism to find out the effect of tightening only the first (purple), only the second (orange) or both constraints (green). **b)** The black triangle indicates the optimal growth rate and fermentation flux predicted by the unperturbed model at certain environmental conditions. Tightening of the constraints will force the fermentation flux over growth rate ratio to increase (first constraint), to decrease (second) or to remain constant (both). **c)** We did the same analysis for three different environmental conditions. Solid (green, purple, orange) lines show the effect of tightening (both, the first, the second) constraint(s).

**Remark:**
We have not yet addressed the situation in which one constraint is tightened more than the other, say we have to solve the following maximisation problem

$$\max_{\boldsymbol{x}^I, \boldsymbol{\lambda}} \left\{ \lambda_1 + \lambda_2 \mid \lambda_i \geq 0, \; \boldsymbol{D}(\boldsymbol{x}_0^E, \boldsymbol{x}^I) \cdot \boldsymbol{\lambda} \leq \begin{bmatrix} 1 - \nu_1 \\ 1 - \nu_2 \end{bmatrix} \right\}, \tag{27}$$

with $\nu_1 \neq \nu_2$. To find out what will happen to the ratio $\frac{\lambda_2}{\mu}$ we can just combine two of the situations in Corollary 12. We take the smallest of $\nu_1$ and $\nu_2$, let us say $\nu_1 < \nu_2$. If we solve the maximisation problem $\boldsymbol{D}(\boldsymbol{x}_0^E, \boldsymbol{x}^I) \cdot \boldsymbol{\lambda} \leq \begin{bmatrix} 1-\nu_1 \\ 1-\nu_1 \end{bmatrix}$, the ratio of $\lambda_2$ and $\mu$ will remain constant. Subsequently we tighten the second constraint further until it is at $1-\nu_2$, i.e. $\boldsymbol{D}(\boldsymbol{x}_0^E, \boldsymbol{x}^I) \cdot \boldsymbol{\lambda} \leq \begin{bmatrix} 1-\nu_1 \\ 1-\nu_2 \end{bmatrix}$. This will decrease the ratio. Overall we will thus see an increase of $\frac{\lambda_2}{\mu}$. In this manner all experimental perturbations that tighten the constraints can be analysed.

### 7 The effect on overflow metabolism of a non-lethal dose of chloramphenicol

In the previous section we have focused on experimental perturbations that result in a tightening of a constraint ($\sum_i w_i e_i \leq C$). These perturbations either affect the right side of the constraint, e.g. by taking up a volume in the cytosol when the constraint is a limited cytosolic volume, or by affecting all enzymatic costs on the left side by the same factor, e.g. by the synthesis of an 'average' dummy protein. There are however also perturbations that have a kinetic effect on one specific enzyme, e.g. by reducing the maximal catalytic rate $k_{\text{cat}}$. In this section we show that for chloramphenicol the results in 6 are still valid. Other kinetic inhibitions should be treated on a case by case basis.

Recall the definition of a cost vector

$$d_k^i(\boldsymbol{x}) = \sum_{j=1}^{M} w_j^{(k)} \frac{V_j^i}{f_j(\boldsymbol{x})}. \tag{28}$$

We model the effect of chloramphenicol by a inhibition of the ribosome, say $e_r$, which will result in the multiplication of its activity by a number $\alpha < 1$, i.e. $f_r(\boldsymbol{x}) \to \alpha f_r(\boldsymbol{x})$. Note that we model the ribosome as an enzyme that catalyses the synthesis of proteins. We need to prove a new theorem for this situation.

**Theorem 13.** *Consider the following maximisation problem*

$$\max_{\boldsymbol{x}^I, \boldsymbol{\lambda}} \left\{ \lambda_1 + \lambda_2 \mid \lambda_i \geq 0,\ \boldsymbol{D}(\boldsymbol{x}^E, \boldsymbol{x}^I) \cdot \boldsymbol{\lambda} \leq \begin{bmatrix} 1 \\ 1 \end{bmatrix} \right\}, \tag{29}$$

*where* $\boldsymbol{D} = \begin{bmatrix} \boldsymbol{d}^1(\boldsymbol{x}^E, \boldsymbol{x}^I), & \boldsymbol{d}^2(\boldsymbol{x}^E, \boldsymbol{x}^I) \end{bmatrix}$ *is the cost vector matrix. Assume that the cost vectors are in mixable position, that* $\boldsymbol{x}^E$ *is such that the global maximum has* $\lambda_2 = 0$ *and additionally that at this optimum* $\frac{\mathrm{d}(\lambda_1+\lambda_2)}{\mathrm{d}x^i}|_{x=x_0} = 0$ *for each* $i \in I$. *Then the maximisation problem*

$$\max_{\boldsymbol{x}^I, \boldsymbol{\lambda}} \left\{ \lambda_1 + \lambda_2 \mid \lambda_i \geq 0,\ \tilde{\boldsymbol{D}}(\boldsymbol{x}^E, \boldsymbol{x}^I) \cdot \boldsymbol{\lambda} \leq \begin{bmatrix} 1 \\ 1 \end{bmatrix} \right\}, \tag{30}$$

*with* $\tilde{\boldsymbol{D}} = \begin{bmatrix} d_1^1(\boldsymbol{x}^E, \boldsymbol{x}^I) + w_r^{(1)}\tilde{\rho} & d_1^2(\boldsymbol{x}^E, \boldsymbol{x}^I) + w_r^{(1)}\tilde{\rho} \\ d_2^1(\boldsymbol{x}^E, \boldsymbol{x}^I) & d_2^2(\boldsymbol{x}^E, \boldsymbol{x}^I) \end{bmatrix}$ *has a global maximum at* $\lambda_2 > 0$.

*Proof.* The beginning of the proof of Theorem 7 can be applied here, such that we end up with an equation for the feasible positions of $\boldsymbol{d}^1$:

$$\boldsymbol{d}^1(\boldsymbol{x}) = \boldsymbol{d}^1(\boldsymbol{x}_0) + \left[c\boldsymbol{\delta x} \cdot \left(\boldsymbol{d}^2(\boldsymbol{x}_0) - \boldsymbol{d}^1(\boldsymbol{x}_0)\right)\right] + \left[\boldsymbol{\delta x} \cdot (H(\boldsymbol{d}^1)(\boldsymbol{x}_0) \cdot \boldsymbol{\delta x})\right] + \cdots . \tag{31}$$

In the new maximisation problem of Equation (30) both cost vectors will be shifted. For these new cost vectors we have two options for the maximum of $\lambda_1 + \lambda_2$: $\lambda_2 = 0$ or $\lambda_2 > 0$, again leading to the strategies 1) adjust the metabolite concentrations such that the first cost vectors will be on the diagonal, and 2) take a combination of the new cost vectors to end up on the diagonal. For option 2) we have at least one feasible combination, i.e. using the old metabolite concentrations. This will give an intersection with the diagonal somewhere along the line

$$\boldsymbol{d}^1(\boldsymbol{x}_0)+\begin{bmatrix} w_r^{(1)}\tilde{\rho} \\ 0 \end{bmatrix}+m(\boldsymbol{d}^2(\boldsymbol{x}_0)+\begin{bmatrix} w_r^{(1)}\tilde{\rho} \\ 0 \end{bmatrix}-\boldsymbol{d}^1(\boldsymbol{x}_0)-\begin{bmatrix} w_r^{(1)}\tilde{\rho} \\ 0 \end{bmatrix}) = \boldsymbol{d}^1(\boldsymbol{x}_0)+\begin{bmatrix} w_r^{(1)}\tilde{\rho} \\ 0 \end{bmatrix}+m(\boldsymbol{d}^2(\boldsymbol{x}_0)-\boldsymbol{d}^1(\boldsymbol{x}_0)).$$

The linear map of Equation (12) can now again be applied to both options, yielding

$$\omega\left(\boldsymbol{d}^1(\boldsymbol{x}_0) + \begin{bmatrix} w_r^{(1)}\tilde{\rho} \\ 0 \end{bmatrix} + m(\boldsymbol{d}^2(\boldsymbol{x}_0) - \boldsymbol{d}^1(\boldsymbol{x}_0))\right) = \begin{bmatrix} v \\ w \end{bmatrix} \cdot \begin{bmatrix} d_1^1(\boldsymbol{x}_0) + w_r^{(1)}\tilde{\rho} \\ d_2^1(\boldsymbol{x}_0) \end{bmatrix} = v(d_1^1(\boldsymbol{x}_0)+w_r^{(1)}\tilde{\rho})+wd_2^1(\boldsymbol{x}_0),$$

for the second strategy (mixture), and

$$\begin{aligned} \omega\left(\boldsymbol{d}^1(\boldsymbol{x}) + \begin{bmatrix} w_r^{(1)}\tilde{\rho} \\ 0 \end{bmatrix}\right) &= v\left(d_1^1(\boldsymbol{x}_0) + w_r^{(1)}\tilde{\rho}\right) + wd_2^1(\boldsymbol{x}_0) + \begin{bmatrix} v \\ w \end{bmatrix} \cdot \left[\boldsymbol{\delta x} \cdot (H(\boldsymbol{d}^1)(\boldsymbol{x}_0) \cdot \boldsymbol{\delta x})\right] + \cdots , \\ &= v\left(d_1^1(\boldsymbol{x}_0) + w_r^{(1)}\tilde{\rho}\right) + wd_2^1(\boldsymbol{x}_0) + \left[\boldsymbol{\delta x} \cdot (H(vd_1^1 + wd_2^1)(\boldsymbol{x}_0) \cdot \boldsymbol{\delta x})\right] + \cdots . \end{aligned}$$

for the first strategy (pure solution). We can, as we did in Theorem 7, use the convexity of the cost vectors to see that

$$\omega\left(\boldsymbol{d}^1(\boldsymbol{x}_0) + \begin{bmatrix} w_r^{(1)}\tilde{\rho} \\ 0 \end{bmatrix} + m(\boldsymbol{d}^2(\boldsymbol{x}_0) - \boldsymbol{d}^1(\boldsymbol{x}_0))\right) < \omega\left(\boldsymbol{d}^1(\boldsymbol{x}) + \begin{bmatrix} w_r^{(1)}\tilde{\rho} \\ 0 \end{bmatrix}\right)$$

The intersection of a strategy with the diagonal can be paramaterized by suitable positive scalars $s_{\lambda_2=0}, s_{\lambda_2>0}$, such that $s_{\lambda_2=0}[1,1]^\intercal = \boldsymbol{d}^1(\boldsymbol{x}) + \begin{bmatrix} w_r^{(1)}\tilde{\rho} \\ 0 \end{bmatrix}$ and $s_{\lambda_2>0}[1,1]^\intercal = \boldsymbol{d}^1(\boldsymbol{x}_0) + \begin{bmatrix} w_r^{(1)}\tilde{\rho} \\ 0 \end{bmatrix} + m(\boldsymbol{d}^2(\boldsymbol{x}_0) - \boldsymbol{d}^1(\boldsymbol{x}_0))$. It is now easy to see that

$$\begin{aligned} s_{\lambda_2=0}\,(v+w) = \omega\left(s_{\lambda_2=0}[1,1]^\intercal\right) = \omega\left(\boldsymbol{d}^1(\boldsymbol{x}) + \begin{bmatrix} w_r^{(1)}\tilde{\rho} \\ 0 \end{bmatrix}\right) &> \omega\left(\boldsymbol{d}^1(\boldsymbol{x}_0) + \begin{bmatrix} w_r^{(1)}\tilde{\rho} \\ 0 \end{bmatrix} + m(\boldsymbol{d}^2(\boldsymbol{x}_0) - \boldsymbol{d}^1(\boldsymbol{x}_0))\right) \\ &= \omega\left(s_{\lambda_2>0}[1,1]^\intercal\right) = s_{\lambda_2>0}\,(v+w)\,, \end{aligned}$$

so that $s_{\lambda_2=0} > s_{\lambda_2>0}$. Combined with Lemma 4 this implies that the new maximisation problem will lead to a combination at environmental conditions where a pure strategy was first used.

□

The following corollary and theorem show how the addition of chloramphenicol affects overflow metabolism.

**Corollary 14.** *Assume that we choose the gradual critical environmental conditions (following Definition 3). We experimentally inhibit an enzyme that is part of the pool that inhibits the active EFM most, but which is used in both EFMs in the same amount. Say that the enzyme is inhibited in a linear fashion $f_i(\boldsymbol{x}) \to \alpha f_i(\boldsymbol{x})$. This will force the microorganism to start mixing EFMs at these conditions.*

*Proof.* The multiplicative inhibition will increase the enzymatic cost for this reaction and Equation (28) will change to:

$$\tilde{d}_k^i(\boldsymbol{x}) = \sum_{j \neq r} w_j^{(k)} \frac{V_j^i}{f_j(\boldsymbol{x})} + w_j^{(k)} \frac{V_r^i}{\alpha f_r(\boldsymbol{x})}.$$

Now write $\alpha = 1 - \beta$ with $\beta > 0$, we can rewrite

$$\tilde{d}_k^i(\boldsymbol{x}) = \sum_{j \neq r} w_j^{(k)} \frac{V_j^i}{f_j(\boldsymbol{x})} + w_j^{(k)} \frac{V_r^i}{f_r(\boldsymbol{x})} + w_j^{(k)} \frac{\beta}{1-\beta} \frac{V_r^i}{f_r(\boldsymbol{x})} = d_k^i(\boldsymbol{x}) + w_j^{(k)} \frac{\beta}{1-\beta} \frac{V_r^i}{f_r(\boldsymbol{x})} = d_k^i(\boldsymbol{x}) + w_j^{(k)} \rho \frac{V_r^i}{f_r(\boldsymbol{x})},$$

where $\rho = \frac{\beta}{1-\beta} = \frac{1-\alpha}{\alpha}$. In the case of the ribosome we can make a simplifying assumption. The synthesis of proteins is essential for cell growth and will therefore be present in every EFM that leads to cell synthesis. Moreover, we normalized the cost vector such that it show the costs to produce one unit cell synthesis flux; therefore, the flux through the protein synthesis reaction $V_r$ will be approximately equal in both EFMs, i.e. $V_r^2 \approx V_r^1$. Hence, if we use $\rho \frac{V_r^2}{f_r(\boldsymbol{x})} = \rho \frac{V_r^2}{f_r(\boldsymbol{x})} \approx \tilde{\rho}$ we can write

$$\tilde{d}_k^i(\boldsymbol{x}) = d_k^i(\boldsymbol{x}) + w_j^{(k)} \tilde{\rho}.$$

The costs of both EFMs will thus increase with the same amount in the direction of both constraints. This is exactly the situation in which Theorem 13 applies.

It can be shown that the other conditions necessary for Theorem 13 are also satisfied using the same proof as for Corollary 8 □

We can also prove statements analogous to Corollary 12 in the case of chloramphenicol addition.

**Theorem 15.** *Consider a growth maximising wildtype microorganism that shows a metabolic shift from one EFM to two EFMs when the growth rate is increased. The cell synthesis flux through the added EFM can be plotted against the growth rate. A point on this line $(\mu_{WT}, \lambda_{2,WT})$ is associated to a vector of metabolite concentrations $\boldsymbol{x}$. If, at those metabolite concentrations, an enzyme is inhibited in a linear fashion $f_i(\boldsymbol{x}) \to \alpha f_i(\boldsymbol{x})$ for some $\alpha > 0$, this will affect the coordinates of the point in the following way:*

a. *inhibiting an enzyme in the first constrained pool will increase the ratio $(\frac{\lambda_2}{\mu})$ until 1, which corresponds with the usage of one EFM*

b. *inhibiting an enzyme in the second constrained pool will decrease the ratio $(\frac{\lambda_2}{\mu})$ to zero*

c. *inhibiting an enzyme which forms an equal part of both constraints will keep the ratio $(\frac{\lambda_2}{\mu})$ constant while the growth rate is decreased to zero*

*Proof.* In 6 we explicitly solved the growth rate maximisation in the case of overflow metabolism:

$$\lambda_1(\boldsymbol{x}_0) = \frac{d_2^2(\boldsymbol{x}_0) - d_1^2(\boldsymbol{x}_0)}{\det \boldsymbol{D}}, \quad \lambda_2(\boldsymbol{x}_0) = \frac{d_1^1(\boldsymbol{x}_0) - d_2^1(\boldsymbol{x}_0)}{\det \boldsymbol{D}}.$$

The cost vectors however change as described above (recall the monotonous relation between $\tilde{\rho}$ and the perturbation $1-\alpha$), resulting in the following $\lambda$'s:

$$\lambda_1(\boldsymbol{x}_0) = \frac{d_2^2(\boldsymbol{x}_0) + w_r^{(2)}\tilde{\rho} - d_1^2(\boldsymbol{x}_0) - w_r^{(1)}\tilde{\rho}}{\det \tilde{\boldsymbol{D}}}, \quad \lambda_2(\boldsymbol{x}_0) = \frac{d_1^1(\boldsymbol{x}_0) + w_r^{(1)}\tilde{\rho} - d_2^1(\boldsymbol{x}_0) - w_r^{(2)}\tilde{\rho}}{\det \tilde{\boldsymbol{D}}}.$$

Note that these expressions for $\lambda_i$ are only valid when the cost vectors are in mixable position. The expression for the fraction $\frac{\lambda_2}{\mu}$ becomes

$$\frac{\lambda_2}{\mu} = \frac{\lambda_2}{\lambda_1 + \lambda_2} = \frac{d_1^1 - d_2^1 + w_r^{(1)}\tilde{\rho} - w_r^{(2)}\tilde{\rho}}{d_2^2 - d_2^1 - d_1^2 + d_1^1}. \tag{32}$$

In the regime where the cost vectors are in mixable position, the denominator is positive and the derivatives with respect to $\tilde{\rho}$ can be calculated from Equation (32). Because of the monotonous relation between $\tilde{\rho}$ and the perturbation $1-\alpha$, we indeed see that the fraction $\frac{\lambda_2}{\mu}$ increases, decreases or is stable when the inhibited enzyme is in the first, second or both constrained pools respectively.

Eventually the cost vectors will however not be in mixable position anymore. When the inhibited enzyme is part of the first constrained pool, then $\boldsymbol{d}^2(\boldsymbol{x})$ will become lower-diagonal. In that case $\frac{\lambda_2}{\mu}$ becomes one. The inhibition of an enzyme of the second pool forces $\boldsymbol{d}^1(\boldsymbol{x})$ to become above-diagonal, in which case only $d^1(\boldsymbol{x})$ will be used and $\frac{\lambda_2}{\mu} = 0$. When the enzyme is an equal part of both constrained pools, we can mimic the proof of Theorem 11 to show that the fraction will remain constant. □

### 8 Experimental evidence of a small number of EFMs

We here reinterpret two sets of experimental data that indicate the usage of only a small number of EFMs. Firstly, we show that the substrate uptake fluxes increase proportionally to the growth rate over a large range of environmental conditions for many organisms, Figure **??** in main text. We will explain below that this is a strong indication that there is only 1 active EFM in that range of conditions.

Secondly, Stosch et al [10] performed a variance analysis on measured flux data of *S. cerevisiae* and *P. pastoris*. They could explain 90 percent of the *S. cerevisiae* data by using only 10 EFMs and 99 percent of the *P. pastoris* data with only 6 EFMs. Note that this analysis was performed on a dataset compiled out of different experiments, using several strains and different conditions. That the major part of variance across these conditions can be explained from only 6-10 EFMs provides an upper bound on the number of active EFMs at the separate conditions.

Combining these two analyses, we estimate the number of simultaneously active EFMs to be in the order of 1 to 3.

#### 8.1 Constant growth rate over flux ratios indicate the use of only one EFM

In a metabolic network with a substrate uptake flux $v_1$ and a flux of interest $v_r$ we define the substrate $r$-yield as the ratio $Y^r = v_r/v_1$. Within the $i^{\text{th}}$ EFM, this value $Y^i$ is fixed, since all ratios between flux values are fixed within an EFM.

A combination of EFMs, as in Equation (1), will lead to a yield that combines the yields of the pure EFMs:

$$Y = \frac{\lambda_1 Y^1 + \cdots + \lambda_F Y^F}{\lambda_1 + \cdots + \lambda_F}, \text{ where } \lambda_i \geq 0. \tag{33}$$

The $\lambda_i$ factors are dependent on metabolite concentrations and will therefore vary between different experiments. A cell that uses a combination of EFMs is thus expected to show different yields at different conditions.

We observe a constant ratio of growth rate over substrate uptake fluxes (glucose and oxygen) in a large range of environmental conditions, see data in the main text. The growth rate $\mu$ is proportional to the cell synthesis flux, which is now the flux of interest $v_r$. The most straightforward explanation of the experimental data is that only one EFM is used in this range of conditions (only one nonzero $\lambda_i$ in (1)). Within one EFM, the flux through the EFM can change, but the ratio between fluxes remain constant, leading to the straight line before the critical growth rate.

After the critical growth rate, an additional EFM is clearly used, since the ratio of growth rate over uptake flux changes. Note that the glucose yield decreases rapidly after this critical growth rate, which can be expected because of the onset of overflow metabolism.

An alternative hypothesis is that a combination of EFMs is used in exactly the same proportion in the full range of growth rates below the critical growth rate. This however seems rather unlikely because metabolite concentrations will influence the $\lambda$-factors, thereby changing the relative use of active EFMs which would lead to changed growth rate over flux ratios.

A more probable alternative is the use of a combination in which one EFM is very dominant. Fluctuations in the use of the other EFMs could then become smaller than the experimental noise, hence showing constant yields. A last alternative could be the simultaneous use of a combination of EFMs with exactly the same flux ratios. In that case, a combination would not give rise to a changing yield. We however see that both the glucose and the oxygen yield are very constant. Therefore, both of these yields should be the same in the separate EFMs. These last two alternatives cannot be completely excluded, but will only result in small deviations from our theory.

### 9 Core model of overflow metabolism

The core model of overflow metabolism which was referred to in the main text was made using Matlab (Matlab-files are supplemented). In this model we picked an objective reaction: the reaction towards biomass. The flux through the objective reaction was fixed and we minimized the substrate concentration, which is analogous to the actual selective pressure in a chemostat. The Matlab programs (which are attached to the SI) return the optimal concentrations of all metabolites and enzymes.

The cost vectors that were shown in the main text in **??** were calculated at the optimal metabolite concentrations. We calculated the necessary fractions of the constrained enzyme pools to obtain one unit of enzyme flux, according to the definition of a cost vector given in Equation (**??**). The alternative positions of the cost vectors are obtained by starting from the optimal concentrations and then vary one of the intracellular metabolite concentrations.

#### 9.1 Overflow metabolism

The rate equations used were all of the form $v_i = k_{\text{cat},i} e_i f_i(\boldsymbol{x})$. In the overflow metabolism model we used a transporter ($v_{tr}$) reaction, a respiration ($v_r$) and a fermentation ($v_f$) reaction and a biomass $v_{BM}$ reaction. More detailed results of this model can be found in 6.

A remarkable aspect of this simulation is that the flux through respiration is before the critical growth rate not proportional to the concentration of respiration proteins. This behaviour is found in various experiments [11,12] and cannot be explained by current models that fix enzyme saturations. In fact, the presence of two separate enzyme constraints causes this behaviour: the respiration proteins (which are part of the first constrained pool) are expressed at low growth rates to reduce the cost of the EFM for the transporter pool by reducing product inhibition by keeping internal glucose levels low.

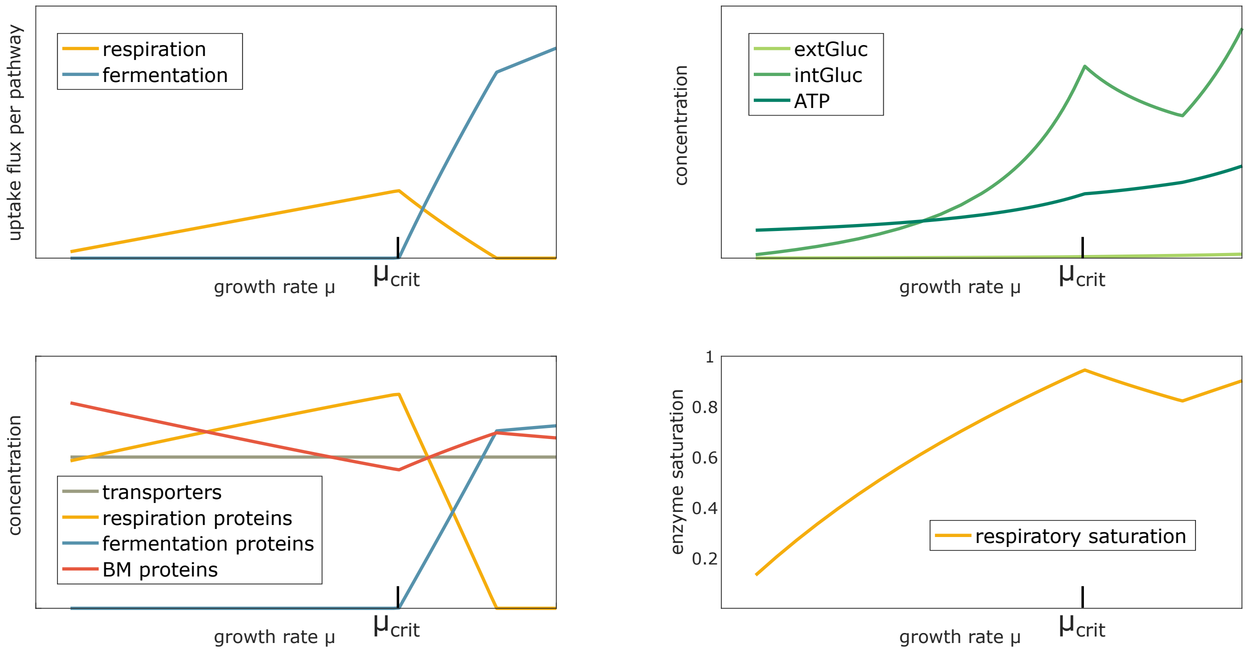


Figure 6: Additional model results from our core model of overflow metabolism. The flux through respiration increases proportionally to the growth rate until the critical growth rate $\mu_{\text{crit}}$, after which respiratory flux decreases and fermentation flux increases. In our model, the membrane constraint is always saturated, indicated by a constant level of transporters. Respiratory proteins do increase slightly with growth rate, but not with the same factor as respiratory flux, indicating an increase in their activity: indeed, internal glucose levels increase up to the critical growth rate.

##### 9.2 Source code

**Code for running kinetic model of overflow metabolism** The Matlab-code used for modeling overflow metabolism is attached in a compressed folder as a supplement. In the compressed folder, we have also added a text-file with instructions.

#### 10 Core model of *L. lactis* switch

The core model of the metabolic switch by *L. lactis* which was referred to in the main text was also made using Matlab (Matlab-files are supplemented). In this model we picked the reaction towards biomass as the objective function again. This objective function is maximized under enzyme constraints (a membrane and a cytosolic constraint), and given glucose and pyruvate concentrations. Here, we make the assumption that the pyruvate concentration is not only determined by glycolysis, but also by other cellular processes. To model this more accurately, these cellular processes should be included, but this is beyond the scope of the paper. This model was merely made to emphasize that protein concentrations can remain constant while pathway usage changes. The Matlab programs (which are attached to the SI) return the optimal concentrations of ATP and all enzymes, and an estimated growth rate.

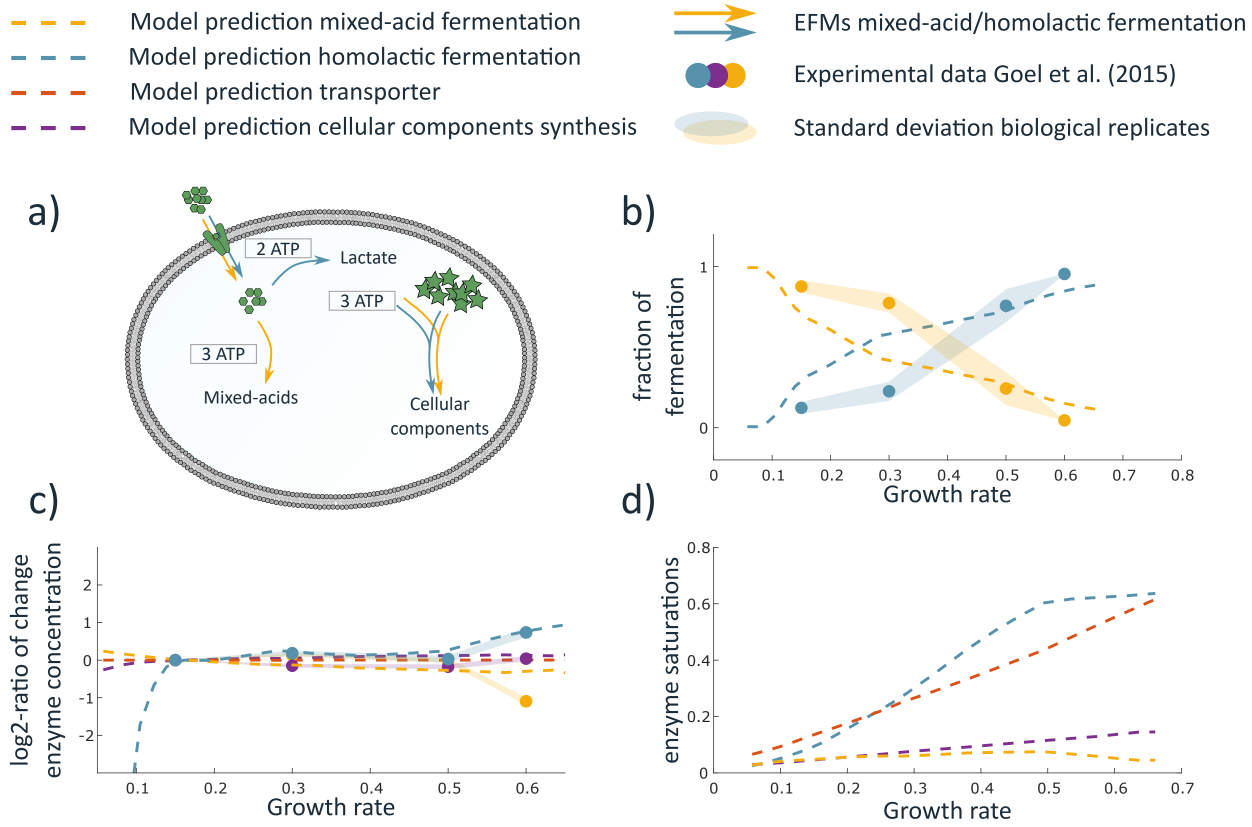

Figure 7: **a)** Core model of energy-limited *L. lactis*-metabolism. ATP can be generated with mixed-acid fermentation (yielding a total of 3 ATP/glucose) and homolactic fermentation (2ATP/glucose). We assume that the cell synthesis reaction is limited only by ATP-concentration. **b-c)** Model predictions and data of the fermentation fluxes and enzyme concentrations. **d)** The change of enzyme saturations as a function of growth rate. The growth rate increases mainly because transporter saturation increases, such that more intracellular metabolite (we will call this pyruvate from now on) becomes available. At first, both mixed-acid and homolactic fermentation proteins become saturated, although saturation of homolactic fermentation goes faster because of the different kinetic properties. Since this makes homolactic fermentation more favourable, this induces a switch from mixed-acid to homolactic fermentation. Then, it becomes important that product inhibition by ATP is less for homolactic fermentation. The higher the fraction of fermentation flux through homolactic fermentation becomes, the more favourable it becomes to have higher ATP-concentrations. High ATP-concentrations lead to a more saturated cell synthesis reaction. As a consequence of this adapting ATP-concentration, saturation of homolactic fermentation is faster than linear with growth rate, while the saturation of mixed-acid fermentation decreases. During this switch, proteins are re-allocated from mixed-acid fermentation to homolactic fermentation. However, because proteins are re-allocated within these pathways too (from the homolactic acid reaction to the cell synthesis reaction, and from the cell synthesis reaction to mixed acid fermentation), enzyme concentrations can still remain constant.

As in our model of overflow metabolism, we used a membrane and a cytosolic protein pool constraint, but *L. lactis* differs from for example *E. coli* in that it takes up amino acids instead of synthesizing them. We therefore hypothesized that in a glucose-limited chemostat, *L. lactis* would be energy-limited rather than carbon-limited. We implemented this in our model by considering a biomass reaction of which the rate was determined solely by the ATP-concentration and the corresponding enzyme concentration. Both mixed-acid fermentation and homolactic fermentation generate ATP, but the amount of product inhibition by ATP differs.

At low growth rates, the membrane constraint is the most dominant constraint, and *L. lactis* will therefore maximize ATP-production no matter the cytosolic protein costs involved. Therefore, only the mixed-acid fermentation pathway is used. At higher growth rates (and thus increasing saturation of the glucose transporter) the cytosolic protein pool becomes limiting too. To use the biomass-producing proteins as efficiently as possible, the ATP-concentration must rise, inducing strong product inhibition on the mixed-acid fermentation pathway. This causes a switch from mixed-acid fermentation to homolactic fermentation, and resources are re-allocated from one pathway to another. As a consequence, one could expect enzyme concentrations in the homolactic pathway to rise. However, since at the same time resources are re-allocated within the pathways, the enzyme concentrations can remain constant, see 7.

##### 10.1 Source code

**Code for running kinetic model of *L. lactis*** The Matlab-code used for the kinetic model of *L. lactis* is attached in a compressed folder as a supplement. In the compressed folder, we have also added a text-file with instructions.

### 11 Finding EFMs that co-consume carbon sources

We showed that the number of active EFMs in a growth-maximising micro-organism is bounded by the number of active enzymatic constraints. Since the number of these constraints seems to be low in many experiments, we asked if it could be optimal to co-consume multiple carbon sources given this small number of constraints. We therefore investigated if there is an EFM that by itself consumes two or even three carbon sources and found EFMs for all combinations that we have tested.

The Python- and Matlab-program that we used to find these EFMs is attached to the SI. It works by first performing a Flux Balance Analysis on a genome-scale metabolic network. We have used the *E. coli* model: `iECSE_1348` [13]. In this FBA, the exchange reactions that are set reversible are either essential (trace elements, oxygen, ammonium) or the carbon sources of interested. This makes sure that the optimal solution of this FBA will use these carbon sources. Subsequently, the inactive reactions in the optimal solution are deleted from the network. The resulting smaller network is loaded into Matlab, where a package is used [14] to enumerate the EFMs. These EFMs are checked for co-consumption of the carbon sources.

Note that we find only *if* there is an EFM that co-consumes the carbon sources of interest, not how many EFMs exist that do this.

#### 11.1 Source code

**Code for finding coconsumption EFMs** The Python and Matlab-code used for finding co-consuming EFMs are attached in a compressed folder as a supplement. In the compressed folder, we have also added a text-file with instructions.

### 12 Coconsumption experiment

#### 12.1 Method

**Strain information**
All experiments were performed with *E. coli* strain MG1655.
**Growth conditions**
The medium employed was the N- C- minimal medium from Gutnick et al. [15], which contains (per liter): $K_2SO_4$ (1 g), $K_2HPO_4$ (13.5 g), $KH_2PO_4$ (4.7 g), $MgSO_4 \cdot 7H_2O$ (0.1 g) and NaCl (2.5 g), supplemented with 20 mM $NH_4Cl$ and thiamine (1mg). After adding saturating amounts of either a single carbon substrate or a combination of multiple (see Table 1), the pH was set to 7.1 using KOH.

Table 1: Carbon mixtures that have been used in combination with the N- C- minimal medium.

| Glucose 0.4 % (w/v) | Mannose 20 mM | Maltose 20 mM | Succinate 15 mM | Xylose 20 mM | Abbr |
|---|---|---|---|---|---|
| x | | | | | G |
| | x | | | | M |
| | | x | | | L |
| | | | x | | S |
| | | | | x | X |
| | | x | x | | SL |
| | x | x | | | ML |
| | | x | | x | XL |
| | | | x | x | XS |
| | x | | x | | SM |
| | | x | x | x | SLX |
| | x | x | x | | SLM |

Cells were seeded from a frozen glycerol stock into 5 ml liquid N-C- minimal medium +glucose and cultured in 30x115 mm conical tubes, shaking with 220 rpm at 37°C. During the subsequent 12 hours they were sequentially diluted into tubes with the desired medium, to achieve exponential growth and removal of undesired carbon. Then, depending on the specific growth rate in each condition, cells of the various cultures were diluted to different densities and 200 $\mu$l of each was transferred to a Greiner 96-well, flat bottom plate. The plate was kept shaking at 37°C and densities were measured at 600 nm using a Spectramax 384 plus (Molecular Devices). Cell densities were chosen such that 8 doublings could take place before growth in the plate could be detected. Every condition was represented by 10 micro-wells (i.e. technical replicates) during each experiment, to make sure enough volume was available for sampling. Samples were taken during growth, filtrated and stored at -20°C for further analysis. This experiment was performed in triplo, meaning that three biological replicates were done on separate days.

**Carbon substrate uptake measurements**

50$\mu$l of undiluted samples were analysed on their mannose, maltose, succinate and xylose content using HPLC (Shimadzo, LC-20AT) at a flow rate of 0.5 mL/min. Calibration samples were made for individual- and triple carbon sources in N-C- minimal medium, to determine the concentrations and validate good separation. Compounds were separated on an ROA-Organic Acid H+ column (Phenomenex, Rezex) and detected using refractive index (Shimadzu, RID-10A) and UV-Vis (Shimadzu, SPD-20A).

Acetate concentrations were measured using an enzyme essay described by Smith et al. [16]. Samples were diluted either 10 or 100 times to stay in the linear range of NADH detection. The essay was conducted in a 96-well, flat bottom plate at 37°C and NADH oxidation was measured at 340 nm using a Spectramax 384 plus (Molecular Devices).

#### 12.2 Data analysis

The three plate-reader experiments resulted in two types of data: OD measurements and HPLC (High Performance Liquid Chromatography) analyses of growth medium samples. The OD measurements were taken every 5 minutes during the full growth experiment and in total 104 growth medium samples were taken at different ODs for all conditions.

The OD measurements were analysed using Matlab. Background OD was subtracted and time windows of at least two hours were selected in which the natural logarithm of the measurements was sufficiently linear: ($R^2 > 0.95$). For these windows the specific growth rate ($\mu = \frac{1}{OD}\frac{\mathrm{d}OD}{\mathrm{d}t}$) was calculated and the maximum is reported below.

The HPLC analyses were normalized using a peak in the chromatogram that corresponded to a constant compound (phosphate) in the medium. Compound concentrations were calculated using a linear calibration curve that was made for all compounds. Since we were interested in the decrease of substrate concentration, rather than in the absolute value of these concentrations, the concentrations were normalized such that the $t_0$-concentration is equal to the intended initial concentration for that compound, thereby correcting for small pipetting errors.

##### 12.3 Results

The calculated growth rates for all 12 conditions are summarized in 8. Data can be found in file: `SI_growth_rates.txt`. In almost all cases the growth rate increases or remains equal when an extra compound is added: only the combination of mannose and maltose leads to a lower growth rate than on maltose alone. The addition of succinate to the medium always leads to an increase in growth rate.

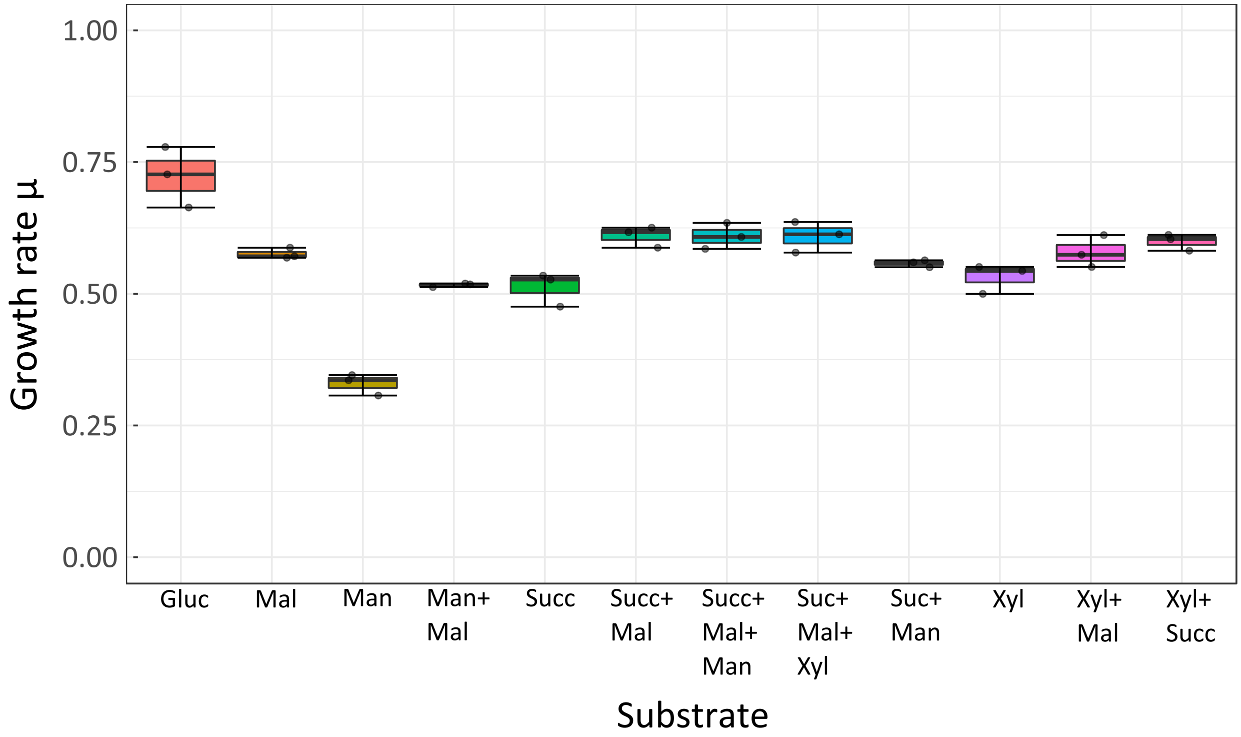


Figure 8: Growth rates were measured for *E. coli* on media with combinations of glucose, maltose, mannose, succinate or xylose. Data from three biological replicates are shown. For each of these replicates, the growth rates from five technical replicates were averaged.

We determined for all conditions the mean specific uptake rate for all the compounds. The resulting data is shown in Table 2 and the standard errors of the means are shown in Table 3. This data can be found in `SI_q_S_comp_cond.xlsx`.

Table 2: Mean specific uptake rate for the 12 conditions and 4 compounds in units: $\frac{\text{mmol}}{\text{OD}_{600} \cdot \text{hour}}$. The letters that indicate the conditions denote the available carbon sources in the medium: S=Succinate, L=maLtose, M=Mannose, X=Xylose, G=Glucose. The uptake rates were estimated from the fitted lines in 9.

| | S | SL | M | ML | L | XL | G | XS | X | SLX | SM | SLM |
|---|---|---|---|---|---|---|---|---|---|---|---|---|
| Succinate | 80.5 | 51.0 | 0.0 | 0.0 | 0.0 | 0.0 | 0.0 | 70.0 | 0.0 | 46.9 | 57.0 | 51.9 |
| Maltose | 0.0 | 59.2 | 0.0 | 32.5 | 71.5 | 42.0 | 0.0 | 0.0 | 0.0 | 35.8 | 0.0 | 30.2 |
| Xylose | 0.0 | 0.0 | 0.0 | 0.0 | 0.0 | 31.3 | 0.0 | 58.0 | 64.8 | 26.9 | 0.0 | 0.0 |
| Mannose | 0.0 | 0.0 | 33.1 | 27.8 | 0.0 | 0.0 | 0.0 | 0.0 | 0.0 | 0.0 | 19.0 | 25.3 |

Table 3: Standard errors of the mean of specific uptake rate for the 12 conditions and 4 compounds in units: $\frac{\text{mmol}}{\text{OD}_{600} \cdot \text{hour}}$. The letters that indicate the conditions denote the available carbon sources in the medium: S=Succinate, L=maLtose, M=Mannose, X=Xylose, G=Glucose. Standard errors of the mean were calculated from the three biological replicates.

| | S | SL | M | ML | L | XL | G | XS | X | SLX | SM | SLM |
|---|---|---|---|---|---|---|---|---|---|---|---|---|
| Succinate | 8.6 | 4.1 | 0.0 | 0.0 | 0.0 | 0.0 | 0.0 | 10.1 | 0.0 | 6.6 | 9.5 | 7.0 |
| Maltose | 0.0 | 2.1 | 0.0 | 1.8 | 4.0 | 4.6 | 0.0 | 0.0 | 0.0 | 2.1 | 0.0 | 2.1 |
| Xylose | 0.0 | 0.0 | 0.0 | 0.0 | 0.0 | 1.8 | 0.0 | 2.9 | 1.9 | 2.2 | 0.0 | 0.0 |
| Mannose | 0.0 | 0.0 | 1.0 | 2.4 | 0.0 | 0.0 | 0.0 | 0.0 | 0.0 | 0.0 | 1.4 | 3.4 |

In Figure 9 we plot, for all conditions except for glucose, the relation between the optical density of cells in the sample and the concentrations of the compounds succinate, maltose, mannose and xylose. The dataset can be found in the file SI_OD_conc_per_cond.xlsx.

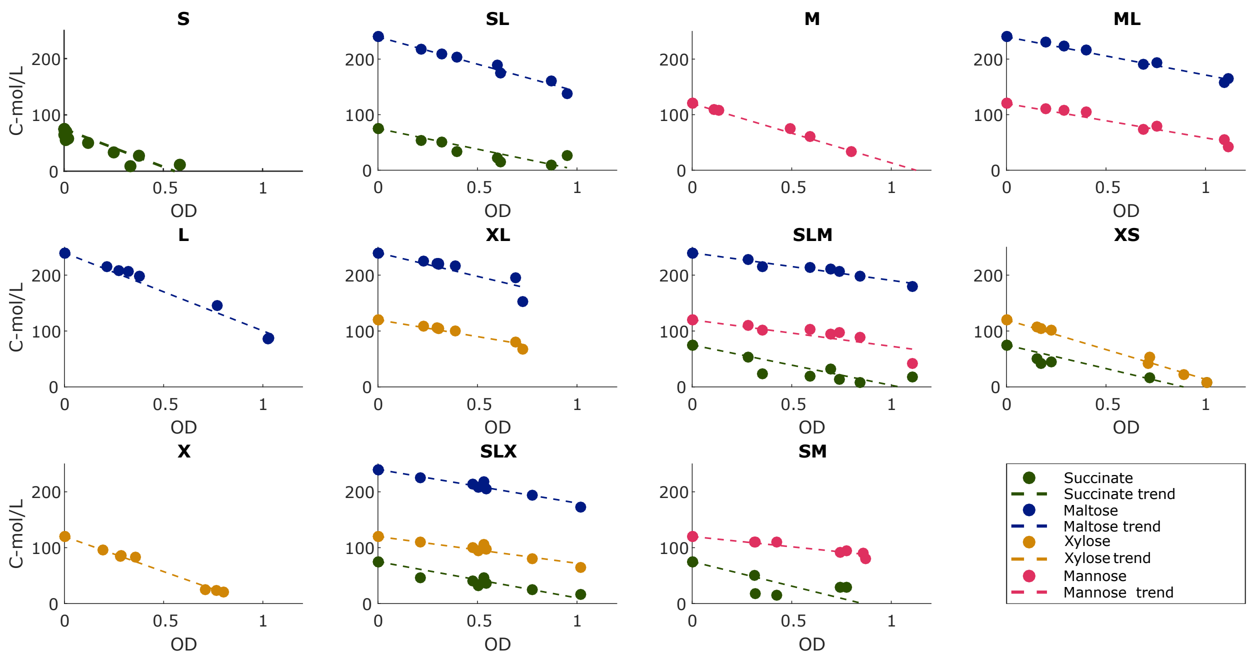

Figure 9: During the growth experiments, the concentration of carbon sources was measured. The letters that indicate the conditions denote the available carbon sources in the medium: S=Succinate, L=maLtose, M=Mannose, X=Xylose, G=Glucose. We here show the decrease in these concentrations as the OD (Optical Density) of the culture increases. Data from three biological replicates was normalized for initial concentration and then shown together. The best linear approximation was calculated and shown by a dashed line.

##### 12.4 Additional datasets and source code for data analysis

**Source Code for Data Analysis Coconsumption Experiment** All raw data and the Matlab-code used for data analysis can be found in the compressed folder attached to the supplements.

**Dataset 1.** `SI_growth_rates.txt` Estimated growth rates from separate biological replicates.

**Dataset 2.** `SI_OD_conc_per_cond.xlsx` For all different growth media, we include an excell-sheet. Shown are the measured concentrations of carbon sources (normalized for initial concentration), with the corresponding Optical Density (OD). The letters that indicate the conditions denote the available carbon sources in the medium: S=Succinate, L=maLtose, M=Mannose, X=Xylose, G=Glucose.

**Dataset 3.** `SI_q_S_comp_cond.xlsx` Shown are the estimated uptake rates (mean and standard deviation) of different carbon sources (normalized for initial concentration) on the different growth media. The letters that indicate the conditions denote the available carbon sources in the medium: S=Succinate, L=maLtose, M=Mannose, X=Xylose, G=Glucose.
